# The evolution of sexual dimorphism in the weapon system of leaf-footed bugs (Hemiptera: Coreoidea)

**DOI:** 10.64898/2026.11.17.767353

**Authors:** Michael Forthman, Rebecca T. Kimball, Christine W. Miller

## Abstract

The natural world contains an extraordinary diversity of morphologically elaborate traits, yet the evolutionary processes underlying the emergence and diversification of sexual dimorphism in these structures are not fully resolved. Some of the most extravagant of these traits are the mechanical weapons of sexual selection. They include such noteworthy structures as the tusks of narwals and the mandibles of stag beetles. Such traits are notoriously sexually dimorphic – often elaborated in one sex while absent or reduced in the other sex. The mechanisms by which sexual dimorphism evolves in such traits is only just beginning to be understood. Existing macroevolutionary studies have examined simple structures or have simplified elaborate and complex weapons using binary coding or metrics, limiting the insights that can be gained. Here, we investigated the macroevolutionary dynamics of sexual dimorphism across 14 traits in bodies and complex, elaborate weaponry of leaf-footed bugs (Hemiptera: Coreoidea) using phylogenomic data and quantitative and qualitative trait analyses. Phylogenetic generalized linear mixed models across 27 species showed that females are generally larger-bodied, whereas males exhibit greater exaggeration in most putative weapon traits, often with steeper allometric scaling. Ancestral state estimation across 154 species revealed that sexual dimorphism is evolutionarily labile and heterogeneous across traits. Contrary to expectations from classical models, sexual dimorphism most commonly arose when new traits originated in both sexes already in a dimorphic state, rather than through subsequent divergence from ancestrally shared, monomorphic traits or as a sex-limited trait. At the level of entire weapon system, sexual dimorphism varied widely across lineages, clustered phylogenetically, and reached extreme levels through coordinated changes across multiple traits. By showing that complex weapon systems diversify through semi-independent changes and that new traits often arise already dimorphic, our study highlights the need for further studies on the developmental mechanisms of sexual dimorphism, how weapon traits covary within and between sexes, and the relative roles of different selective pressures in shaping weapon systems.

## Introduction

From the distinct horns of male and female pronghorn antelope to the presence and absence of exaggerated claws of fiddler crabs, phenotypic differences between males and females are common across taxa. Such differences can be surprising given that males and females of the same species share much of the same genome, raising fundamental questions of how sexual differences evolve. One important driver behind the evolution of sexual dimorphism is differences in the degree and form of intrasexual competition. When intrasexual competition occurs over mating opportunities, it can lead to the evolution and exaggeration of sexually selected weapons (Emlen 2008). Sexually selected weapons, where they exist, can exhibit enormous differences across the sexes.

The *mechanical weapons* of sexual selection (see Glossary)—including enlarged horns, claws, antlers, tusks, and spurs—are functional traits that must be integrated within the overall phenotype so that they can effectively deliver and withstand force during physical conflict. As a result, they are typically part of multicomponent *weapon systems* involving modifications across the body, including muscular and physiological systems (Miller et al. 2026). Further, many have multiple external *components*, with each component often varying in one or more *properties*, such as size, shape, number, and anatomical location. In many cases, both sexes possess many of the same components, though with different properties. Males often, though not always, have the more elaborate versions of these components and use them to drive away other males and monopolize potential mates. The female versions of these components, even if reduced, are typically not without function; yet, their use is often understudied in many taxa.

Mechanical weapons are rarely shaped by combat alone. Many occur on body regions that also function in locomotion, signaling, mating, defense, or other behaviors. Thus, not every component of a weapon system will necessarily function as a weapon in every species, context, or sex. Some components may directly contact rivals during combat, whereas others may support weapon performance receiving or deflecting force, protecting vulnerable regions, or signaling during competitive interactions. Throughout this study, we use “weapon system” in this broad sense: a suite of morphologically elaborated traits associated with structures known to function or hypothesized to function in combat in at least some species, while recognizing that individual components may not be weaponized in all species or in both sexes.

Sex differences in the function of these integrated trait systems means that sexual dimorphism may arise not only through gains or losses of weapon components, but also through divergence in the component properties across the sexes. Despite this complexity, most macroevolutionary studies have characterized relatively few mechanical components and/or properties (Table 1). Additionally, many studies have represented component properties with simplified descriptors such as presence/absence or composite characters (Table 1). Consequently, these limitations have restricted our ability to understand how weapon systems diversify and how sexual dimorphism evolves over evolutionary time. Studies are needed that take an expanded view of mechanical weapons, examining how multiple components of weapon systems arise and come together over evolutionary time, as well as how this process differs in males and females. Such work has the potential to provide valuable insights into the routes by which sexual dimorphism arises in functional traits.

**Table 1.** Summary of macroevolutionary studies of sexual dimorphism in mechanical weapon systems. Components and properties summarize the scope of weapon morphology evaluated in each study. Coding scheme indicates whether weapon components and/or sexual dimorphism were treated as binary, multistate, continuous, or metric characters. Inferred pathways summarize the evolutionary transitions that can be inferred from the published analyses, including transitions inferred here from published sex-specific reconstructions rather than only those emphasized by the original authors. Additional details, including full details on coding schemes, main findings, limits on evolutionary inference, and transition counts, are provided in Table S1.

| Study | Weapon system | Components | Properties | Coding scheme | Inferred evolutionary pathways |
|---|---|---|---|---|---|
| (Calamari 2016) — Ruminants (Ruminantia) | Cranial appendages | **1:** Cranial appendage | **5:** Presence, orientation, appendage number, curvature, surface texture | Binary | Sex-limited properties to shared sexually monomorphic, shared sexually dimorphic properties, and absence in both sexes; and shared sexually dimorphic to shared sexually monomorphic properties. |
| (Kimball et al. 2011) — Pheasants and partridges (Phasianidae) | Spurs, wings, and body | **3:** Spurs, wing, body | **3:** Presence, number, size; not all properties coded for each component | Binary and metric | Specific pathways cannot be inferred from the published figures or results. |
| (Van Kleeck-Hann and Wiens 2023) — Chameleons (Chamaeleonidae) | Head, trunk, and tail horns, crests, casques, and spines | **11:** Casque, rostral appendage, supraorbital spines, occipital lobe, dorsal crest and spines, gular crest and spines, ventral spines, tail crest and spines | **1:** Presence | Binary and metric | Absence of components in both sexes to sex-limited presence or presence in both sexes; sex-limited presence to absence in both sexes or presence in both sexes; and presence in both sexes to absence in both sexes or sex-limited. |
| (Santana et al. 2025) — Frogs (Leptodactylinae) | Male forelimb and chest spines | **3:** Thumb spines, hypertrophied forearms, chest spines | **1:** Presence | Multistate | Absence of components in both sexes to sex-limited components and sex-limited to absence of components. |
| (Anker et al. 2006) — Snapping shrimp (Alpheidae) | Snapping claw and orbital hood | **23:** Orbital hood, cheliped (major and/or minor), palm, adhesive plaques, 6 palm depressions or excavations, merus depressions, dactylus, carpal setae, coxal process, finger cutting-margin complex, major chela plunger setation, finger processes, carpus, ischium processes, finger balaeniceps, chela processes, merus processes, finger tips | **8:** Presence, development, size, asymmetry, shape, location, configuration, texture; not all properties considered for each component | Binary and multistate | Absence of cheliped enlargement in both sexes to enlarged in both sexes or in males only; from enlarged in both sexes to absent in both sexes or enlarged in males only. |
| (Buzatto et al. 2014) — Harvestmen (Opiliones) | Fourth leg | **2:** Coxal apophysis, femur | **1:** Size | Binary | Shared sexually monomorphic to shared sexually dimorphic components and shared sexually dimorphic to shared sexually monomorphic components. |
| (Machado et al. 2021) — Harvestmen (Opiliones) | Fourth leg | **1:** Femur | **1:** Size | Continuous | Because the only component investigated was treated as a continuous character, no specific evolutionary pathways can be readily inferred. |
| (Miller 2003) — Diving beetles (Dytiscidae) | Male protarsal suction cups, female dorsal cuticle | **2:** Male protarsal suction cups, female modified dorsal cuticle | **1:** Presence | Binary | Absence of components in both sexes to sex-limited components and sex-limited to absence of components. |
| (Bergsten and Miller 2007) — Diving beetles (Dytiscidae: *Acilius*) | Male protarsal suction cups, female elytra | **5:** Four types of male protarsal suction cups, female elytra | **2:** Size, presence; not all properties considered for each component | Binary, metric, and continuous | Absence of setose furrows in both sexes to sex-limited spines and sex-limited to absence of setose furrows. |
| (Hosoya and Araya 2005) — Stag beetles (Lucanidae) | Mandibles | **1:** Mandibles | **1:** Shape | Multistate | At least two gains of increased sexual dimorphism. |
| (Emlen et al. 2005a) — Dung beetles (Scarabaeidae: *Onthophagus*) | Head and thoracic horns | **5:** Physical locations treated as separate horn types | **1:** Presence | Binary | Absence of horns in both sexes to shared sexually monomorphic, shared sexually dimorphic, and sex-limited horns; sex-limited horns to absence in both sexes, shared sexually dimorphic, and shared sexually monomorphic horns; and shared sexually monomorphic to shared sexually dimorphic horns. |
| (Emlen et al. 2005b) — Dung beetles (Scarabaeidae: *Onthophagus*) | Head and thoracic horns | **5:** Physical locations treated as separate horn types | **1:** Presence | Binary | Absence of horns in both sexes to shared sexually monomorphic, shared sexually dimorphic, and sex-limited horns; sex-limited horns to absence in both sexes, shared sexually dimorphic, and shared sexually monomorphic horns; shared sexually monomorphic to sex-limited horns; and shared sexually dimorphic to sex-limited horns. |
| (Puniamoorthy et al. 2008) — Sepsid flies (Sepsidae) | Foreleg | **1:** Spines | **1:** Presence | Binary | Absence of spines in both sexes to sex-limited spines and sex-limited to absence of spines. |
| (Rohner and Blanckenhorn 2018)— Black scavenger flies (Sepsidae: *Sepsis*) | Foreleg | **2:** Forefemur, foretibia | **1:** Size | Continuous | Because the only component investigated was treated as a continuous character, no specific evolutionary pathways can be readily inferred. |
| (Andersen 1997) — Water striders (Gerridae) | Male legs, male and female genitalia | **3:** Male clasping legs, male clasping genitalia; female anticlasping genitalia | **3:** Presence, size, shape; not all properties considered for each component | Binary | Absence of components in both sexes to sex-limited presence and sex-limited presence to absence in both sexes. |
| (Crumière et al. 2019) — Riffle bugs (Vellidae: *Rhagovelia*) | Male fore and hind legs, female pronotum and abdomen | **10:** Male foreleg, including sex combs, three femoral spine types, hind tibia spines and curvature, hind trochanter spines; female pronotal projection and abdomen | **3:** Presence, size, shape; not all properties considered for each component | Binary and metric | Absence of components in both sexes to sex-limited presence and sex-limited presence to absence in both sexes. |
| (Armisén et al. 2022) — Semi-aquatic bugs (Gerromorpha) | Hind leg | **1:** Hind leg | **1:** Modification | Multistate | Absence of hind leg modification in both sexes to male-limited modification or to modification in both sexes; modification in both sexes to male-limited modification or to absence in both sexes. |
| (Roth et al. 2024) — Bedbugs (Cimicidae) | Male intromittent organ, female paragenitalia/spermalege | **2:** Male paramere, female spermalege | **5:** Size, shape, three types of locations; not all properties considered for each component | Binary and multistate | All components evaluated were sex-limited in expression and always present in the taxa sampled. As such, no specific evolutionary pathways can be inferred. |
| (Tatarnic and Cassis 2010) — Plant bugs (Miridae: *Coridromius*) | Male intromittent organ, female paragenitalia/spermalege and associated structures | **5:** Male paramere; female dorsal laterotergites, ectospermalege, hemelytron, metepimeron | **4:** Presence, form, size, shape; not all properties considered for each component | Metric | Because a complexity index was analyzed no specific evolutionary pathways can be readily inferred. |

### Glossary of key terms used in this study

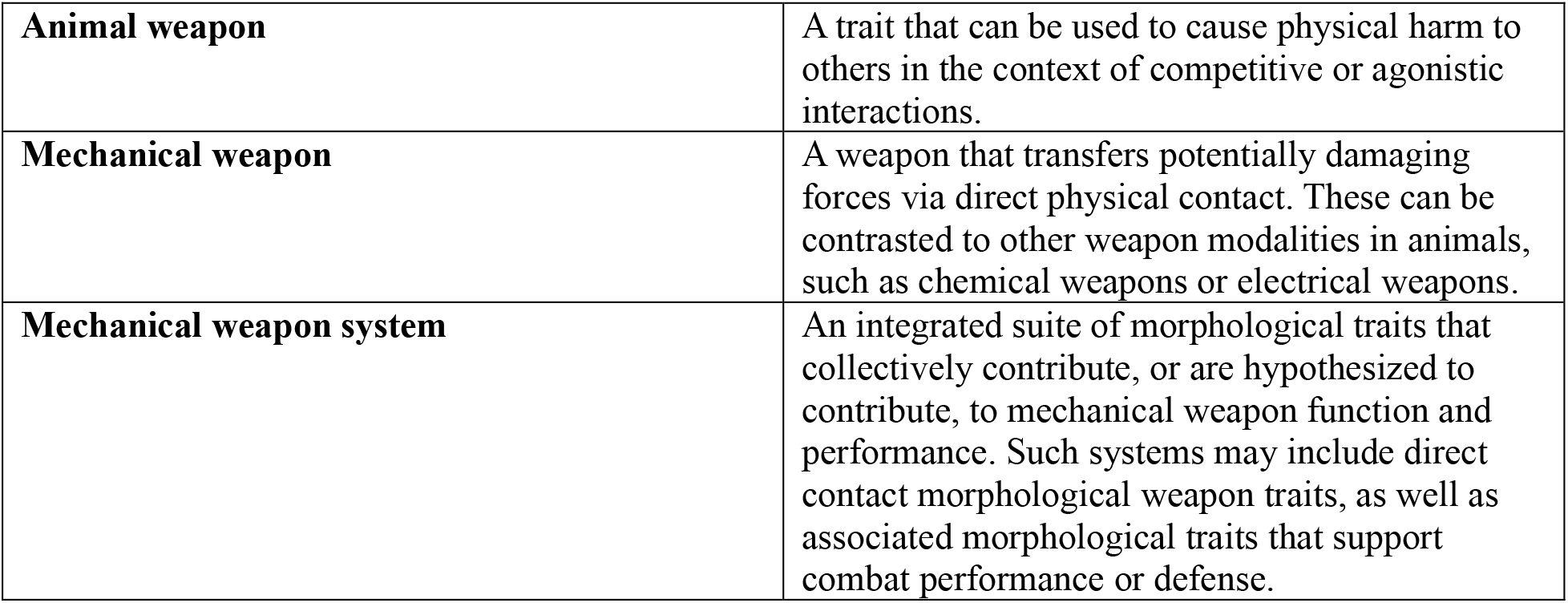
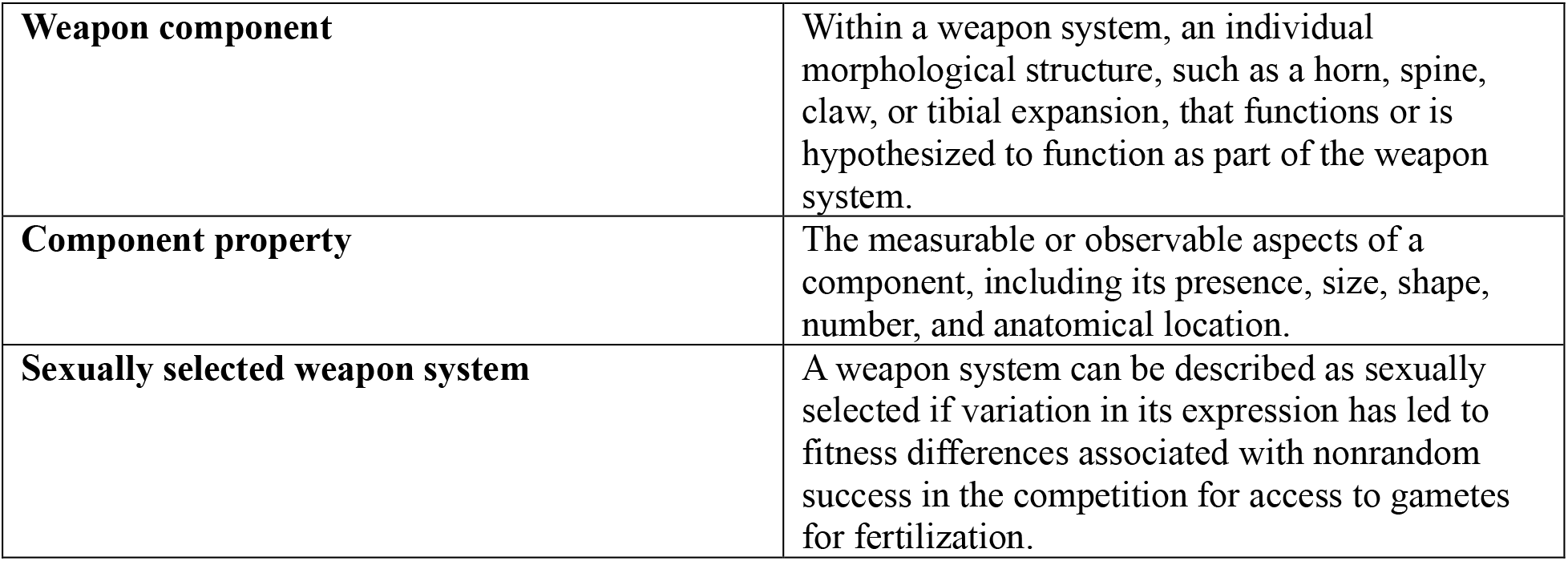

|  |  |
|---|---|
| **Animal weapon** | A trait that can be used to cause physical harm to others in the context of competitive or agonistic interactions. |
| **Mechanical weapon** | A weapon that transfers potentially damaging forces via direct physical contact. These can be contrasted to other weapon modalities in animals, such as chemical weapons or electrical weapons. |
| **Mechanical weapon system** | An integrated suite of morphological traits that collectively contribute, or are hypothesized to contribute, to mechanical weapon function and performance. Such systems may include direct contact morphological weapon traits, as well as associated morphological traits that support combat performance or defense. |
| **Weapon component** | Within a weapon system, an individual morphological structure, such as a horn, spine, claw, or tibial expansion, that functions or is hypothesized to function as part of the weapon system. |
| **Component property** | The measurable or observable aspects of a component, including its presence, size, shape, number, and anatomical location. |
| **Sexually selected weapon system** | A weapon system can be described as sexually selected if variation in its expression has led to fitness differences associated with nonrandom success in the competition for access to gametes for fertilization. |

In this study, we ask two overarching questions about the evolution of sexual dimorphism. <u>First,</u> how does sexual dimorphism in *animal weapons* arise (Figure 1)? Do weapons (a) first show up equally in both sexes, with one sex then losing or reducing expression? Can weapons instead (b) originate in just one sex, or even (c) originate in both sexes, but in a sexually dimorphic form? Macroevolutionary studies across diverse taxa indicate that sexual dimorphism in weapon components is evolutionarily labile, with repeated gains, losses, and transitions between sex-limited and shared expression (Table 1). Further evidence has shown that sexually dimorphic components can arise through routes (a) and (b), while (c) has been inferred less often (Table 1). While these studies have many merits, many have analyzed sexual dimorphism of mechanical weapons as a binary (e.g., “dimorphic” vs. “monomorphic”) or as a metric or continuous (e.g., a score or ratio) character, often limiting the ability to distinguish among specific pathways by which sex differences evolve (Table 1). Thus, studies additional studies are needed that explicitly capture sex differences across multiple weapon components and properties to determine how weapon systems assemble over evolutionary time and how their diversity becomes sexually dimorphic.

**Figure 1.**
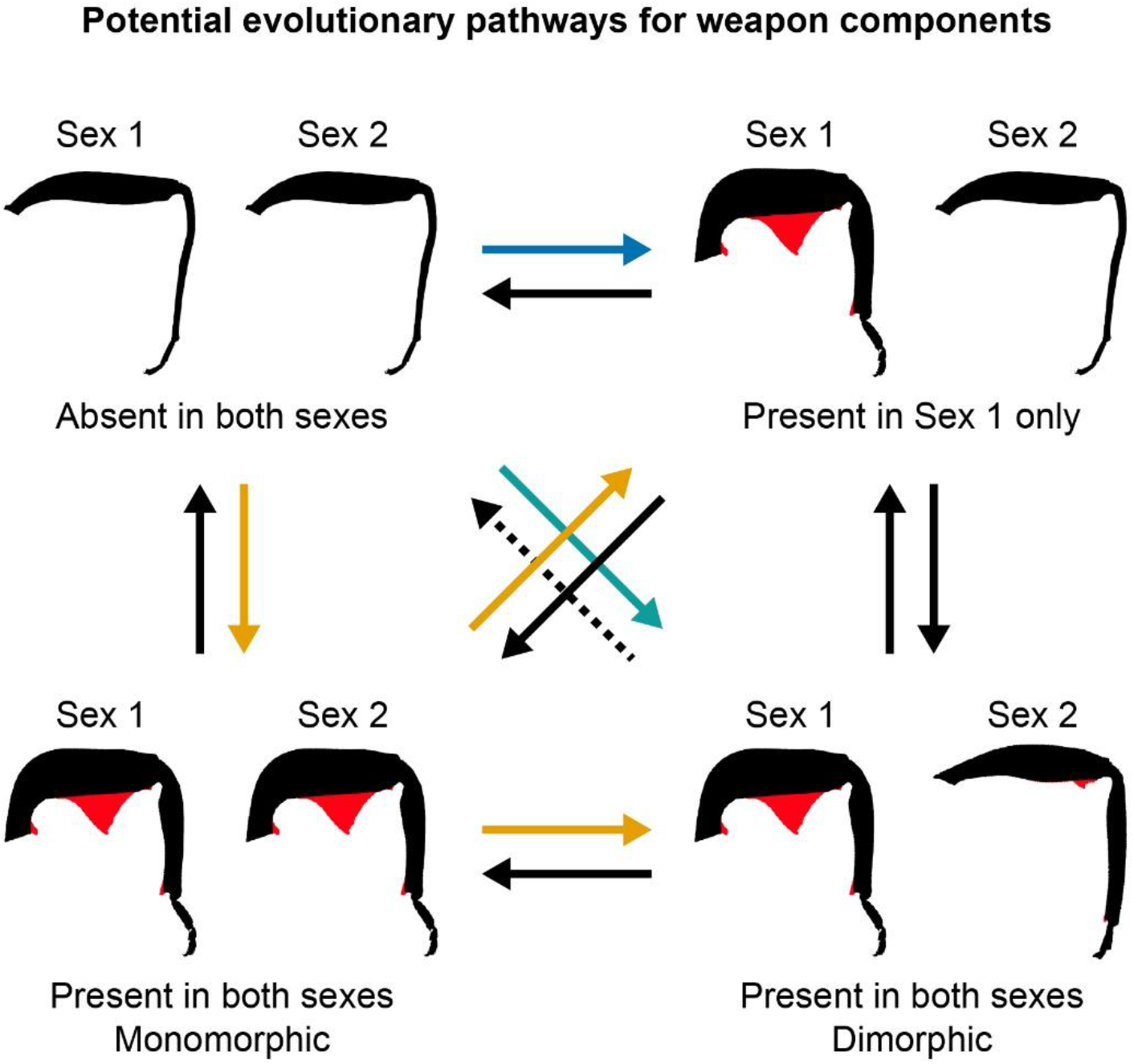
Conceptual pathways for the evolution of sexual dimorphism in mechanical weapons. Two primary evolutionary routes based on classical and quantitative genetic models by which sexual dimorphism may arise are shown in orange and blue arrows (Darwin 1871; Wallace 1889; Fisher 1930; Lande 1980; Rice 1984). In route (a) (orange arrows), a weapon component initially evolves in both sexes with a monomorphic expression but is subsequently lost, reduced, or otherwise modified in one sex, resulting in sexual dimorphism. In route (b) (blue arrow), a novel component originates with sex-limited expression. An additional potential route, (c) (teal arrow) involves the origin of a component in both sexes already in a sexually dimorphic form. Additional transitions (black arrows) may include traits component gains or losses in either sex, transitions between sex-limited and jointly expressed states, and other shifts between sexually monomorphic and dimorphic expression. Sexual dimorphism may involve differences in, for example, the presence, size, number, and/or anatomical location of components. Solid arrows represent evolutionary transitions supported by macroevolutionary studies of mechanical weapons, whereas dashed arrows indicate potential transitions not yet supported in such studies. Silhouettes of hind legs lacking weapon components are schematic and do not represent any particular species. Silhouettes with highlighted components (red) are based on the leaf-footed bug *Anoplocnemis curvipes* (Fabricius 1781), with the lower right panel illustrating accurate sex-specific morphology of this species (Sex 1= male; Sex 2 = female; not drawn to scale).

A <u>second</u> overarching question is how the overall magnitude of sexual dimorphism in weapon systems evolves. While individual weapon components are known to originate, diversify, and be lost repeatedly over evolutionary time (Table 1), it remains unclear whether the degree of sexual dimorphism across entire weapon systems tends to increase, decrease, or fluctuate across lineages. Multicomponent mechanical weapon systems are ideal for addressing this question because they often include discrete, externally visible components and properties that are straightforward to measure and score across species and along which sexual dimorphism can increase, decrease, or shift in form. In these complex systems, sexual dimorphism in the overall system might evolve incrementally—such as through the stepwise addition or exaggeration of individual components over time—, a pattern observed in the chameleon weapon system (Van Kleeck-Hann and Wiens 2023). However, multiple components within a weapon system can be evolutionarily correlated and selected to function as an integrated unit (Kimball et al. 2011; Anderson et al. 2014; Miller et al. 2024, 2026), and sexual dimorphism may evolve more abruptly if selection on a component also produces simultaneous changes across other correlated components or the entire system. Despite extensive work on the evolutionary origins and distribution of individual weapon components, we still lack a clear understanding of how the overall magnitude of sexually dimorphism varies across weapon systems, how this variation is distributed phylogenetically, and whether lineages show repeated escalation or reduction of sexual dimorphism over time.

Leaf-footed bugs and allies (Hemiptera: Coreoidea; hereafter referred to as leaf-footed bugs) are an exceptional invertebrate superfamily for investigating the evolution of sexual dimorphism in multicomponent weapon systems. In many species, males and females express elaborated traits on the hind legs, including robust spines, serrated surfaces, and leaf-like expansions, with additional elaborations sometimes occurring on the thorax or abdomen (Emlen 2008; Miller et al. 2024). We follow (Miller et al. 2024) in treating these traits as components of a weapon system because they occur on structures (e.g., the hind legs and abdomen) known to function in combat among males in many species, including components that directly contact rivals and putative weapons or weapon-associated traits that are evolutionarily correlated with other known weapon components. At the same time, we emphasize that individual components may not function as weapons in all species or sexes. Some may instead, or additionally, function in signaling, defense, deflecting force, or locomotion. Together, this combination of documented combat function and putative weapon-associated traits in leaf-footed bugs, as well as their extensive variation across species and sexes (Miller et al. 2024), provide a rare opportunity to examine macroevolutionary patterns of sexual dimorphism in a highly labile, multicomponent weapon system.

Leaf-footed bugs use their incredible diversity of morphological elaborations in multiple ways. Most prominently, males of many species engage in one-on-one physical battles over host plant territories that females need for mating, feeding, and laying eggs. In these cases, elaborated hind legs and associated traits are used as sexually selected weapons by the males. Males signal by raising one or both hind legs, with escalated contests involving ventral-ventral or end-to-end grappling and squeezing. Males that establish territories achieve greater mating, and likely fertilization, success (Miller et al. 2024).

Females of some species lack elaborations entirely, express components very similar to males, or exhibit elaborations different in, for example, location, shape, and/or size. When female hind legs are elaborated, it is not uncommon for females to kick and scrape at other individuals using spines on their femora or tibia. These behaviors appear to be used to secure feeding and/or oviposition sites (Eberhard 1998)( M. Forthman, C. W. Miller, pers. obs.) or reject mating attempts by males (J. Yan, pers. obs.). Sometimes female fights are as aggressive and involved as male fights (C.W. Miller pers. obs.), but as of yet no females have been documented fighting with other females over mating opportunities. Thus, females can use their legs and associated components as weapons, but no evidence exists to-date that these weapons are subject to sexual selection in females.

As is the case in many species, the weapon system of leaf-footed bugs serves multiple functions (Miller et al. 2026). In addition to use in competition over mating opportunities, feeding, and oviposition sites, some traits likely serve in deflection or defense against predation (Evans-Blake et al. 2026). Further, both females and males must walk using their hind legs, even when highly elaborated. Thus, the hind legs and other components of the weapon system have likely been selected from multiple sources over evolutionary time, including some not yet known.

Here, we capitalized on the incredible diversity and complexity of elaborate weapon and weapon-associated traits in the leaf-footed bugs to investigate the evolutionary dynamics of sexual dimorphism in their multicomponent, mechanical weapon system. Drawing from a newly assembled morphological dataset of 14 commonly elaborated traits for 154 species and a new phylogenetic framework, we tested several predictions.

First, we predicted that females are generally larger-bodied across species, whereas males exhibit greater exaggeration in weapon components. Prior work suggests that sexual dimorphism in leaf-footed bugs is not simply a matter of one sex being larger overall. In several well-studied coreids, females are larger-bodied than males, whereas males of some species, such as *Acanthocephala declivis* (Say 1832), *Leptoscelis tricolor* (Westwood 1842), *Narnia femorata* (Stål 1862), and *Thasus neocalifornicus* Brailovsky & Barrera, 1994 (in (Brailovsky et al. 1994), exhibit disproportionately enlarged hind femora after accounting for body size (e.g., (Eberhard 1998; Miller and Emlen 2010; Allen and Miller 2020; Graham et al. 2022). Thus, body size and weapon-associated traits may follow different sex-specific evolutionary trajectories.

Second, given prior evidence that weapons are evolutionarily labile in male leaf-footed bugs (Miller et al. 2024), we predicted frequent gains and losses of traits, as well as transitions among sexually monomorphic and dimorphic states in weapon components. However, because individual components may differ in function, development, and degree of integration with other components, we also predicted that these patterns would vary among traits and between sexes, with some components showing, for example, repeated gains of male-biased expression and others showing similar numbers of transitions in males and females or greater turnover in females.

Third, we predicted that the multicomponent weapon system of leaf-footed bugs would reveal a broader set of evolutionary pathways to and from sexual dimorphism than has typically been recovered in prior macroevolutionary studies. Previous macroevolutionary studies of weapons—often focused on simpler weapon systems or focused on fewer components and/or properties—have recovered or permitted the inference of only a subset of theoretically possible transitions, with some pathways inferred rarely or not at all (Table 1). Because leaf-footed bugs possess a highly complex weapon system in which numerous easily observed components vary in presence and expression between the sexes, we expected to recover multiple transition types and to determine whether these pathways occur at similar frequencies or whether one pathway predominates relative to others.

Lastly, we predicted that the overall degree of sexual dimorphism across the entire weapon system will vary phylogenetically. Prior work in leaf-footed bugs showed that most male weapon components are correlated with one another and accumulated unevenly across the phylogeny, producing some clades in which males possess much more complex, multicomponent weapon systems compared to other clades (Miller et al. 2024). Because sexual dimorphism of the weapon system reflects the difference between the sexes across all these components and females often express reduced or different forms of components or lack them altogether, we predicted that clades with the most highly elaborated male weapons would exhibit the most extreme, sexually dimorphic weapon system. Furthermore, because most of these components are correlated with one another, at least in males, we predicted that increases in overall sexual dimorphism would appear abruptly rather than through gradual changes.

By integrating a multicomponent perspective with phylogenetic comparative approaches, this work provides a detailed view of how sexual dimorphism evolves within complex functional trait systems.

## Methods and Material

### Molecular data collection and phylogenetic inferences

For specific details on our molecular data collection and phylogenetic approaches, see Supplementary Materials. In brief, we assembled a phylogenomic dataset of 3,040 ultraconserved element (UCE) loci for 336 taxa, combining newly generated sequence-capture data for 20 taxa with previously published UCE datasets and genome/transcriptome assemblies (Tables S2–S3). All DNA extraction, library preparation, and enrichment procedures were described in (Forthman et al. 2019, 2020, 2022, 2023).

Demultiplexed, adapter-trimmed, and error-corrected sequence reads were assembled into contigs following (Forthman et al. 2022). The UCE loci were aligned and screened for orientation issues, trimmed to remove poorly aligned regions, and filtered to produce two datasets: a matrix of loci represented in ≥50% of taxa (“50p”) and a subset containing the 25% most parsimony-informative loci (“50p25mi”). A summary of alignment lengths, variable sites, and numbers of loci for both datasets can be found in Table S4. Phylogenetic analyses were conducted under both maximum likelihood (ML) and multispecies coalescent (MSC) frameworks. Maximum-likelihood trees were inferred from concatenated alignments using IQ-Tree v2.1.3 (Minh et al. 2020) with model partitioning, and MSC species trees were estimated using weighted ASTRAL-hybrid (wASTRAL-h), implemented in ASTER v1.22.3.7 (Zhang and Mirarab 2022; Zhang et al. 2025).

Prior to downstream ancestral state estimation (ASE) analyses, ML and MSC topologies were converted to ultrametric trees after removing outgroup taxa, as well as 15 ingroup taxa that displayed extremely long branches relative to closely related ingroup taxa. These ingroup taxa had the fewest UCE loci recovered (52–278 loci) and the highest levels of mean within-locus missing data (∼46–78%). This combination of low locus representation and short sequences likely contributed to their extreme branch lengths during phylogenetic inference. Because ASE analyses depend on accurate branch lengths for inferring ancestral states and character transitions, we excluded these taxa from subsequent analyses.

### Statistical evaluation of sexual dimorphism across 27 species

#### Morphometric dataset

To characterize the prevalence of sexual dimorphism in leaf-footed bug weapons, identify which components are sex-limited versus shared, and assess whether males exhibit greater exaggeration, we assembled a morphometric dataset for 27 species represented by at least four specimens of each sex, with species-level sample sizes ranging from 10 to 58 individuals (hereafter the “pGLMM taxa” dataset; Table S5). For each species, we measured pronotal width as a proxy for body size (Procter et al. 2012) and up to 15 weapon components in males and females: length of the longest process on the (1) metapleuron, (2) coxa, (3) trochanter, (4) dorsal femoral surface, (5) anterior/posterior femoral surface (hereafter lateral femoral surface), (6) ventral femoral surface, (7) internal tibial surface (i.e., ventral surface), (8) external tibial surface (i.e., dorsal surface), and (9) ventral abdominal surface; greatest femoral width across the (10) lateral and (11) dorsal surfaces; lengths of the (12) internal and (13) external tibial flags; and the widths of the (14) internal and (15) external tibial flags. Because the number and position of processes often differ between sexes and among species, homology assignments for the processes were not possible. We therefore measured the longest process on each structure to capture the maximum extent in which the component can be exaggerated. Although some of these components are rare across the species available for quantitative analysis (i.e., processes on the metapleuron, coxa, trochanter, and ventral abdomen were expressed in one or two species), we retained them because they are discrete components included in the broader coreoid weapon system described by (Miller et al. 2024). Furthermore, we retained these uncommon components because for some of these they can be conspicuous and strongly sexually dimorphic in species that express them (pers. obs.; also see Results). Measuring these traits allowed us to document their sex-specific size and scaling, even when their limited occurrence prevented their inclusion in phylogenetic comparative analysis. All components were measured using a Nikon SMZ1500 stereomicroscope and Leica Application Suite X v5.1.0.25593 (Leica Microsystems).

We used descriptive statistics to evaluate component presence and sex-specific distributions across species, including the range, mean, and median component sizes. We further examined allometric scaling relationships between component size and body size by fitting linear regressions of log-transformed component values against log-transformed pronotal width for each sex per species.

We assessed each component for normality and homogeneity of variances using Shapiro–Wilk and Levene’s tests, respectively (Levene 1960; Shapiro and Wilk 1965). Multiple-testing corrections were applied using the Benjamini–Hochberg false discovery rate procedure (Benjamini and Hochberg 1995). Because components violated assumptions of normality and/or homoscedasticity, all measurements were log-transformed prior to statistical analyses.

#### Phylogenetic generalized linear mixed models

We evaluated sex differences in components while accounting for shared ancestry using phylogenetic generalized linear mixed models (pGLMMs), implemented with the *pglmm* function in *phyr* v1.1.3 (Li et al. 2020) in R v4.5.1 (R Core Team 2025). To account for phylogenetic uncertainty, we evaluated our models using each of the four ultrametric trees. First, we compared two models for pronotal width, which both included a Gaussian error distribution and sex as a fixed effect. However, one model included species identity as a random effect (1|Species.ID) while the other one included a phylogenetic covariance random effect in addition to a random effect for species identity (1|Species.ID). For each of the 11 components expressed in seven or more species, we tested the same models, as well as additional models that included pronotal width as a covariate with a sex-by-pronotal-width interaction. When the interaction term was not statistically significant, we analyzed a reduced model that excluded the interaction term. All these additional models included species identity with and without a phylogenetic covariance random effect. We evaluated model fit using the Akaike Information Criterion (AIC) and likelihood ratio tests (LRTs).

#### Species-specific generalized linear models for rare components

Four components—processes on the metapleuron, coxa, trochanter, and ventral abdominal surface—were expressed in only one or two species and therefore could not be analyzed using pGLMM. For these components, we instead fitted non-phylogenetic generalized linear models (GLMs) independently within each species using the base-R *glm* function. For each component-species combination, we evaluated three models with Gaussian error distributions: (1) component ∼ sex, (2) component ∼ sex × pronotal width, and, when the interaction term was non-significant, (3) component ∼ sex + pronotal width. Model support was also assessed using AIC and LRTs.

### Characterizing sexual dimorphism and inferring evolutionary transitions across 154 species

#### Qualitative coding of weapon components and sexual dimorphism

To broaden our understanding of sexual dimorphism across leaf-footed bugs, we assembled a second dataset comprising 154 species (hereafter the “all-taxa” dataset). Because limited specimen availability prevented quantitative analyses of weapon components within species, this dataset relied on conservative qualitative coding of sexual dimorphism. For each species, we examined males and females to assess the presence and expression of components evaluated in the pGLMM dataset, with some exceptions. First, lateral and dorsal femoral widths were treated as a single “femoral enlargement” component. Unlike processes, the femur is always present in both sexes, and thus, this character captures whether the femur is conspicuously enlarged in one sex relative to the other. Second, for both the internal and external tibial flags, the lengths and widths were combined into a single component. Treating flag length and flag width as separate components in an ASE analysis would effectively count the same structure twice and could imply independent gains or losses of different components when the underlying change concerns the size or shape of a single flag. Additionally, femoral and tibial curvatures were included to capture exaggerated shapes that cannot be adequately represented by linear measurements. In total, 14 weapon components were visually evaluated for sexual dimorphism. As in our statistical evaluations of sexual dimorphism, we retained rare components in the all-taxa dataset because they may represent lineage-specific parts of the weapon system and can contribute to changes in overall weapon complexity and sexual dimorphism. Their inclusion allowed our downstream analyses to capture the full range of weapon complexity rather than only the most commonly expressed weapon components. For components involving processes, we followed the same rationale used in the pGLMM dataset and evaluated sexual dimorphism based on the longest process in each sex. To account for body size, we size-matched male and female specimens using pronotal width as a proxy for overall body size. This was necessary because females are often larger-bodied in well-studied coreids, whereas male weapon components can be disproportionately enlarged relative to body size in some species. When available, multiple male–female pairs for each species representing a range of body sizes (e.g., smallest pair, largest pair, and an intermediate pair) were examined. We selected a range of body sizes because, where sexual dimorphism in weapon components has been evaluated within a species, it is often disproportionately greater at the largest body sizes and minimal at small body size (e.g., (Eberhard 1998; Miller and Emlen 2010). Only specimens in good condition (i.e., no deformities or evidence of severe wear) were used.

For each component in each species, we assigned a discrete sexual dimorphism state, with all visual evaluations made by a single observer (MF) to maintain consistency in coding and minimize observer bias. Components involving processes and tibial flags were coded as: (0) Absent in both sexes, (1) Present in both sexes and apparently monomorphic, (2) Present in males only, and (4) Present in both sexes and distinctly dimorphic. Femoral enlargement was coded as a binary character: (0) Apparently monomorphic and (1) Distinctly dimorphic. Femoral and tibial curvatures were coded as: (0) Straight in both sexes, (1) Curved in males only, and (2) Curved in both sexes and apparently monomorphic. For each component, sexual dimorphism assessments were based on all observable properties—such as the relative size, number, and distribution of processes—such that the presence of sexual dimorphism in any one of these properties resulted in the component being coded as sexually dimorphic. While these finer-scale features informed our qualitative assessment of whether a component was monomorphic or dimorphic, they were not analyzed as independent components in our comparative analyses. We also note that some species coded as sexually monomorphic may in fact exhibit subtle sexual dimorphism that is not detectable through visual inspection. Thus, we employed a conservative coding framework in which only clear, conspicuous differences were classified as sexually dimorphic. We then summarized component presence by sex across species and identified which sex expressed the more exaggerated condition for components present in both sexes and that were dimorphic.

#### Ancestral state estimation

To infer how sexual dimorphism in components evolved across leaf-footed bugs, we performed ASE analyses on each of the components in the all-taxa dataset using each of the four ultrametric trees. For each component, we first pruned the phylogeny to match the taxa coded for the component. We then modeled character evolution under three Markov models—equal-rates (ER), symmetric (SYM; equivalent to ER for binary characters), and all-rates-different (ARD)—using the *fitMk* function in the R package *phytools* v2.4-4 (Revell 2012, 2024). Following the model-averaging procedure in Revell (2024), we compared ER, SYM, and ARD models using AIC and calculated AIC weights to quantify the relative support for each model. Rather than selecting a single best model, we performed model-averaged ASE in which stochastic character maps were weighted according to the AIC weights for each model. Sampling stochastic maps across multiple models allows the analyses to incorporate model uncertainty, weighting each model’s contribution according to the support it receives (Revell 2024). For each component, we generated 1,000 stochastic character maps using the *make.simmap* function under the model-average weighting scheme. For each transition between two states, the *density* function summarized the distribution of the number of occurrences per stochastic map and reported the median, mean, range, and 95% highest posterior density (HPD) interval across 1,000 maps. These summaries therefore described how often each individual type of state change occurred within a sampled history but do not retain information about the sequence of multiple transitions within lineages. For example, separate counts of transitions from absence to present in both sexes and apparently sexually monomorphic and from the latter condition to sexual dimorphism do not indicate how often those changes occurred successively along the same lineage.

Thus, we also examined the ordered character histories retained within each stochastic character map to quantify complete evolutionary “routes” from plesiomorphic conditions (i.e., absence of a component; see Results) to the first appearance of sexual dimorphism. For every inferred origin of component presence in both sexes that is sexually dimorphic, we traced the lineage forward from the plesiomorphic state until it first reached a sexually dimorphic state, defined as presence in males only or presence in both sexes with distinct sexual dimorphism. If a lineage returned to the plesiomorphic state before reaching a sexually dimorphic state, the route was reset from that most recent plesiomorphic occurrence. Once a sexually dimorphic state was reached, subsequent transitions between sexually dimorphic states were not included in that route and were not counted as additional origins unless the lineage first returned to the plesiomorphic state. We counted the occurrence of each complete route separately within each stochastic map and summarized route frequencies across the 1,000 maps using the median, mean, range, and 95% HPD interval. We performed this route analysis only for component characters with three or more modeled states; the binary femoral-enlargement character was therefore excluded.

#### Degree of sexual dimorphism

*across the weapon system*. We also generated a sexual dimorphism score (SDS) for each of the 154 species from these components, along with additional properties of these components not directly incorporated into our all-taxa and pGLMM datasets (e.g., number and distribution of processes along a surface; Table S6). For each species, a component property received a score of 0 if it appeared sexually monomorphic or 1 if sexually dimorphic, and the SDS was calculated by summing property scores across all components. Species were then assigned to one of four discrete categories representing increasing degrees of overall sexual dimorphism: (0) Monomorphic (SDS = 0), (1) Limited sexual dimorphism (SDS = 1–4), (2) Moderate sexual dimorphism (SDS = 5–8), and (3) Extreme sexual dimorphism (SDS = 9–14). We used the same ASE protocol outlined above for the individual weapon components to summarize evolutionary transitions in the degree of sexual dimorphism across Coreoidea.

## Results

### Phylogeny of the Coreoidea

Our phylogenomic analyses recovered a topology for the Coreoidea (Fig. 2, Supplementary Files 1–4), that is largely congruent with prior UCE-based investigations, including the non-monophyly of the families Coreidae and Alydidae, the presence of the Alydidae + Hydarinae + Pseudophloeinae clade, and the major lineage arrangements within the Coreinae + Meropachyinae (Forthman et al. 2019, 2020, 2022, 2024; Kieran et al. 2019; Emberts et al. 2020; Miller et al. 2022, 2024). Detailed descriptions of these previously recovered relationships, along with their corresponding nodal support values across our ML and multispecies coalescent MSC phylogenies, are provided in the Supplementary Material. Here, we only report novel relationships recovered in UCE-based phylogenomic analyses.

**Figure 2.**
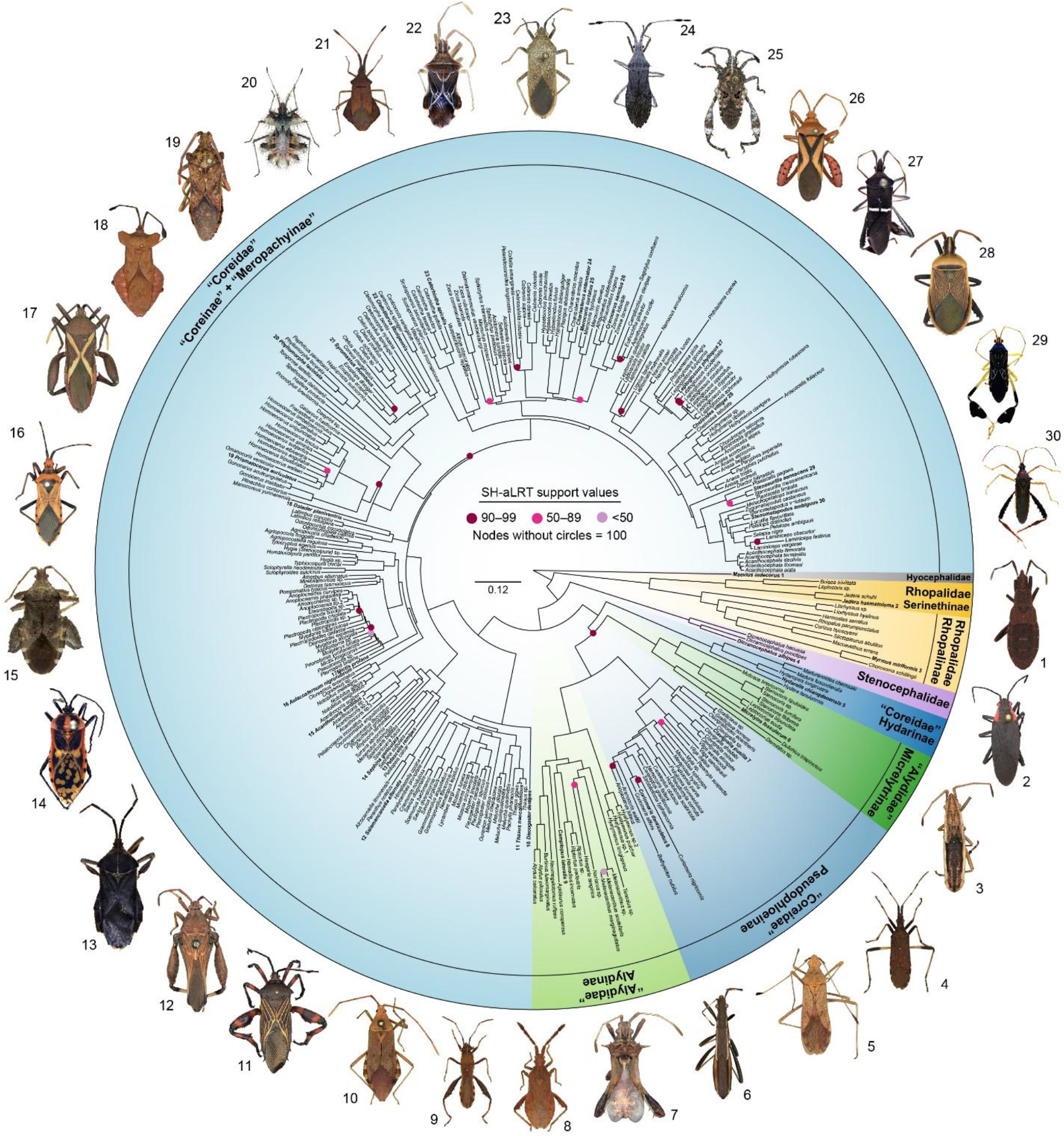
Maximum likelihood (ML) best tree of the Coreoidea based on the 50p concatenated ultraconserved element alignment. For visualization, outgroup taxa are pruned from the tree. Circles at nodes represent Shimodaira-Hasegawa-like approximate likelihood ratio test (SH-aLRT) support; nodes without circles have 100% support. Dorsal habitus images of select representatives (not to scale) are given to show a range of diversity within the Coreoidea (terminal taxa with images bolded and have corresponding numbers associated with images).

Across all analyses, Coreoidea was recovered as a fully supported clade, with Hyocephalidae consistently forming the earliest-diverging lineage sister to all remaining coreoids. Rhopalidae was recovered as a monophyletic group with high support and placed as sister to a clade comprising Stenocephalidae + Alydidae + Coreidae (ML 50p/50p25mi = 100/100; MSC 50p/50p25mi = 0.9998/0.9999). Within Rhopalidae, the subfamilies Rhopalinae and Serinethinae were each monophyletic with full support. However, within Rhopalinae, the tribe Rhopalini was rendered paraphyletic by Chorosomatini and Harmostini, with all relationships within this clade highly supported (SH-aLRT = 100, LPP = 0.9155–1). Finally, Stenocephalidae was recovered as monophyletic and strongly supported as the sister group to Alydidae + Coreidae (100/100/1/0.976).

Within the Coreinae + Meropachyinae, our analyses also expanded upon the previously documented polyphyly of the tribes Anisoscelini and Hypselonotini by recovering one additional independent lineage for each. Specifically, the anisosceline *Dalmatomammurius vandoesburgi* (Brailovsky 1982) was recovered as sister to *Zicca* (Hypselonotini) with poor support in ML analyses (50p/50p25mi = 57.3/53.6) or with the *Acidomeria* + *Sethenira* + *Sphictyrtus* + *Mamurius* hypselonotine lineage (*Mamurius* currently *incertae sedis* in tribal placement) with higher support in MSC analyses (50p/50p25mi = 0.9426/0.9935). The hypselonotine *Namacus annulicornis* (Stål 1870), was nested within a large, fully supported clade of anisoscelines (*Leptoglossus* + *Leptoscelis* + *Narnia* + *Phthia* + *Phthiacnemia* + *Phthiadema* + *Ugnius*).

#### Statistical evaluation of sexual dimorphism across 27 species

For all 11 components analyzed using pGLMMs, patterns of sexual dimorphism were highly consistent across all four phylogenic hypotheses. Thus, we report results from the ML 50p phylogeny, with full model comparisons for all trees provided in Table S7. Across components, sex was consistently an important predictor, with body size often influencing the magnitude of sexual dimorphism. Phylogenetic covariance improved model fit for some components but not others, suggesting that shared ancestry contributed unevenly across traits.

Females were generally larger-bodied than males across the sampled taxa (Tables S5, S7). Mean pronotal width was greater in females compared to males in 23 of 27 species. The four exceptions were *Acanthocephala femorata* (Fabricius 1775) (4.2% smaller; n = 29 males, 28 females), *Camptischium clavipes* (Fabricius 1803) (14.4% smaller; n = 6, 4), *Leptoglossus occidentalis* (Heidemann 1910) (5.7% smaller; n = 10, 10), and *Merocoris curtatus* (McAtee 1919) (1.0% smaller; n = 27, 25). Averaged across individuals of all species, females were 4.6% larger overall. At the species-level, average female mean pronotal width exceeded those of males by ∼5.8%. This pattern was supported by the best-fitting pGLMM that included sex as a fixed effect and both species identity and phylogenetic covariance as random effects (sex coefficient = -0.0586, p < 2.20e^-16^).

Weapon components showed two broad patterns of sexual dimorphism with respect to body size (Figs. 3, S1). For some components, the difference between males and females changed with body size, such that one sex became increasingly exaggerated relative to the other among larger individuals. For other components, one sex was consistently larger across the observed body-size range, with little evidence that the magnitude of the sex difference changed with body size.

**Figure 3.**
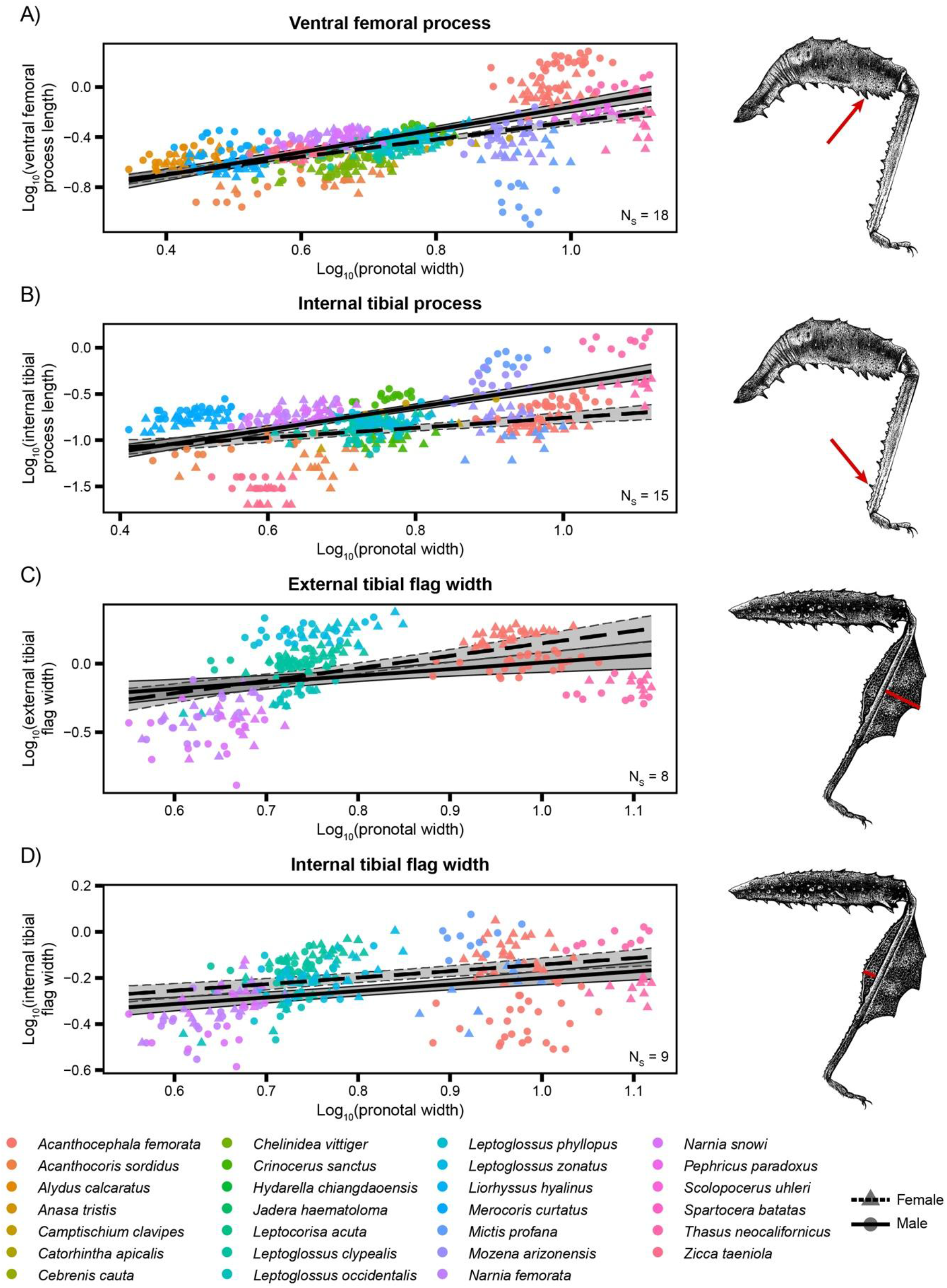
Males show greater exaggeration in most selected weapon components, but tibial flag width is female-biased. Sex-specific allometric relationships are shown for four representative hind leg components of the weapon system of leaf-footed bugs. (A) Ventral femoral process and (B) internal tibial process lengths became increasingly male-biased with increasing body size. (C) External tibial flag width shifted from male-biased among smaller individuals to female-biased among larger individuals, whereas (D) internal tibial flag width remained consistently female-biased across the observed body-size range. Points represent log-transformed component values plotted against log-transformed pronotal width for individual specimens, colored by species. Circles and triangles denote male and female specimens, respectively. Solid and dashed lines indicate male and female trends, respectively. Shaded regions represent 95% confidence intervals (CI), bounded by solid lines for males and dashed lines for females. Number of species (N_S_) is indicated in each plot. For these and all other components not illustrated here, measurements and sample sizes are provided in Table S5, and full pGLMM results are provided in Table S7. Plots for all other components analyzed with PGLMMs are shown in Figure S1. Illustrations of the hind leg of *Camptischium clavipes* (Fabricius, 1803) accompany panels A and B, and those of *Leptoglossus gonagra* (Fabricius, 1775) accompany panels C and D (not drawn to scale; illustrations by David J. Tuss); red arrows and lines indicate the components measured.

Most components followed the first pattern, which generally showed stronger male-biased exaggeration at larger body sizes. For dorsal and lateral femoral width, dorsal and ventral femoral processes, and internal and external tibial processes, female had equal or larger component values among smaller individuals, but males became increasingly exaggerated as pronotal width increased (sex×PW = 0.151–0.693, all p ≤ 0.00146; Figs. 3A, 3B, S1; Table S7). Thus, sexual dimorphism in these traits generally became more pronounced among larger-bodied individuals.

Two components also showed sex-specific allometry but did not follow the same pattern. Lateral femoral process length remained male-biased across the observed body-size range, despite a significant negative sex-by-pronotal-width interaction (sex × PW = -0.236, p = 0.0267; Fig. S1; Table S7). In contrast, external tibial flag width showed a shift in the direction of sexual dimorphism; the flag was larger in males among smaller individuals, but females showed increases with increasing pronotal width (sex×PW = -0.363, p ≤ 0. 4.64E^-7^; Figs. 3C; Table S7).

Several tibial flag traits showed consistent sex differences across the observed range of body sizes. Males had longer internal and external tibial flags than females (sex = 0.0538–0.0780, p ≤ 0.000264), whereas females had wider internal tibial flags than males (sex = -0.1306, p = 1.06E^-5^; Fig. 3D, S1; Table S7). In all three models, pronotal width also had a positive effect on component size (PW = 0.429–0.956, p ≤ 0.000373), but sex-by-pronotal-width interactions were not retained in the best-supported models.

Rare components expressed in one or two species showed more variable patterns of sexual dimorphism. Because the metapleural, coxal, trochanteral, and ventral abdominal processes occurred in too few species, these components were analyzed within species using non-phylogenetic GLMs. Sexual dimorphism was detected in three of the six component–species combinations analyzed, in which males generally expressed larger processes than females (Fig. 4; Table S8). In *Acanthocoris sordidus* (Thunberg 1783), males had longer coxal processes than females across body sizes (sex = 1.019, p = 6.09e^-11^). In *Camptischium clavipes*, females showed a steeper increase with pronotal width than males (sex×PW = - 0.481, p = 0.0452), but males retained larger process lengths across the observed body-size range. In *Mictis profana* (Fabricius, 1803), male had longer ventral abdominal processes than females across the body-size range (sex = 1.732, p = 4.64E^-9^). In contrast, coxal, trochanteral, or ventral abdominal processes in *Pephricus paradoxus* (Sparrman 1777) showed little evidence of sexual dimorphism (sex = -0.0560–0.0040, p ≥ 0.348; Fig. S2; Table S8).

**Figure 4.**
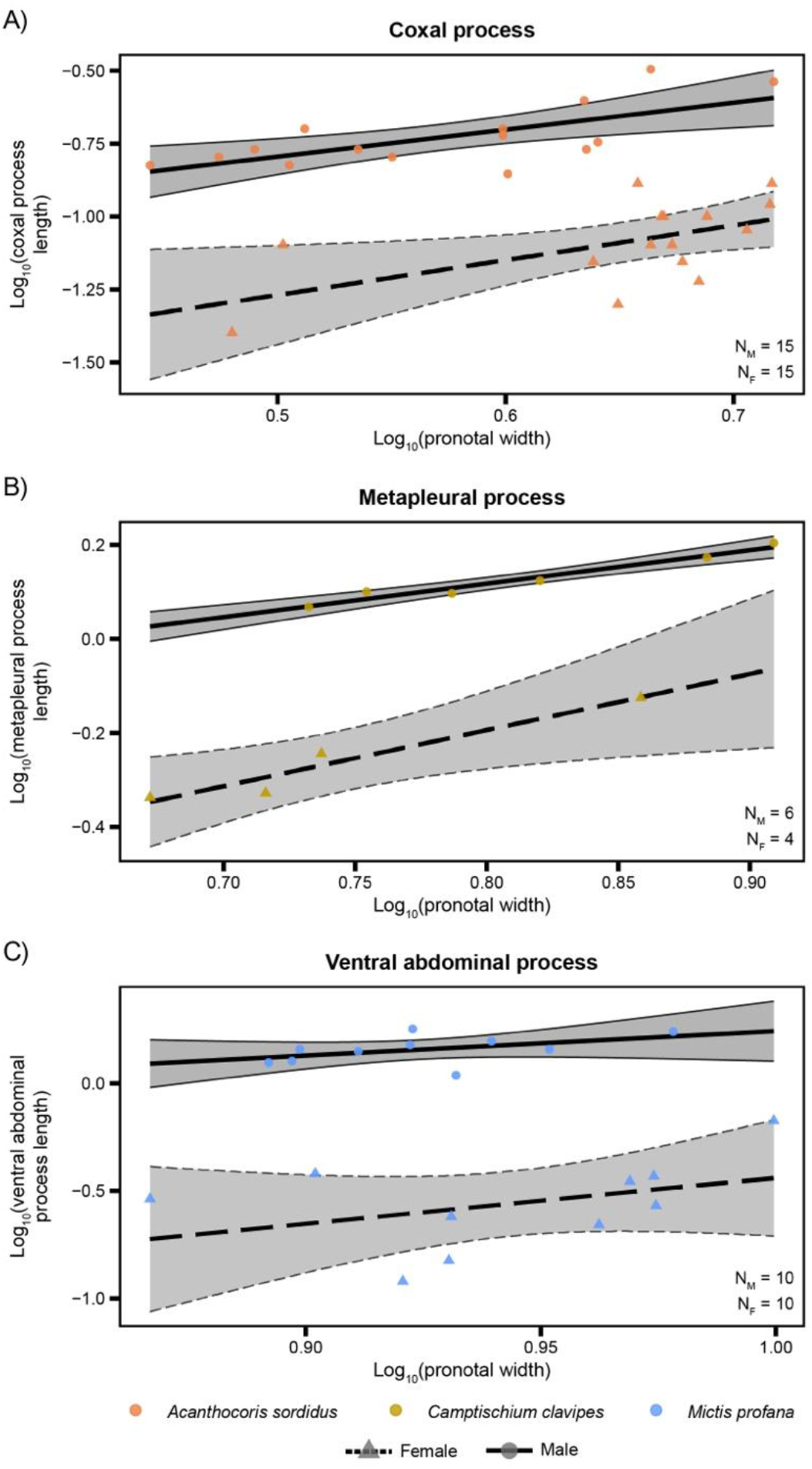
Rare weapon components show male-biased sexual dimorphism. Species-specific allometric relationships are shown for weapon components expressed in only one or two species analyzed: (A) coxal processes in *Acanthocoris sordidus* (Thunberg, 1783), (B) metapleural processes in *Camptischium clavipes* (Fabricius, 1803), and (C) ventral abdominal processes in *Mictis profana* (Fabricius, 1803). Each component was larger in males than in females. Points represent log-transformed component values plotted against log-transformed pronotal width for individual specimens. Circles and triangles denote male and female specimens, respectively. Solid and dashed lines indicate male and female trends, respectively. Shaded regions represent 95% CI, bounded by solid lines for males and dashed lines for females. The number of male (N_M_) and female (N_F_) specimens are indicated in each plot. Measurements and sample sizes are provided in Table S5, and full GLM results are provided in Table S8. For allometric relationships corresponding to the coxal, trochanteral, and ventral abdominal processes in *Pephricus paradoxus* (Sparrman, 1777), see Fig. S2.

### Characterizing sexual dimorphism and inferring evolutionary transitions across 154 species

Across the 154 species included in the all-taxa dataset, weapon components varied widely in both presence and sexual dimorphism (Fig. 5). Most species in the Alydinae + Pseudophloeinae and Coreinae + Meropachyinae clades expressed at least one of the 14 components examined, and the number and types of components differed substantially among lineages. Most of the hind femoral and tibial components were the most frequently expressed overall, whereas those on the metapleuron, coxae, trochanters, and abdomen were comparatively rare. For most components, expression in both sexes—either appearing sexually monomorphic or dimorphic—was more common, but male-only expression was particularly frequent for processes on the coxae, trochanters, and abdomen, as well as for femoral and tibial curvatures. For those components expressed in both sexes, sexual dimorphism was observed across many species, while sexual monomorphism appeared slightly less frequently.

**Figure 5.**
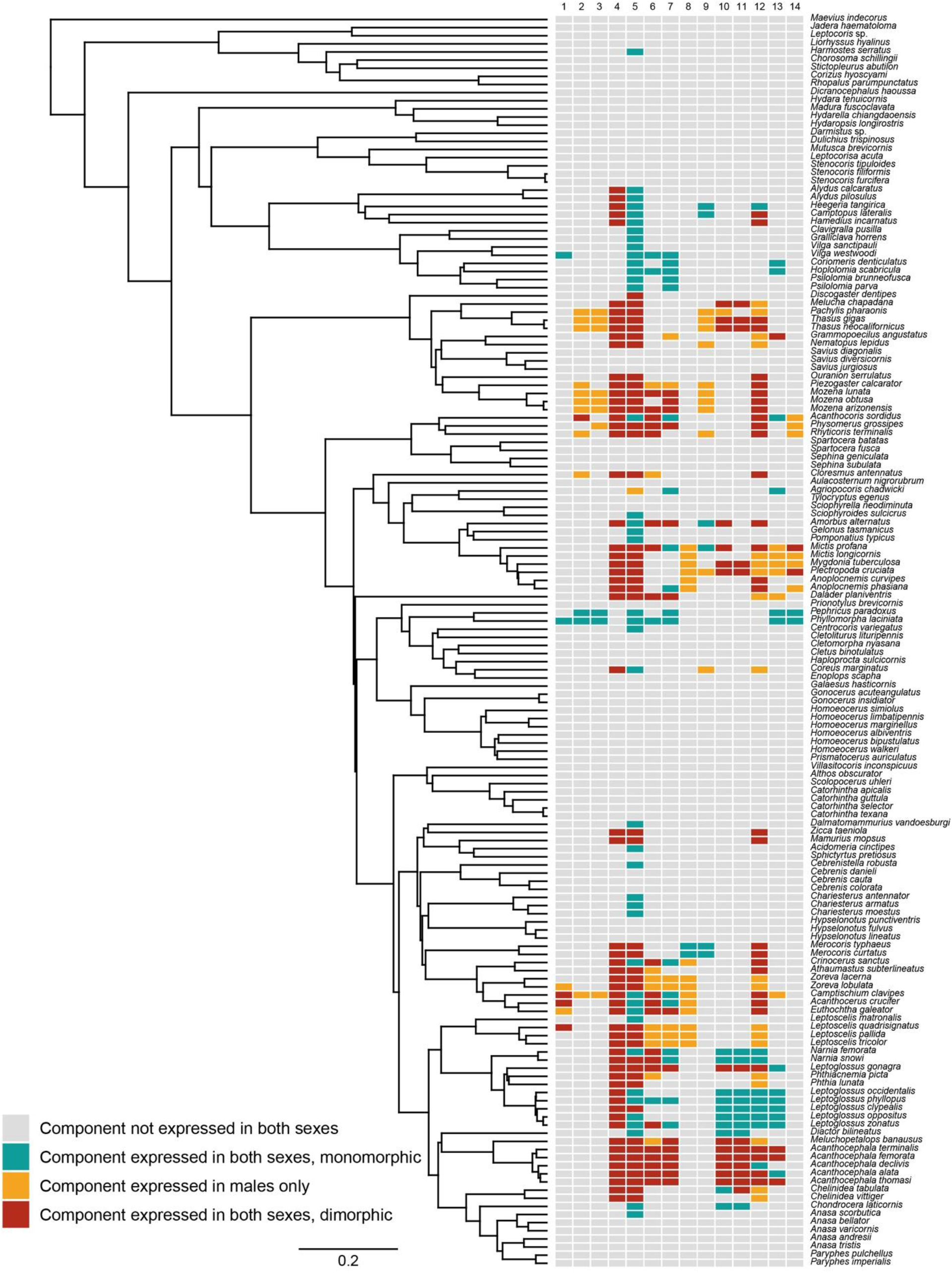
Ultrametric tree based on the ML 50p best tree, with components and sexual dimorphism states displayed for terminal taxa on the right. Names of terminal taxa are removed for visualization purposes. Numerical labels above boxes correspond to the following components: (1) metapleural process, (2) coxal processes, (3) trochanteral processes, (4) femoral enlargement, (5) ventral femoral processes, (6) lateral femoral processes, (7) dorsal femoral processes, (8) femoral curvature, (9) tibial curvature, (10) internal tibial flag, (11) external tibial flag, (12) internal tibial processes, (13) external tibial processes, and (14) ventral femoral processes.

Ancestral state estimation of sexual dimorphism in weapon components was broadly congruent across the four phylogenetic datasets in terms of inferred ancestral states, the relative frequency of transitions among states, and the dominant transition pathways recovered for individual components. As such, we primarily present the ML 50p ASE results below, with analyses based on the other ML and MSC topologies noted only where they differed meaningfully. Across components, ASE recovered substantial heterogeneity in both the number and direction of transitions among sexual dimorphism states, with some components exhibiting extensive evolutionary turnover and others showing relatively few inferred changes (Table S9).

To summarize sex-specific patterns of component gain and loss independent of sexual dimorphism, we quantified the minimum median numbers of gains and losses of each components in males and females across the phylogenies (Tables 2, S9); femoral enlargement was excluded from these summaries because it was coded as sexual dimorphism in the degree of enlargement of a structure always present in both sexes, allowing inference of gains and losses of sexual dimorphism but not sex-specific gains or losses of the femur itself. These sex-specific patterns were largely congruent across alternative phylogenetic hypotheses, although the relative magnitude of gains and losses for the lateral femoral processes varied among trees, particularly under the MSC 50p25mi topology (Table S9). When comparing between sexes, males exhibited higher numbers of inferred gains for several components: the metapleural, coxal, trochanteral, dorsal and ventral femoral, and external tibial processes, as well as the femoral and tibial curvatures. In contrast to males, females exhibited higher numbers of gains for the internal tibial flags and lateral femoral, internal tibial, and ventral abdominal processes. Gains were equal between sexes for the external tibial flag. Losses were higher in females than males for the metapleural and lateral and dorsal femoral processes. In contrast, losses were higher in males for the femoral and tibial curvatures, as well as the internal and external tibial processes. All other components exhibited similar patterns of gains and losses between sexes.

**Table 2.**
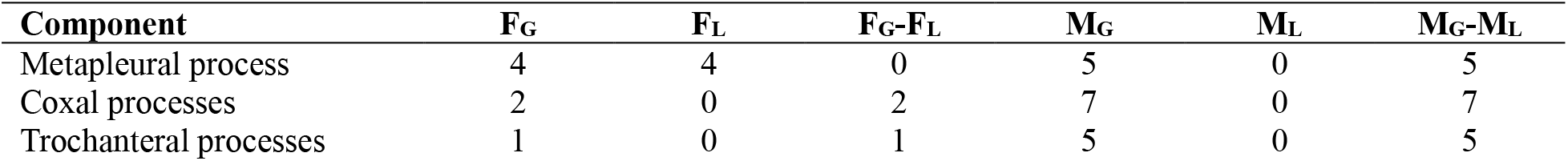
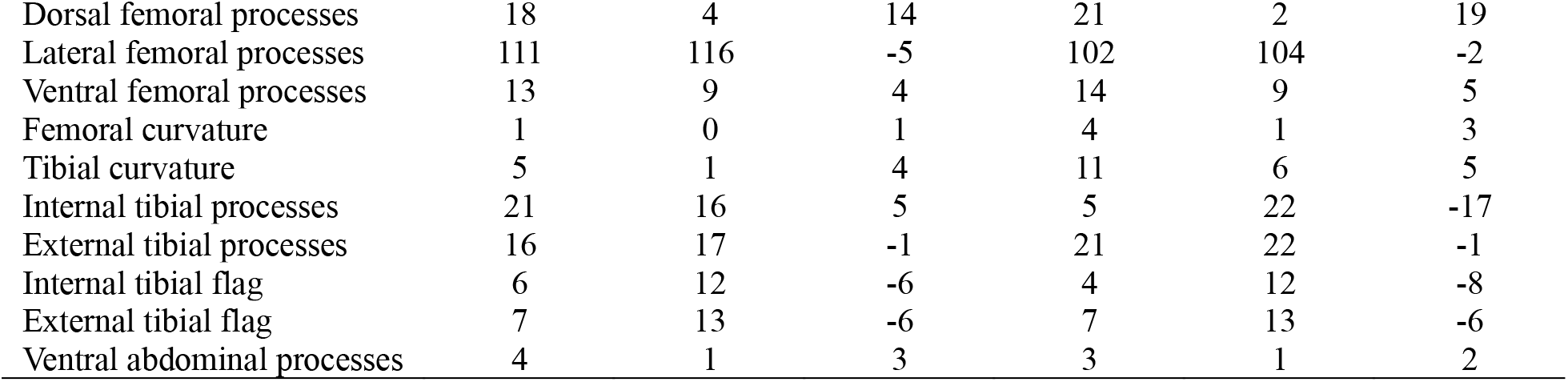
Minimum median numbers of gains and losses of weapon components in females and males inferred under the ML 50p phylogenetic hypothesis. Values were derived from stochastic character mapping and summarize transitions between component absence and presence. For each component, columns report the minimum median number of gains (F_G_, M_G_) and losses (F_L_, M_L_) inferred for females and males, respectively, along with net gains for each sex (F_G_–F_L_; M_G_–M_L_). Femoral enlargement is excluded because the femur is present in both sexes in all taxa.

When patterns of gains and losses were compared within each sex, gains exceed losses for many of the weapon components examined. Only the lateral femoral and external tibial processes, as well as both tibial flag components, exhibited more losses than gains in both sexes. Additionally, the internal tibial processes were more frequently lost than gained in males. For metapleural processes, gains and losses were equal in females but gains predominated in males.

To summarize evolutionary changes in sexual dimorphism independent of overall component presence, we quantified the minimum median numbers of gains and losses of sexual dimorphism— including sex-limited expression or components dimorphically expressed in both sexes—across phylogenies (Tables 3, S9). Several components exhibited net gains of sexual dimorphism, with several— the metapleural, coxal, trochanteral, and dorsal femoral processes—having no corresponding losses.

**Table 3.** Minimum median numbers of gains and losses of sexual dimorphism in weapon components inferred under the ML 50p phylogenetic hypothesis. Sexual dimorphism includes sex-limited expression and dimorphic expression in both sexes. Values were derived from stochastic character mapping and report the minimum median number of gains (SD_G_), losses (SD_L_), and net change (SD_G_–SD_L_) of sexual dimorphism for each trait.

| **Component** | **SD<sub>G</sub>** | **SD<sub>L</sub>** | SD<sub>G</sub>–SD<sub>L</sub> |
|---|---|---|---|
| Metapleural process | 3 | 0 | 3 |
| Coxal processes | 6 | 0 | 6 |
| Trochanteral processes | 4 | 0 | 4 |
| Femoral enlargement | 17 | 5 | 12 |
| Dorsal femoral processes | 11 | 0 | 11 |
| Lateral femoral processes | 14 | 16 | -2 |
| Ventral femoral processes | 19 | 11 | 8 |
| Femoral curvature | 3 | 1 | 2 |
| Tibial curvature | 6 | 5 | 1 |
| Internal tibial processes | 10 | 28 | -18 |
| External tibial processes | 21 | 24 | -3 |
| Internal tibial flag | 5 | 6 | -1 |
| External tibial flag | 10 | 10 | 0 |
| Ventral abdominal processes | 2 | 1 | 1 |

Femoral enlargement and ventral femoral processes also showed relatively high net gains of sexual dimorphism despite some losses. In contrast, lateral femoral, internal tibial, and external tibial processes exhibited net losses of sexual dimorphism, which was particularly high for the internal tibial processes. Several components showed balanced turnover between gains and losses of sexual dimorphism, including the tibial curvature and tibial flags.

All major evolutionary pathways by which sexual dimorphism was gained and lost were recovered across weapon components, excluding femoral enlargement because it was not coded as a presence–absence character (Fig. 6A; Table S9). These pathways were inferred from stochastic character maps by counting transitions among discrete component states. When aggregated across these components, the most frequently inferred pathway to sexual dimorphism was the origin of a new component in both sexes that was already sexually dimorphic (median = 44 transitions; Fig. 6A); in other words, this pathway represents cases in which a component was inferred to shift from absent in both sexes to present in both sexes, but with males and females already differing conspicuously in the component’s expression. This pattern was especially evident for external tibial processes (median = 16 transitions) and lateral femoral processes (median = 14 transitions). Gains of male-limited sexual dimorphism from absent or straight components in both sexes (Fig. 1, blue arrow) were also common (median = 37 transitions). In contrast, transitions from components already present in both sexes but apparently sexually monomorphic to a sexual dimorphic state (Fig. 1, orange arrows) were inferred less frequently overall; sexual dimorphism more often arose via phenotypic divergence of components retained in both sexes (median = 28 transitions) than through loss of expression in females (median = 5 transitions). Reversions from sexual dimorphism to sexually monomorphic expression were inferred along all major pathways by which sexual dimorphism evolved (Fig. 6A) and occurred at frequencies comparable to gains of sexual dimorphism.

**Figure 6.**
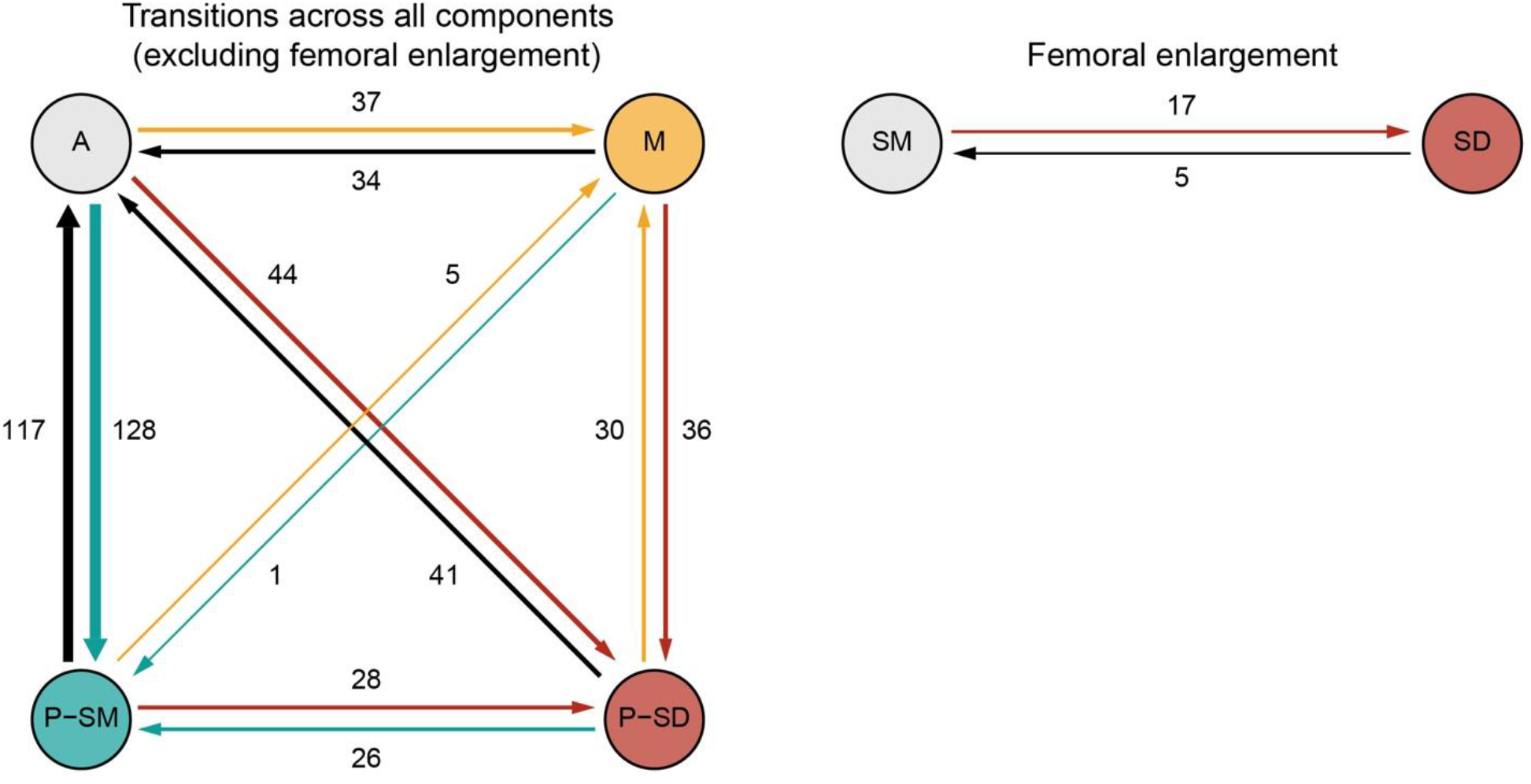
Sexual dimorphism evolves through multiple evolutionary transitions, with the most frequent transition involving the origin of new components that are already sexually dimorphic and with gains and losses of sexual dimorphism occurring at comparable frequencies across all routes. Values shown represent median transition counts. Arrow widths are proportional to median numbers of inferred transitions. (A) Summary of median numbers of transitions among component absence (A), sexually monomorphic expression of components present in both sexes (P–SM), male-limited component expression (M), and sexually dimorphic expression of components present in both sexes (P–SD), aggregated across all weapon components excluding femoral enlargement, based on stochastic character mapping under the ML 50p phylogeny. Sexual dimorphism includes both male-limited expression and dimorphic expression in components present in both sexes. (B) Corresponding transition summary for femoral enlargement, shown separately due to the femora always being present in both sexes (i.e., lacks an absence state).

With regards to femoral enlargement, which lacks an absence state, transitions to sexual dimorphism were inferred relatively frequently (median = 17 transitions; Fig. 6B). In contrast, transitions from sexually dimorphic to sexually monomorphic were recovered at roughly one-third the frequency of gains of sexual dimorphism (median = 5 transitions).

Because these transition counts do not indicate whether successive changes occurred along the same evolutionary lineage, we additionally examined complete ordered routes from component absence or a straight component to the first occurrence of sexual dimorphism (Tables 4, S10). These analyses revealed substantial variation among components in how sexual dimorphism originated. For several components, sexual dimorphism arose predominantly through direct routes. For example, direct origins of male-limited expression were especially common for coxal and trochanteral processes and for femoral and tibial curvature, whereas direct origins of components already distinctly dimorphic in both sexes predominated for lateral femoral and external tibial processes (Table 4). In contrast, ventral femoral processes predominantly acquired sexual dimorphism through an indirect route, first arising in both sexes as apparently sexually monomorphic before becoming distinctly dimorphic (median = 14 routes). The external tibial flag showed equal median numbers of direct and indirect origins of distinct sexual dimorphism (5 each).

**Table 4.** Median numbers of complete evolutionary routes to the first sexually dimorphic state inferred under the ML 50p phylogenetic hypothesis. Values were derived from stochastic character mapping. Direct routes reached a sexually dimorphic state directly from an absent or straight, whereas indirect routes passed through sexually monomorphic expression in both sexes before reaching the first sexually dimorphic state. Thus, indirect routes to M correspond to A > P-SM > M, whereas indirect routes to P-SD correspond to A > P-SM > P-SD. The “>” symbol indicates the direction of a transition from one state (left of symbol) to the next state (right of symbol). A dash (—) indicates that a route is not possible under the modeled state space for that component. Femoral enlargement was excluded because it was coded as a binary character. Abbreviations: A, absence of a component in both sexes; M, male-limited component expression; P-SD, sexually dimorphic expression of a component present in both sexes; P-SM, sexually monomorphic expression of a component present in both sexes.

| **Component** | **Direct A > M** | **Direct A > P-SD** | **Indirect A > M** | **Indirect A > P-SD** |
|---|---|---|---|---|
| Metapleural process | 1 | 2 | 0 | 0 |
| Coxal processes | 5 | 1 | 0 | 0 |
| Trochanteral processes | 4 | — | 0 | — |
| Dorsal femoral processes | 3 | 6 | 1 | 1 |
| Lateral femoral processes | 0 | 14 | 0 | 0 |
| Ventral femoral processes | 1 | 0 | 0 | 14 |
| Femoral curvature | 3 | — | 0 | — |
| Tibial curvature | 6 | — | 0 | — |
| Internal tibial processes | 5 | 0 | 0 | 0 |
| External tibial processes | 5 | 16 | 0 | 0 |
| Internal tibial flag | 2 | 0 | 0 | 3 |
| External tibial flag | — | 5 | — | 5 |
| Ventral abdominal processes | 2 | 0 | 0 | 0 |

Lateral femoral processes were the exception to the general congruence among phylogenetic hypotheses. Under the two ML topologies, sexual dimorphism arose predominantly through direct routes to a component already distinctly dimorphic in both sexes (medians = 14 and 13.5 routes), with essentially no stepwise routes inferred. Under the MSC topologies, however, direct routes had a median of 0, whereas the route from absence through apparent monomorphism to distinct sexual dimorphism had medians of 6 and 8. Thus, although sexual dimorphism in lateral femoral processes was repeatedly inferred to evolve, whether those origins were predominantly direct or stepwise depended strongly on phylogenetic topology.

We additionally quantified overall sexual dimorphism within taxa by calculating a sexual dimorphism score for each species based on qualitative assessments across weapon components (Table S6). Across the 154 species, sexual dimorphism scores ranged from 0 (completely sexually monomorphic) to 14 (highly sexually dimorphic), reflecting substantial variation in the extent of sexual dimorphism across Coreoidea. A few differed markedly in their sexual dimorphism score with their close relatives (e.g., 7–12 within *Acanthocephala*; 0–8 within *Leptoscelis*; 8–12 within *Mictis*), but relative homogeneity was present in the degree of sexual dimorphism for most other groups.

When grouped into discrete categories, species were distributed across all four sexual dimorphism classes: sexually monomorphic, limited sexual dimorphism, moderate sexual dimorphism, and extreme sexual dimorphism. Mapping these categories onto the ML 50p phylogeny revealed a non-random distribution of overall sexual dimorphism within Coreoidea (Fig. 7). Lineages with moderate to extreme sexual dimorphism were concentrated in specific clades: Acanthocephalini, Acanthocorini, Acanthocerini, Mictini, Nematopodini, and most of *Leptoscelis*. All other parts of the phylogeny were dominated by species exhibiting little or no sexual dimorphism across components.

**Figure 7.**
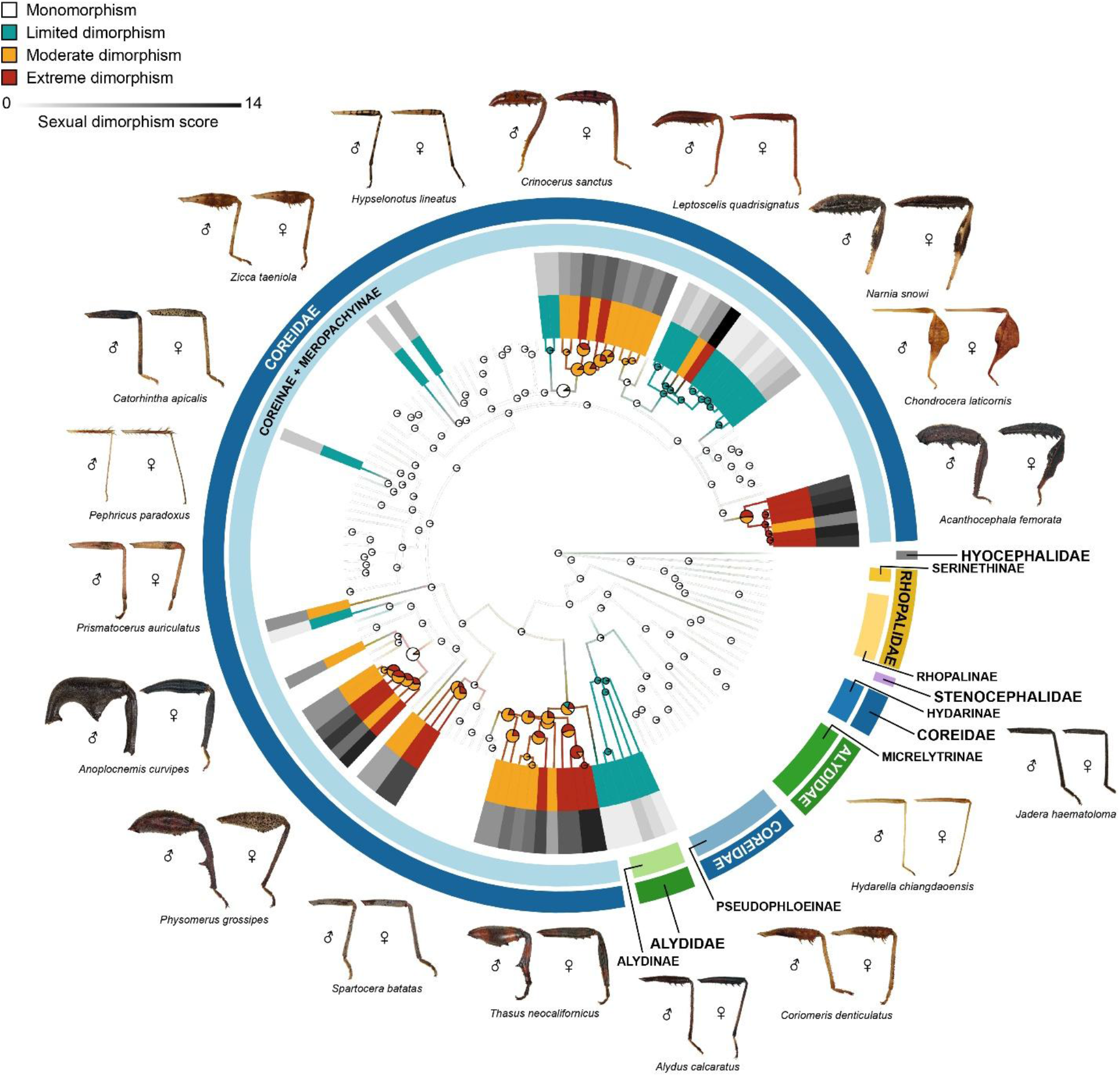
Stochastic character mapping of overall degree of sexual dimorphism on the ML 50p phylogeny. The inner terminal bars show the discrete overall sexual dimorphism category assigned to each species (monomorphic, limited, moderate, or extreme), whereas the outer grayscale terminal bars depict the underlying sexual dimorphism score for each species, scaled from low to high values. Branch colors summarize model-averaged stochastic maps, illustrating inferred histories of transitions among sexual dimorphism categories along the tree. Pie charts at internal nodes represent posterior probabilities of each sexual dimorphism category, with the radius reflecting node certainty: nodes with < 0.90 posterior support for any single state are larger. Taxon labels and outgroup taxa are omitted for visualization.

Ancestral state estimation of the degree of sexual dimorphism under the ML 50p phylogeny recovered multiple transitions among sexual dimorphism categories across the tree (Fig. 7; Table S9). Transitions from sexually monomorphic ancestors most often led to lineages with limited or moderate sexual dimorphism (median = 9 transitions each), whereas transitions from monomorphic to extreme sexual dimorphism were not inferred under the ML 50p topology. Transitions among non-monomorphic states were also common, with the most frequent involving shifts from moderate to extreme sexual dimorphism (median = 17 transitions) and vice versa (median = 12 transitions). In contrast, transitions from non-monomorphic states to complete sexual monomorphism were rare (median = 1 transition from moderate sexual dimorphism to sexually monomorphic). Thus, evolutionary changes in overall sexual dimorphism most often involved shifts in degree rather than complete gains or losses of sexual dimorphism across the entire weapon system.

## Discussion

### Evolutionary dynamics of sexual dimorphism in complex weapon systems

Our analyses revealed that sexual dimorphism in leaf-footed bug weapons is pervasive, heterogeneous, and evolutionarily dynamic. The most frequently inferred pathway involved the origin of a new component in both sexes and already in a sexually dimorphic form—a pathway rarely inferred in macroevolutionary studies of weapons (Table 1). Male-limited components also originated frequently, whereas sexual dimorphism arose less often through subsequent divergence or female loss from an ancestrally shared, monomorphic condition. Examination of complete evolutionary routes further showed that direct origins of sexual dimorphism were common for many components, although others evolved predominantly or partly through an intermediate monomorphic condition. By evaluating weapon components individually using a multistate framework, we were able to distinguish these pathways, which would likely be obscured by binary coding or continuous metrics of sex differences. At the scale of the entire weapon system, the overall magnitude of sexual dimorphism was phylogenetically clustered and generally changed in degree. Monomorphic ancestors most often gave rise to limited or moderate dimorphism, whereas extreme dimorphism arose primarily through changes across multiple components in lineages that were already dimorphic. Together, these results indicate that weapon morphology is not and tightly coupled to overall body size—with females generally larger despite predominantly male-biased weapon exaggeration—the weapon system does not evolve as a single, tightly integrated unit. Instead, its components follow partly independent trajectories whose combined changes produce substantial differences among sexes and lineages.

Consistent with our predictions, weapon components underwent frequent gains and losses, and sexual dimorphism evolved through multiple pathways. Components differed markedly in their transitions among absence, shared monomorphic expression, male-limited expression, and shared dimorphic expression. The ordered transition histories further showed that these differences extended to the complete evolutionary routes by which sexual dimorphism originated. For example, coxal and trochanteral processes most often became sexually dimorphic through a direct route to male-limited expression, whereas external tibial processes most often originated directly in both sexes in a sexually dimorphic form. In contrast, ventral femoral processes usually arose first in both sexes with apparently monomorphic expression before subsequently becoming sexually dimorphic, while other components evolved through both direct and stepwise routes. Sexual dimorphism was also lost through multiple routes; existing components sometimes returned to similar expression in both sexes, whereas in other cases the component disappeared from both sexes. Comparable gains, losses, and reversals of sexual dimorphism have been documented in, for example, beetle horns, sepsid-fly foreleg armature, harvestman leg structures, and chameleon weapon components (Emlen et al. 2005b,a; Puniamoorthy et al. 2008; Buzatto et al. 2014; Van Kleeck-Hann and Wiens 2023) (also see Table 1 for more examples). Thus, we recovered the broader set of pathways predicted, however, the frequencies of these transitions varied widely among components.

The frequent origin of components already exhibiting sexual dimorphism raises questions about the mechanisms underlying these apparently abrupt, direct evolutionary transition in our comparative analyses. Given our phylogeny, character data, and fitted evolutionary models, these changes were often reconstructed in the stochastic character maps as direct transitions from absence or a straight state to a sexually dimorphic state. Examination of the ordered transition histories likewise showed that, for many components, direct routes to sexual dimorphism were more frequent than routes passing through an intermediate state in which the component was shared and apparently monomorphic between the sexes. However, this does not rule out the possibility that some lineages passed rapidly through a short-lived intermediate condition that left little evolutionary signal. Thus, although sexual dimorphism evolved through several pathways, some components appear to have acquired sex differences at or very near the time they first originated.

This pattern raises questions about how sexual dimorphism could arise so early in the evolutionary history of a new component. One possibility is that the developmental capacity to produce certain traits exists latently even when a trait is unexpressed. Activating the growth of a trait may require only relatively small shifts in developmental regulatory pathways, such as modifications to juvenile hormone or insulin signaling (Emlen et al. 2005a; Stillwell et al. 2010). Theoretical models and empirical evidence suggest that a new trait can be dimorphic at inception via sex-specific development that is controlled by simple and evolutionarily labile regulatory mechanisms (Bonduriansky 2011). Additionally, males and females can differ in aspects of their genetic architecture, and the same regulatory change may produce different phenotypes for a given trait in each sex, even instantly when a trait appears (Williams et al. 2008; Kijimoto et al. 2012; van der Bijl and Mank 2021). These developmental and genetic mechanisms may also interact. For example, when a hormonal change activates latent trait growth, it acts upon an already sex-specific genetic architecture. The development of exaggerated beetle horns provides an example of this. In these holometabolous insects, minor endocrine shifts interact with genetic “switches” to regulate imaginal disc proliferation during metamorphosis and produce divergent, sex-specific weapons from the same tissue (Emlen et al. 2005a; Kijimoto et al. 2012). While leaf-footed bugs are hemimetabolous (i.e., adult traits develop gradually across nymphal instars), the broader principle that regulatory changes can interact with sex-specific genetic architecture to generate sexual dimorphism in a new trait is plausible and warrants further investigation.

Even though individual traits may follow different evolutionary trajectories, broader macroevolutionary patterns were evident when sexual dimorphism is considered at the level of the entire weapon system. The overall degree of sexual dimorphism across the weapon system clustered phylogenetically, with lineages characterized by highly elaborated male weapons exhibiting the strongest dimorphism in their weapons. In these clades, extreme sexual dimorphism did not occur through the isolated exaggeration of a single trait, but rather with gradual and coordinated modifications across multiple traits, such as simultaneous femoral enlargement and exaggeration of femoral and tibial processes. Because different lineages reached high levels of sexual dimorphism through different combinations of exaggerated traits, our findings demonstrate that complex weapon systems diversify through uneven changes across traits over evolutionary time.

Our prediction that patterns of sexual dimorphism would vary among weapon components was supported: individual components followed different evolutionary trajectories in the gains and losses of components and sexual dimorphism, with some showing more gains in males and others showing similar or greater gains in females. These differences do not necessarily imply that the components evolved independently from one another. For example, the various structures and components of the hind legs function collectively during male–male combat to grasp, squeeze, or strike rivals (Mitchell 1980; Eberhard 1998; Tatarnic and Spence 2013; Emberts et al. 2021), and previous work indicates that many of these components are evolutionarily correlated (Miller et al. 2024). Together, these findings suggest that weapon-system components are only partly integrated: some evolve together, whereas others respond more independently, and the degree of integration may vary among lineages. If the hind leg weapon system were shaped primarily by a single function, such as male–male combat, stronger integration among components might be expected in at least one sex. However, the same structures may also be involved in locomotion, signaling, defense, feeding-site competition, oviposition-site competition, and mate rejection, exposing them to multiple selective pressures that may reinforce integration among some components while weakening it among others (Miller et al. 2026). Thus, this partial integration may allow individual traits to respond to selection semi-independently, shaping the variety of weapon configurations and sex differences seen across lineages (Emlen et al. 2005b,a; Miller et al. 2024, 2026). Consequently, treating multicomponent weapons as single functional units may obscure meaningful variation in how sexual dimorphism arises and evolves among components within the system in respose to different selective pressures (Miller et al. 2024).

Our results suggest that body size and weapon morphology are not tightly linked across leaf-footed bugs. Across the species we examined, females were generally larger overall, a pattern commonly attributed to fecundity selection favoring increased reproductive output (Andersson 1994; Fairbairn 2013) and supported in leaf-footed bugs by positive associations between female body size and reproductive output (Nishida 1994; Miller et al. 2013). Conversely, other areas of the body were overwhelmingly exaggerated in males, consistent with expectations of strong sexual selection on structures used in male–male competition (Darwin 1871; Emlen 2008; McCullough et al. 2016). Experimental and behavioral studies have shown that variation in the size and shape of such traits influences fighting performance and contest outcomes in several species of leaf-footed bugs (Miller and Emlen 2010; Emberts et al. 2021; Graham et al. 2022).

The presence of elaborated legs in females and, in some species, sexual monomorphism of traits highlights that weapons are likely subject to selective pressures beyond male–male competition. For example, the weaponized hind legs of leaf-footed bugs are multifunctional; they are used in locomotion, aggressive interactions over mates or resources, and potentially predator defense or avoidance (Goodchild 1977; Eberhard 1998; Emberts and Wiens 2021; Evans-Blake et al. 2026). This multifunctionality of the hind legs may constrain the evolution of extreme exaggeration in some traits for both sexes while allowing others to respond more freely to sexual selection. One exception to male-biased weapon exaggeration is the width of the tibial flags, which are often monomorphic or even larger in females and likely reflect different selection pressures. Monomorphic tibial flags may have been shaped primarily by natural selection, with experimental work on *Anisoscelis alipes* (Guérin-Méneville 1833), suggesting that monomorphic flags likely function in anti-predator defense or avoidance rather than as combat weapons (Evans-Blake et al. 2026). In contrast, female-biased tibial flag exaggeration appears more consistent with social selection (West-Eberhard 1979; Tobias et al. 2012). In *Acanthocephala declivis*, females possess highly prominent tibial flags that are much smaller in males, and behavioral observations show that females use these traits as aggressive signals and during contests with rival females over access to feeding sites (Eberhard 1998). Taken together, these results suggest that sexually dimorphic weapon systems reflect the combined effects of natural and sexual selection. Clarifying the relative importance of these forces will require comparative behavioral and experimental work across species that differ in the sexual differences of the tibial flags and other weapon traits.

Taken together, our findings show that sexual dimorphism in leaf-footed bug weaponry arose through different evolutionary routes across individual traits within a partially integrated system. A multi-trait weapon system does not evolve as a single, unified structure but instead changes unevenly across traits, sexes, and lineages. This combination of partial integration and evolutionary flexibility helps explain both the striking diversity of weapon configurations among leaf-footed bugs and the varied routes through which sex differences appear and disappear. More broadly, these results highlight the importance of examining complex weapon systems at the level of individual traits when investigating the evolution of sexual dimorphism.

## Conclusions

Our study demonstrates that sexual dimorphism in highly complex animal weapons is extremely dynamic and evolutionarily labile. By investigating the complex multi-trait weapon system of leaf-footed bugs, we show that sexually dimorphic structures do not evolve as a single, integrated unit. Instead, complex weapon systems diversify through a mix of semi-independent changes across different structures. This likely allows individual traits to respond to different selective pressures, generating a wide variety of weapon configurations and sex differences across lineages. Notably, we found that the most common evolutionary pathway to sexual dimorphism in leaf-footed bug weaponry involved the origin of new traits in both sexes that were already dimorphic, rather than through divergence from ancestrally monomorphic traits. This pattern suggests that pre-existing developmental mechanisms might be responsible for the expression of new, sexually dimorphic traits without requiring intermediate monomorphic stages.

At broader macroevolutionary scales, the overall magnitude of sexual dimorphism in weapon systems is phylogenetically structured, with some clades exhibiting hotspots of extreme weapon elaboration and dimorphism. Extreme sexual dimorphism does not arise from the exaggeration of a single trait, but through coordinated elaborations across multiple traits. Furthermore, males typically show greater exaggeration consistent with sexual selection, while females are generally larger-bodied and express reduced or monomorphic traits. These patterns indicate that sexually dimorphic weapon systems may reflect the combined effects of natural, social, and sexual selection, rather than male-male competition alone.

Thus, our study points to a few areas where additional work is needed to understand the evolution of complex weapon systems, not only in in leaf-footed bugs but also other groups. First, the frequent origin of traits in an already sexually dimorphic state highlights a need for studies on the developmental and genetic mechanisms underlying these patterns in leaf-footed bugs and other insects (especially non-holometabolous insects), including whether shifts in hormonal regulation are involved and interact with existing sex-specific genetic architectures to produce dimorphic traits. Second, future work should examine how weapon traits covary and are integrated within and between sexes. Comparing patterns of trait integration in males and females may give insights as to whether different selective pressures might be influencing weapon evolution in each sex, while intersexual correlations can indicate how shared genetic architecture constrains or facilitates the evolution of sexual dimorphism. Finally, distinguishing the roles of natural, social, and sexual selection in shaping these weapon systems will require comparative behavioral and experimental work across species, with particular attention to the largely understudied aggressive and anti-predator roles of female weaponry.

## Supporting information

Supplementary methods and results (Molecular data collection and phylogenetic inferences) and supplementary figures S1-S17

Tree files and character matrix.

## Acknowledgments

We are grateful to Christiane Weirauch, Dimitri Forero, Paul Masonick, Joe Eger, Petr Kment, Marcos Roca-Cusachs, Vasily Grebennikov, Takahisa Miyatake, Nik Tatarnic, Jason Cryan, Mark Deyrup, Wei Song Hwang, Li You, Oliver Keller, John Leavengood, Wanessa da Silva Costa, Sam Noble Oklahoma Museum of Natural History, Field Museum of Natural History, Florida State Collection of Arthropods, California Academy of Sciences, and the National Museums of Kenya for providing specimens. We also thank Kevin Johnson for generously providing whole genome sequence data for *Maevius indeco*rus (NCBI SRA accession SRR6413622) through the Hemipteroid Insect Assembling the Tree of Life Project. We thank Bob McCleery, Cebisile N. Magagula, and the Savannah Research Center for assistance with specimen collection in Eswatini. South African specimens were collected under Ezemvelo KZN Wildlife permit #OP 172/2018 and with permission from the iSimangaliso Wetland Park Authority (World Heritage Site). Additional samples were supported by the University of Florida Research Abroad for Doctoral Students program, a Lee Kong Chian Natural History Museum fellowship, and National Science Foundation (NSF) grant OISE-1614015 to Zach Emberts (Australia permit #01-000204-1; Singapore permit #NP/RP17-012); the Society for the Study of Evolution Rosemary Grant Award, Smithsonian Tropical Research Institute Short Term Fellowship, and Systematics, Evolution, and Biodiversity Endowment Award (Entomological Society of America) to Ummat Somjee; and NSF DEB-0542864 to Michael Sharkey and Brian Brown. Nathan Friedman, Caroline Miller, and Min Zhao assisted with molecular laboratory work. The broader work in this manuscript was supported by the National Science Foundation under Grants IOS-2226881 and IOS-1553100 to C.W.M.

## Supplementary methods and results

Molecular data collection and phylogenetic inferences.

## Supplementary files

**File 1.** Maximum likelihood best tree based on the 50p dataset.

**File 2.** Maximum likelihood best tree based on the 50p25mi dataset.

**File 3.** Multispecies summary coalescent species tree based on the 50p dataset.

**File 4.** Multispecies summary coalescent species tree based on the 50p dataset.

**File 5.** Sexual dimorphism character matrix for traits examined in this study.

## Supplementary figures

**Figure S1.** Additional hind-leg weapon properties show both size-dependent and relatively consistent patterns of sexual dimorphism. Sex-specific allometric relationships are shown for pGLMM-analyzed components not included in Figure 3: (A) dorsal femoral width, (B) lateral femoral width, (C) dorsal femoral process length, (D) lateral femoral process length, (E) external tibial process length, (F) internal tibial flag length, and (G) external tibial flag length. Dorsal and lateral femoral width, dorsal femoral process length, and external tibial process length generally became increasingly male-biased with increasing body size. Lateral femoral process length remained male-biased across the observed body-size range despite showing sex-specific allometry, whereas internal and external tibial flag lengths showed relatively consistent male-biased differences across body sizes. Points represent log-transformed component values plotted against log-transformed pronotal width for individual specimens, colored by species. Circles and triangles denote male and female specimens, respectively. Solid and dashed lines indicate male and female trends, respectively. Shaded regions represent 95% confidence intervals (CI), bounded by solid lines for males and dashed lines for females. Number of species (NS) is indicated in each plot. For these and all other components not illustrated here, measurements and sample sizes are provided in Table S5, and full pGLMM results are provided in Table S7.

**Figure S2.** Rare weapon components in Pephricus paradoxus show little evidence of sexual dimorphism. Species-specific allometric relationships are shown for (A) coxal process length, (B) trochanteral process length, and (C) ventral abdominal process length. Points represent log-transformed component values plotted against log-transformed pronotal width for individual specimens. Circles and triangles denote male and female specimens, respectively. Solid and dashed lines indicate male and female trends, respectively. Shaded regions represent 95% CI, bounded by solid lines for males and dashed lines for females. The number of male (NM) and female (NF) specimens are indicated in each plot. Measurements and sample sizes are provided in Table S5, and full GLM results are provided in Table S8.

**Figure S3.** Stochastic character mapping of the metapleural process on the (A) ML 50p, (B) ML 50p25mi, (C) MSC 50p, and (D) MSC 50p25mi phylogenies. The terminal bars show the character state assigned to each species. Branch colors summarize model-averaged stochastic maps, illustrating inferred histories of transitions among character states. Pie charts at internal nodes represent posterior probabilities of each character state, with the radius reflecting node certainty: nodes with < 0.90 posterior support for the predominant state are larger. Outgroup taxa are omitted for visualization.

**Figure S4.** Stochastic character mapping of the coxal processes on the (A) ML 50p, (B) ML 50p25mi, (C) MSC 50p, and (D) MSC 50p25mi phylogenies. The terminal bars show the character state assigned to each species. Branch colors summarize model-averaged stochastic maps, illustrating inferred histories of transitions among character states. Pie charts at internal nodes represent posterior probabilities of each character state, with the radius reflecting node certainty: nodes with < 0.90 posterior support for the predominant state are larger. Outgroup taxa are omitted for visualization.

**Figure S5.** Stochastic character mapping of the trochanteral processes on the (A) ML 50p, (B) ML 50p25mi, (C) MSC 50p, and (D) MSC 50p25mi phylogenies. The terminal bars show the character state assigned to each species. Branch colors summarize model-averaged stochastic maps, illustrating inferred histories of transitions among character states. Pie charts at internal nodes represent posterior probabilities of each character state, with the radius reflecting node certainty: nodes with < 0.90 posterior support for the predominant state are larger. Outgroup taxa are omitted for visualization.

**Figure S6.** Stochastic character mapping of the femoral enlargement on the (A) ML 50p, (B) ML 50p25mi, (C) MSC 50p, and (D) MSC 50p25mi phylogenies. The terminal bars show the character state assigned to each species. Branch colors summarize model-averaged stochastic maps, illustrating inferred histories of transitions among character states. Pie charts at internal nodes represent posterior probabilities of each character state, with the radius reflecting node certainty: nodes with < 0.90 posterior support for the predominant state are larger. Outgroup taxa are omitted for visualization.

**Figure S7.** Stochastic character mapping of the dorsal femoral processes on the (A) ML 50p, (B) ML 50p25mi, (C) MSC 50p, and (D) MSC 50p25mi phylogenies. The terminal bars show the character state assigned to each species. Branch colors summarize model-averaged stochastic maps, illustrating inferred histories of transitions among character states. Pie charts at internal nodes represent posterior probabilities of each character state, with the radius reflecting node certainty: nodes with < 0.90 posterior support for the predominant state are larger. Outgroup taxa are omitted for visualization.

**Figure S8.** Stochastic character mapping of the lateral femoral processes on the (A) ML 50p, (B) ML 50p25mi, (C) MSC 50p, and (D) MSC 50p25mi phylogenies. The terminal bars show the character state assigned to each species. Branch colors summarize model-averaged stochastic maps, illustrating inferred histories of transitions among character states. Pie charts at internal nodes represent posterior probabilities of each character state, with the radius reflecting node certainty: nodes with < 0.90 posterior support for the predominant state are larger. Outgroup taxa are omitted for visualization.

**Figure S9.** Stochastic character mapping of the ventral femoral processes on the (A) ML 50p, (B) ML 50p25mi, (C) MSC 50p, and (D) MSC 50p25mi phylogenies. The terminal bars show the character state assigned to each species. Branch colors summarize model-averaged stochastic maps, illustrating inferred histories of transitions among character states. Pie charts at internal nodes represent posterior probabilities of each character state, with the radius reflecting node certainty: nodes with < 0.90 posterior support for the predominant state are larger. Outgroup taxa are omitted for visualization.

**Figure S10.** Stochastic character mapping of the femoral curvature on the (A) ML 50p, (B) ML 50p25mi, (C) MSC 50p, and (D) MSC 50p25mi phylogenies. The terminal bars show the character state assigned to each species. Branch colors summarize model-averaged stochastic maps, illustrating inferred histories of transitions among character states. Pie charts at internal nodes represent posterior probabilities of each character state, with the radius reflecting node certainty: nodes with < 0.90 posterior support for the predominant state are larger. Outgroup taxa are omitted for visualization.

**Figure S11.** Stochastic character mapping of the tibial curvature on the (A) ML 50p, (B) ML 50p25mi, (C) MSC 50p, and (D) MSC 50p25mi phylogenies. The terminal bars show the character state assigned to each species. Branch colors summarize model-averaged stochastic maps, illustrating inferred histories of transitions among character states. Pie charts at internal nodes represent posterior probabilities of each character state, with the radius reflecting node certainty: nodes with < 0.90 posterior support for the predominant state are larger. Outgroup taxa are omitted for visualization.

**Figure S12.** Stochastic character mapping of the internal tibial processes on the (A) ML 50p, (B) ML 50p25mi, (C) MSC 50p, and (D) MSC 50p25mi phylogenies. The terminal bars show the character state assigned to each species. Branch colors summarize model-averaged stochastic maps, illustrating inferred histories of transitions among character states. Pie charts at internal nodes represent posterior probabilities of each character state, with the radius reflecting node certainty: nodes with < 0.90 posterior support for the predominant state are larger. Outgroup taxa are omitted for visualization.

**Figure S13.** Stochastic character mapping of the external tibial processes on the (A) ML 50p, (B) ML 50p25mi, (C) MSC 50p, and (D) MSC 50p25mi phylogenies. The terminal bars show the character state assigned to each species. Branch colors summarize model-averaged stochastic maps, illustrating inferred histories of transitions among character states. Pie charts at internal nodes represent posterior probabilities of each character state, with the radius reflecting node certainty: nodes with < 0.90 posterior support for the predominant state are larger. Outgroup taxa are omitted for visualization.

**Figure S14.** Stochastic character mapping of the internal tibial flag on the (A) ML 50p, (B) ML 50p25mi, (C) MSC 50p, and (D) MSC 50p25mi phylogenies. The terminal bars show the character state assigned to each species. Branch colors summarize model-averaged stochastic maps, illustrating inferred histories of transitions among character states. Pie charts at internal nodes represent posterior probabilities of each character state, with the radius reflecting node certainty: nodes with < 0.90 posterior support for the predominant state are larger. Outgroup taxa are omitted for visualization.

**Figure S15.** Stochastic character mapping of the external tibial flag on the (A) ML 50p, (B) ML 50p25mi, (C) MSC 50p, and (D) MSC 50p25mi phylogenies. The terminal bars show the character state assigned to each species. Branch colors summarize model-averaged stochastic maps, illustrating inferred histories of transitions among character states. Pie charts at internal nodes represent posterior probabilities of each character state, with the radius reflecting node certainty: nodes with < 0.90 posterior support for the predominant state are larger. Outgroup taxa are omitted for visualization.

**Figure S16.** Stochastic character mapping of the ventral abdominal processes on the (A) ML 50p, (B) ML 50p25mi, (C) MSC 50p, and (D) MSC 50p25mi phylogenies. The terminal bars show the character state assigned to each species. Branch colors summarize model-averaged stochastic maps, illustrating inferred histories of transitions among character states. Pie charts at internal nodes represent posterior probabilities of each character state, with the radius reflecting node certainty: nodes with < 0.90 posterior support for the predominant state are larger. Outgroup taxa are omitted for visualization.

**Figure S17.** Stochastic character mapping of the overall degree of sexual dimorphism on the (A) ML 50p, (B) ML 50p25mi, (C) MSC 50p, and (D) MSC 50p25mi phylogenies. The terminal bars show the character state assigned to each species. Branch colors summarize model-averaged stochastic maps, illustrating inferred histories of transitions among character states. Pie charts at internal nodes represent posterior probabilities of each character state, with the radius reflecting node certainty: nodes with < 0.90 posterior support for the predominant state are larger. Outgroup taxa are omitted for visualization.

## Supplementary tables

**Table S1.** Summary of macroevolutionary studies of sexual dimorphism in mechanical weapon systems. Coding approach summarizes how weapon components and/or sexual dimorphism were treated in the original study. Main findings and inferred evolutionary pathways of weapon components and sexual dimorphism summarize relevant patterns, including patterns inferred here from published sex-specific reconstructions rather than only the conclusions explicitly emphasized by the original authors.

**Table S2.** Taxon sampling and NCBI Sequence Read Archive accession numbers.

**Table S3.** Summary of sequence data newly generated in the present study. Abbreviations: bp, base pairs; EtOH, ethanol; Max, maximum; Min, minimum; UCE, ultraconserved element.

**Table S4.** Summary of informative sites and number of UCE loci in datasets generated in this study.

**Table S5.** Component measurements (in mm) across individuals in both sexes of 27 species.

**Table S6.** Scores of sexual dimorphism for each weapon component and component property per taxon. The "–" symbol denotes inapplicable due to absence of component, "0" for no obvious differences between sexes, and "1" for obvious differences between sexes.

**Table S7.** Summary of phylogenetic generalized linear mixed model (pGLMM) results for weapon components across 27 species, with each phylogenetic topology tested. All measurements were log-transformed prior to analysis. Models including a phylogenetic covariance random effect are indicated with (1|Species.ID). Abbreviations: AIC, Akaike Information Criterion; logLik, log-likelihood; N, number of species; NA, not applicable; Phy, phylogenetic covariance; PW, pronotal width (proxy for body size); SE, standard error.

**Table S8.** Summary of non-phylogenetic generalized linear model (GLM) results for four weapon components observed in only one or two species. All measurements were log-transformed prior to analysis. Abbreviations: AIC, Akaike Information Criterion; logLik, log-likelihood; N, sample size; NA, not applicable; Phy, phylogenetic covariance; PW, pronotal width (proxy for body size); SE, standard error.

**Table S9.** Distributions of character state transitions inferred from stochastic character mapping across alternative phylogenetic analyses. For each weapon component and for overall degree of sexual dimorphism, the number of transitions among discrete character states (direction indicated by “->”) are summarized. Transition counts are reported as the median, mean, range (minimum–maximum), and 95% highest posterior density (HPD) interval across stochastic maps for each tree.

**Table S10.** Distributions of evolutionary routes to sexually dimorphic character states inferred from stochastic character mapping across alternative phylogenetic analyses. For each weapon component, the number of routes from the plesiomorphic condition to the first occurrence of a sexually dimorphic state are summarized. Routes were traced from the most recent occurrence of absence in both sexes or straight morphology in both sexes. A route terminated at the first occurrence of a sexually dimorphic state, defined as presence in males only or presence in both sexes with distinct sexual dimorphism. If a lineage returned to the plesiomorphic condition before reaching a sexually dimorphic state, the route was reset from the most recent occurrence of that condition. Subsequent transitions among sexually dimorphic states were not considered additional origins unless the lineage first returned to the plesiomorphic condition. Route counts are reported as the median, mean, range (minimum–maximum), and 95% highest posterior density (HPD) interval across stochastic maps for each tree. Femoral enlargement was excluded because it was coded as a binary character.

