## Supplementary methods and results (Molecular data collection and phylogenetic inferences) and supplementary figures S1-S17 for "The evolution of sexual dimorphism in the weapon system of leaf-footed bugs (Hemiptera: Coreoidea)"

*Molecular data collection*

We targeted 3040 ultraconserved element (UCE) loci across 336 ingroup and outgroup taxa. The bait set was primarily derived from two pentatomomorphan genomes (Faircloth 2017; Forthman, Miller, et al. 2019) and supplemented with additional probes published in Forthman, Gordon, et al. (2023).

Raw UCE sequence-capture data for 302 taxa were assembled from numerous prior studies (Kieran et al. 2019; Forthman, Miller, et al. 2019, 2020, 2022; Emberts et al. 2020; C. D. Miller et al. 2022; Forthman, Downie, et al. 2023; Forthman, Gordon, et al. 2023; Forthman, Phan, et al. 2024; C. W. Miller et al. 2024) (Table S2). To broaden our ingroup and outgroup representation, we also extracted UCEs from 14 taxa: two species within Coreoidea (Hyocephalidae: *Maevius indecorus* Stål, 1874; Rhophalidae: *Boisea trivittata* [Say, 1825]) and 12 additional pentatomomorphan taxa (Table S2; Sparks et al. 2020; Johnson et al. 2018; Panfilio et al. 2019).

We generated new UCE sequence data for 20 taxa. DNA extraction approaches outlined in earlier studies (Forthman, Miller et al. 2020, 2022) were used. Sixteen samples had been preserved in ethanol (EtOH), whereas the remaining four were dried, pinned museum specimens (Table S3). Depending on specimen size, genomic DNA was isolated from whole bodies or from comparable amounts of tissue (e.g., thorax, abdomen, or legs).

For EtOH-preserved samples, extractions were performed with the Qiagen DNeasy Blood and Tissue Kit (DNeasy), with the following adjustments to the manufacturer's protocol: tissues were digested for 24–48 hours in 180 µL Buffer ATL and 20 µL proteinase K, and DNA was subsequently eluted twice in 50 µL of Buffer AE. For dried specimens, we used a combined protocol (DNQIA) that pairs the DNeasy kit with a Qiagen QIAquick PCR Purification Kit (Knyshov et al. 2019; Forthman, Miller et al. 2020, 2022). This method follows the modified DNeasy procedure but substitutes a QIAquick spin column, replaces AW wash buffers with Buffer PE, and elutes DNA twice with 30 µL of Buffer EB.

We assessed DNA quality and concentration using 1% agarose gel electrophoresis and a Qubit 2.0 fluorometer. When possible, extracts were normalized to 10–20 ng/µL. High-molecular-weight DNA was sheared to approximately 200–1000 bp using a Covaris M220 Focused-ultrasonicator (20–60 s). DNA from dried specimens was additionally treated with a PreCR Repair Mix followed by a 3× SPRI bead purification.

Libraries were made following the KAPA Hyper Prep Kit protocol with several adjustments. All reactions were carried out at half the manufacturer's recommended volumes. Fragmented DNA was ligated to iTru universal adapter stubs and dual 8-bp indexes (Glenn et al. 2019). Libraries were then amplified under the following thermal profile: an initial denaturation at 98 °C for 3 minutes; 14–17 amplification cycles of 98 °C for 30 seconds, 60 °C for 30 seconds, and 72 °C for 30 seconds; and a final extension at 72 °C for 5 minutes. Following amplification, library concentration and fragment quality were assessed using Qubit and agarose gel electrophoresis. Equimolar amounts of the libraries were then combined into 1000-ng pools, dried at 60 °C, and rehydrated in 14 μL of IDTE for downstream enrichment.

Library enrichment followed one of three touchdown hybridization schemes described previously (Table S3; Forthman, Miller et al. 2020; Forthman, Gordon et al. 2023). These same hybridization regimes correspond to three separate sequencing efforts, two of which were originally used to test how hybridization temperature influences capture efficiency (Forthman, Gordon et al. 2023). For enrichment, bait volumes were adjusted by preservation type: ethanol-preserved samples received 1/2 the standard bait volume, whereas dried museum specimens received 1/4. Baits were hybridized with libraries using one of the three touchdown hybridization protocols: (1) 65°C for 18 hours followed by 62°C for 28 hours; (2) 65°C for 12 hours, 62°C for 12 hours, and 60°C for 12 hours; and (3) 65°C for 12 hours, 60°C for 12 hours, and 55°C for 12 hours. After hybridization, bait-library complexes were captured using Dynabeads M-280 Streptavidin beads, washed four times at the lowest temperature used in the respective hybridization series, and eluted in 30 μL of IDTE. Post-capture PCR was performed with iTru P5/P7 primers (2.5 μL each of 5 μM; Glenn et al. 2019) for 14–18 cycles, using an annealing temperature of 60–62 °C and a 45-second extension step. Amplified enrichments were purified with hydrophobic Sera-Mag SpeedBeads and two 70% ethanol washes. Final enriched libraries were resuspended in 22 μL IDTE, quantified using Qubit, and then pooled in equimolar amounts into a single library, which was sequenced on one Illumina HiSeq 3000 lane (2x100 bp) at the University of Florida’s Interdisciplinary Center for Biotechnology Research.

Unless otherwise noted, all subsequent data-processing steps were run using default program parameters. Adapter trimming of demultiplexed raw reads was performed with Illumiprocessor v2.0 (Faircloth 2013). Duplicate sequences were filtered out using PRINSEQ-lite v0.20.4 (Schmieder & Edwards 2011). The remaining reads were then subjected to error correction with QuorUM v1.1.0 (Marçais et al. 2015). Cleaned reads were assembled *de novo* with SPAdes v3.13.0 using the single-cell and automatic coverage cutoff settings (Nurk et al. 2013). UCE loci were identified from the resulting contigs with PHYLUCE v1.7.0 (Faircloth 2016), following workflows of Forthman, Miller, et al. (2019, 2020, 2022). For genome and transcriptome data, UCE probes were aligned to assemblies, and matched loci—including up to 500 bp of flanking sequence—were extracted through PHYLUCE. Summaries of reads, contigs, and UCE recovery for new libraries and publicly derived assemblies are reported in Table S3.

Initial alignments were generated in PHYLUCE using MAFFT (Katoh, Misawa, et al. 2002; Katoh & Standley 2013) with the --mafft, --no-trim, and --ambiguous options enabled. Visual inspection showed that a subset of sequences may have been incorrectly reverse-complemented by MAFFT’s --adjustdirection routine incorporated into the PHYLUCE mafft.py wrapper. To examine these cases systematically, we applied an approach following Schultz et al. (2024). First, we produced a second full set of alignments in which the --adjustdirection flag was deliberately disabled, while keeping all other MAFFT parameters constant. For each locus, we used an R script (plot_dists_rev.R of Schultz et al. [2024]) to quantify how each potentially reversed sequence differed from the mean pairwise distance of all sequences in the alignment under the two conditions. Distances were calculated in R v4.5.1 (R Core Team 2025) and expressed in units of standard deviation from the mean. For any sequence whose change in standardized distance was ≤ 1 SD between the "reversed" and "not reversed" versions, we manually compared the two alignments in Mesquite v3.81 (Maddison & Maddison 2023). We also confirmed signal orientation by generating dot-plot comparisons for the non-reverse-complemented alignments using the MAFFT online interface (Katoh & Standley 2013; Katoh, Rozewicki, et al. 2019). After the complete review process, 187 sequences flagged as reversed were retained as correctly oriented.

To eliminate ambiguously aligned regions, each locus was first processed with Gblocks v0.91b (Castresana 2000) using relaxed parameter settings (-b1 = 0.5, -b2 = 0.5, -b3 = 12, -b4 = 7, -b5 = h). The resulting alignments were concatenated in PHYLUCE, after which we performed an additional round of quality filtering using Spruceup v2019.2.3 (Borowiec 2019). Spruceup was ran on the concatenated alignment without supplying a guide tree, using Jukes–Cantor p-distance corrections, a window size of 20, an overlap of 15, a 0.6 fraction threshold, and the lognorm criterion. We evaluated several candidate distance thresholds (0.90, 0.95, 0.97, and 0.99) by examining the diagnostic plots generated by Spruceup. We selected a 0.95 lognorm cutoff for our dataset, which removed approximately 1.03% of the total alignment (1,882,495 bp). Following trimming, the concatenated alignment was split back into individual locus alignments using AMAS v1.0 (Borowiec 2016). Loci represented by at least 50% of sampled taxa were retained to form the "50p" dataset. From this set, we constructed an additional matrix comprising the top 25% of loci ranked by parsimony-informative sites, referred to as the "50p25mi" dataset.

*Phylogenetic inferences*

For each dataset, we combined the individual locus alignments into a concatenated alignment for maximum likelihood (ML) inference. We then applied Sliding-Window Site Characteristic Entropy (SWSC-EN) partitioning using PartitionUCE v1.0.0 (Tagliacollo & Lanfear 2018), which generated a configuration file used in IQ-Tree v2.1.3 (Minh et al. 2020) to determine the most appropriate substitution models and partitioning schemes using -m MF+MERGE (Kalyaanamoorthy et al., 2017), -rcluster 10, and -mrate E,I,G,R. We excluded I+G combinations from model selection because these parameters are statistically non-independent (Sullivan et al. 1999; Yang 2006).

Using these partitions, we performed 10 independent ML analyses for each dataset (Chernomor et al. 2016). Nodal support was quantified using 1000 Shimodaira–Hasegawa-like approximate likelihood ratio test (SH-aLRT) replicates (Guindon et al. 2010). The tree with the best log-likelihood was selected for each dataset.

To account for gene tree discordance due to incomplete lineage sorting, estimated species trees under the multispecies coalescent (MSC) using weighted ASTRAL-hybrid (wASTRAL-h), implemented in ASTER v1.22.3.7 (Zhang & Mirarab 2016; Zhang et al. 2025). Before running wASTRAL-h, each locus alignment was analyzed individually in IQ-Tree to identify its best-fitting substitution model (-mrate E,I,G,R) and to subsequently infer a gene tree (-m MFP), with branches effectively at zero length collapsed (-czb). Branch support on the final species trees was evaluated using local posterior probabilities (LPP; Sayyari & Mirarab 2016). A summary of alignment lengths, variable sites, and numbers of loci for both datasets can be found in Table S4.

To prepare our trees for ancestral state estimation (ASE), ML and MSC topologies were converted to ultrametric trees. Branch lengths on the 50p and 50p25mi MSC trees were first re-estimated in IQ-Tree in units of substitutions. We then removed all outgroup taxa and 15 ingroup taxa that displayed disproportionately long branches—these were associated with extensive missing data in degraded samples and could adversely affect time-scaling procedures. Ultrametric trees were generated in R using the *chronos* function of the *ape* package v5.8.1 (Paradis & Schliep 2019). We evaluated four molecular clock models (correlated, discrete, relaxed, and strict clock) under five smoothing parameters ($\lambda$ = 0, 0.1, 0.5, 1, 10). The optimal combination of model and smoothing value was selected by minimizing the phylogenetic information criterion (PHIIC; Paradis 2013).

### Supplementary results

#### *Phylogeny of the Coreoidea*

Our phylogenetic analyses were congruent for many recovered relationships (Fig. 2, Supplementary Files 1–4), with most nodes moderately to highly supported. Where reported, support values are given in the following order: ML 50p / ML 50p25mi / MSC 50p / MSC 50p25mi.

Across all four analyses, Coreoidea was recovered as a clade with full support, with Hyocephalidae sister to the remaining coreoids. Rhopalidae was monophyletic with high support and sister to Alydidae + Coreidae + Stenocephalidae (100 / 100 / 0.9998 / 0.9999). Within Rhopalidae, Rhopalinae and Serinethinae were each monophyletic with full support. Within Rhopalinae, Chorosomatini and Harmostini rendered Rhopalini paraphyletic, with all relationships within the subfamily highly supported (SH-aLRT = 100, LPP = 0.9155–1). Stenocephalidae was monophyletic with strong support and sister to Alydidae + Coreidae (100 / 100 / 1 / 0.976).

Although Alydidae + Coreidae formed a fully supported clade, neither family was monophyletic. Hydarinae and Pseudophloeinae—traditionally classified within Coreidae—were more closely related to Alydidae (hereafter the AHP clade). The AHP clade received full support, with Hydarinae sister to Alydidae + Pseudophloeinae. Within Alydidae + Pseudophloeinae, Micrelytrinae was monophyletic with full support and sister to Alydinae + Pseudophloeinae (97.3 / 59.6 / 0.9886 / 0.7242). Within Micrelytrinae, Leptocorisini and Micrelytrini were each monophyletic with full support. The sister-group relationship between Alydinae and Pseudophloeinae was fully supported, as were the monophyly of both subfamilies and the monophyly and sister-group relationship of Clavigrallini and Pseudophloeini within Pseudophloeinae.

The remaining two subfamilies of Coreidae—Coreinae and Meropachyinae—consistently formed a fully supported clade, but neither was monophyletic. Meropachyinae was polyphyletic and rendered Coreinae paraphyletic: Meropachyini and Spathophorini (Meropachyinae) were recovered within a clade comprising Discogastrini and Nematopodini (Coreinae) (hereafter the DMNS clade), whereas Merocorini was placed within a clade comprising Acanthocerini, Chariesterini, and Hypselonotini.

Within the Coreinae + Meropachyinae, the fully supported DMNS clade was sister to all remaining taxa. Although relationships within this clade varied, all analyses supported the non-monophyly of Discogastrini, Nematopodini, and Spathophorini. Meropachyini formed a fully supported clade, but its placement differed among analyses: it was sister to part of Spathophorini in all analyses except MSC 50p, in which it was sister to all other taxa in the clade; all alternative placements of Meropachyini received full support. When a Meropachyini + Spathophorini (part) clade was recovered, it was either sister to all remaining taxa in the DMNS clade (ML and MSC 50p25mi) or to all DMNS taxa except *Discogaster dentipes* (Stål, 1867) (Discogastrini) (ML 50p), with each topology receiving full support.

Petascelini and Spartocerini were monophyletic and sister taxa with full support. Although Acanthocorini was not monophyletic, most sampled taxa formed a fully supported clade sister to Petascelini + Spartocerini, also with full support. *Pomponatius typicus* Distant, 1904, (Acanthocorini) was consistently recovered within Amorbini with full support.

The remaining taxa within Coreinae + Meropachyinae formed four primary lineages, with relationships among these varying in topology and support across analyses : 1) Acanthocorini (part) + Agriopocorini + Amorbini + Cloresmini + Colpurini + Dasynini (part) + Mictini (hereafter Clade A), 2) Coreini + Dasynini (part) + Gonocerini + Homoeocerini + Manocoreini + Phyllomorphini + Prionotylini (Clade B), 3) Daladerini + Latimbini, and 4) Acanthocephalini + Acanthocerini + Anisoscelini + Chariesterini + Chelinideini + Hypselonotini + *Mamurius* + Merocorini + Placoscelini (Clade C). In the ML 50p and 50p25mi analyses, Clade A was sister to the remaining three clades (SH-aLRT = 100), followed by Daladerini + Latimbini sister to Clade B + Clade C (SH-aLRT = 100); Clade B and Clade C were weakly to strongly supported as sister groups (50p SH-aLRT = 99.6; 50p25mi SH-aLRT = 55.7). In the MSC 50p analysis, Clade B was fully supported as the sister to all other lineages. Daladerini + Latimbini formed a fully supported sister-group relationship with Clade A, and this combined clade was sister to Clade C with full support. In the MSC 50p25mi analysis, Clade A was sister to the remaining lineages with full support, followed by Clade B moderately supported as sister to Daladerini + Latimbini + Clade C (LPP = 0.8333). Daladerini + Latimbini and Clade C formed a fully supported sister-group relationship.

Daladerini + Latimbini was fully supported, with Latimbini rendering Daladerini paraphyletic with strong support. *Dalader planiventris* (Westwood, 1842) was more closely related to Latimbini (100 / 100 / 0.9989 / 1).

In clade A, Agriopocorini, Cloresmini, Colpurini, and Mictini were each monophyletic with full support. Cloresmini was sister to the remaining taxa in Clade A (100 / 87 / 1 / 1), followed by *Aulacosternum nigrorubrum* Dallas, 1852, (Dasynini) sister to the remaining Clade

A taxa (100 / 100 / 1 / 0.9982). Agriopocorini and Colpurini formed a highly supported sister-group relationship (100 / 100 / 1 / 0.9999), which together were sister to Amorbini + *Pomponatius* + Mictini with full support. Amorbini + *Pomponatius* was sister to Mictini with full support.

Within Clade B, only Homoeocerini and Phyllomorphini were consistently recovered as monophyletic, both with full support. Gonocerini and Coreini were polyphyletic. Homoeocerini was sister to *Gonocerus* + *Plinachtus* (Gonocerini) with full support. In most analyses, Gonocerini (part) + Homoeocerini was sister to part of Dasynini (ML 50p SH-aLRT = 98.6 / MSC 50p LPP = 1 / MSC 50p25mi LPP = 0.5417), which together were sister to Manocoreini with full support. In contrast, Gonocerini (part) + Homoeocerini were sister to Manocoreini in the ML 50p25mi analysis (SH-aLRT = 77.5), and this clade was sister to Dasynini (part) with full support. Dasynini (part) + Gonocerini (part) + Homoeocerini + Manocoreini was sister to the remaining Clade B taxa with full support. Prionotylini was fully supported as sister to *Haidara producta* Distant, 1908, (Coreini) and this pair was sister to the remaining Coreini (part) + Gonocerini (part) + Phyllomorphini with full support. Phyllomorphini were strongly supported as sister to *Spathocera dahlmanii* (Schilling, 1829) (Coreini) (100 / 100 / 0.9004 / 0.9999), and this pair was sister to the remaining Coreini (part) + Gonocerini (part) with full support. Relationships within the remaining Coreini (part) + Gonocerini (part) varied across analyses. Nevertheless, relationships involving Coreini and Gonocerini within Clade B consistently showed that both tribes are not monophyletic.

Clade C included the following monophyletic tribes, each with full support: Acanthocerini, Acanthocephalini, Chariesterini, Chelinideini, Merocorini, and Placoscelini. Anisoscelini and Hypselonotini were both polyphyletic, with Anisoscelini recovered in four and Hypselonotini in seven distinct lineages. The first lineage of Hypselonotini (*Althos* + *Catorhintha* + *Scolopocerus* + *Vazquesitocoris* + *Villasitocoris*) was sister to all remaining taxa with full support. The second lineage of Hypselonotini (*Zicca*) form a poorly supported sister-group relationship with *Dalmatomammurius vandoesburgi* (Brailovsky, 1982) (Anisoscelini) in the ML analyses (50p SH-aLRT = 57.3, 50p25mi SH-aLRT = 53.6) or with *Dalmatomammurius vandoesburgi* + the third lineage of Hypselonotini (*Acidomeria* + *Sethenira* + *Sphictyrtus*) + *Mamurius* (currently *incertae sedis* in tribal placement) in the MSC analyses (50p LPP = 0.9426, 50p25mi LPP = 0.9935). *Mamurius mopsus* Stål, 1862, was fully supported as sister to the third Hypselonotini lineage. The Anisoscelini (part) + Hypselonotini (part) + *Mamurius* clade was fully supported as sister to the Acanthocerini + Chariesterini + fourth and fifth lineages of Hypselonotini + Merocorini. The fourth lineage of Hypselonotini (*Cebrenis* + *Cebrenistella* + *Collatia* + *Petersitocoroides*) was fully supported as sister to the remaining taxa within this subclade. The fifth lineage of Hypselonotini (*Hypselonotus*) was strongly supported as sister to Chariesterini (100 / 100 / 1 / 0.9999). The Chariesterini + Hypselonotini (part) clade was fully supported as sister to a weakly to moderately supported Acanthocerini + Merocorini clade in our ML analyses (ML 50p SH-aLRT = 78.8, ML 50p25mi SH-aLRT = 58.3). In contrast, the Chariesterini + Hypselonotini (part) clade was poorly to highly supported as sister to Merocorini in our MSC analyses (MSC 50p LPP = 0.9054, MSC 50p25mi LPP = 0.6594), and this combined clade was highly supported as sister to Acanthocerini (MSC 50p LPP = 0.9993, MSC 50p25mi LPP = 0.9997). This Acanthocerini + Chariesterini + first to fifth lineages of Hypselonotini (part) + Merocorini clade was sister to all remaining Clade C taxa with full support. A second lineage of Anisoscelini (*Leptoglossus* + *Leptoscelis* + *Narnia* + *Phthia* + *Phthiacnemia* + *Phthiadema* + *Ugnius*) formed a large, fully supported clade that included *Namacus annulicornis* Stål, 1870, (sixth Hypselonotini lineage), which together was sister to the remaining taxa in Clade C with full support. The third lineage of Anisoscelini (*Diactor bilineatus* [Fabricius, 1803]) was sister to Acanthocephalini + Placoscelini in our ML analyses or to Acanthocephalini + Anisoscelini (part) + Chelinideini + seventh lineage of Hypselonotini + Placoscelini in the MSC analyses; both alternatives were fully supported. Chelinideini was fully supported as sister to a fully supported clade comprising the seventh Hypselonotini lineage (*Anasa* + *Paryphes*) and the fourth Anisoscelini lineage (*Anisoscelis* + *Chondrocera* + *Holhymenia*). The fourth Anisoscelini lineage + Chelinideini + seventh Hypselonotini lineage clade was sister to Acanthocephalini + *Diactor* + Placoscelini in our ML analyses with full support or to Acanthocephalini + Placoscelini with strong support in the MSC analyses (50p LPP = 0.9677, 50p25mi LPP = 0.9627). The sister-group relationship between Acanthocephalini and Placoscelini was fully supported across all analyses.

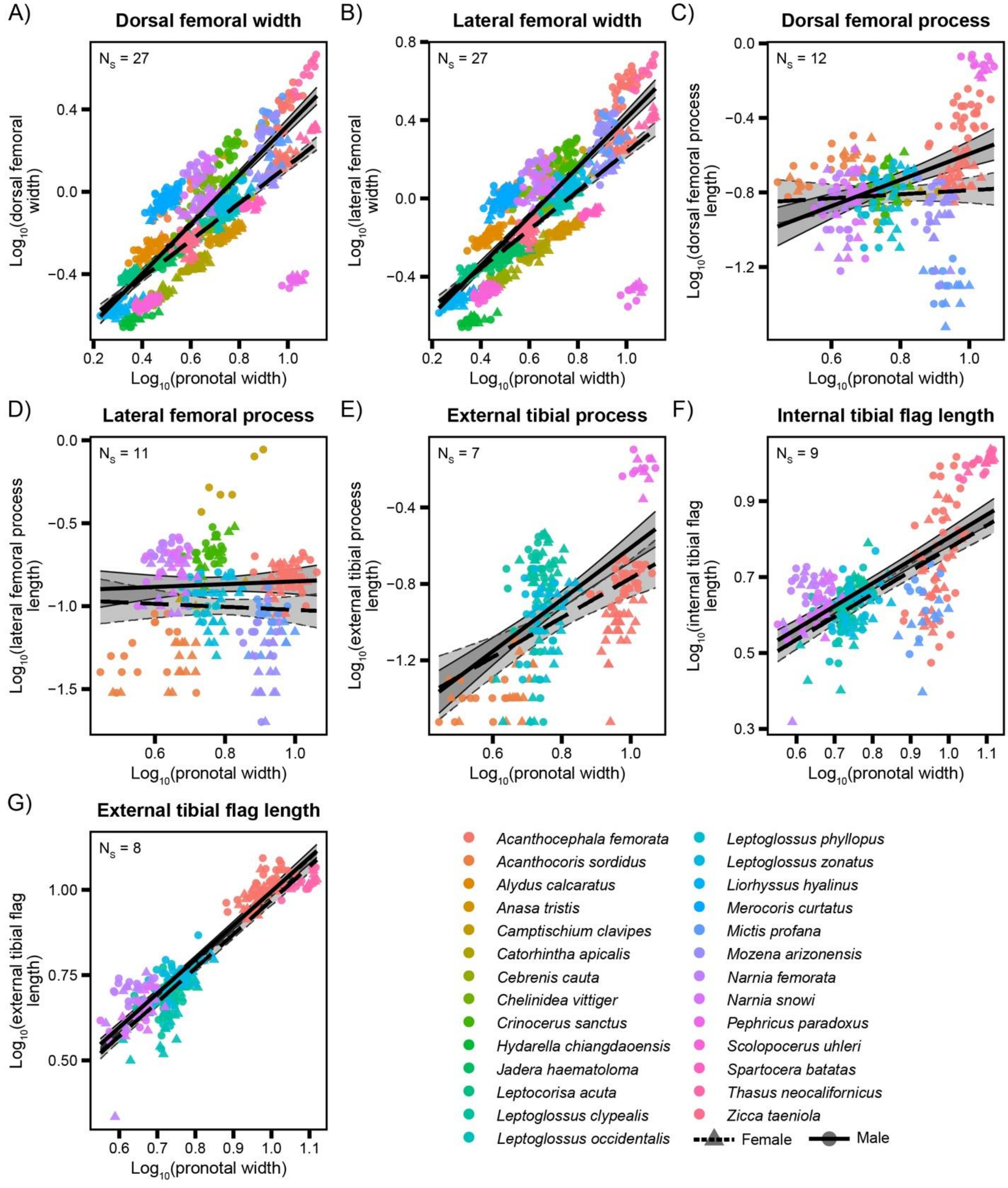


**Figure S1.** Additional hind-leg weapon properties show both size-dependent and relatively consistent patterns of sexual dimorphism. Sex-specific allometric relationships are shown for pGLMM-analyzed components not included in Figure 3: (A) dorsal femoral width, (B) lateral femoral width, (C) dorsal femoral process length, (D) lateral femoral process length, (E) external tibial process length, (F) internal tibial flag length, and (G) external tibial flag length. Dorsal and lateral femoral width, dorsal femoral process length, and external tibial process length generally became increasingly male-biased with increasing body size. Lateral femoral process length remained male-biased across the observed body-size range despite showing sex-specific allometry, whereas internal and external tibial flag lengths showed relatively consistent male-biased differences across body sizes. Points represent log-transformed component values plotted against log-transformed pronotal width for individual specimens, colored by species. Circles and triangles denote male and female specimens, respectively. Solid and dashed lines indicate male and female trends, respectively. Shaded regions represent 95% confidence intervals (CI), bounded by solid lines for males and dashed lines for females. Number of species ($N_S$) is indicated in each plot. For these and all other components not illustrated here, measurements and sample sizes are provided in Table S5, and full pGLMM results are provided in Table S7.

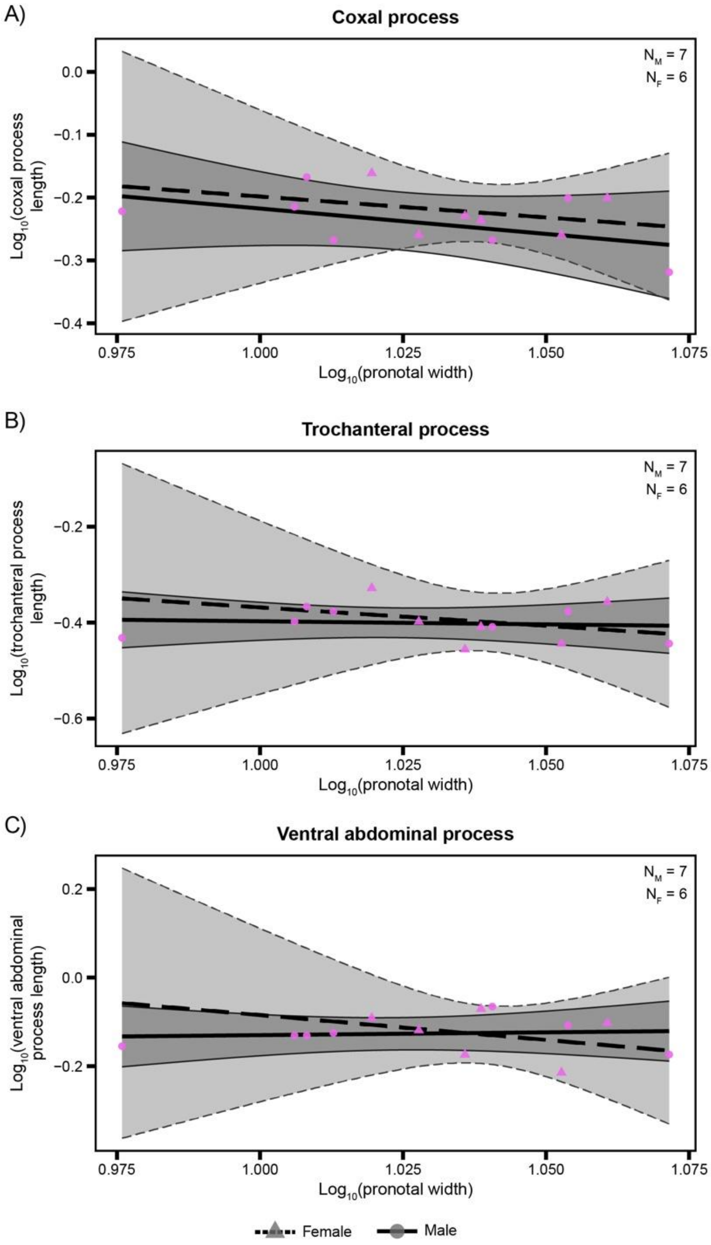


**Figure S2.** Rare weapon components in *Pephricus paradoxus* show little evidence of sexual dimorphism. Species-specific allometric relationships are shown for (A) coxal process length, (B) trochanteral process length, and (C) ventral abdominal process length. Points represent log-transformed component values plotted against log-transformed pronotal width for individual specimens. Circles and triangles denote male and female specimens, respectively. Solid and dashed lines indicate male and female trends, respectively. Shaded regions represent 95% CI, bounded by solid lines for males and dashed lines for females. The number of male ($N_M$) and female ($N_F$) specimens are indicated in each plot. Measurements and sample sizes are provided in Table S5, and full GLM results are provided in Table S8.

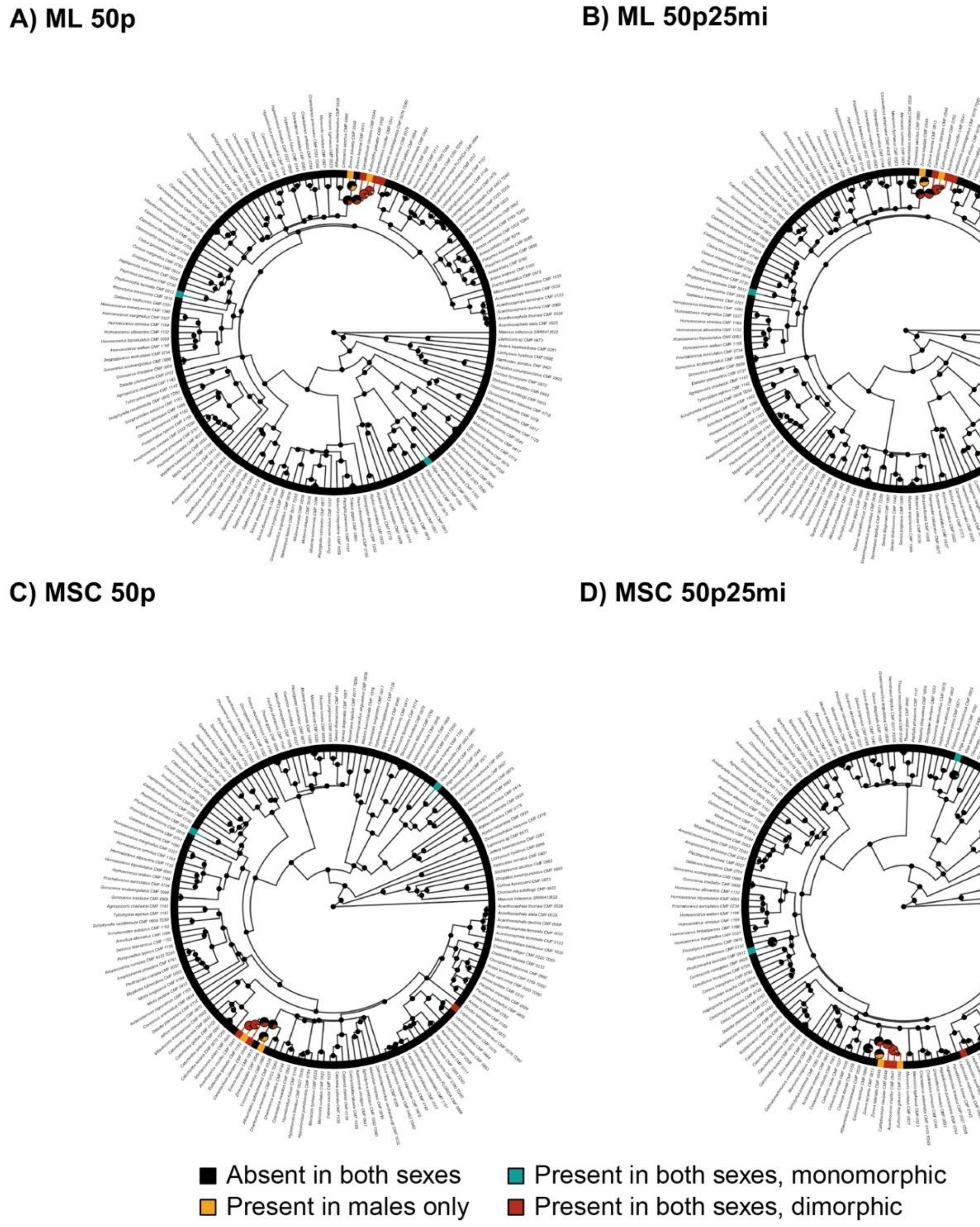


**Figure S3.** Stochastic character mapping of the metapleural process on the (A) ML 50p, (B) ML 50p25mi, (C) MSC 50p, and (D) MSC 50p25mi phylogenies. The terminal bars show the character state assigned to each species. Branch colors summarize model-averaged stochastic maps, illustrating inferred histories of transitions among character states. Pie charts at internal nodes represent posterior probabilities of each character state, with the radius reflecting node certainty: nodes with < 0.90 posterior support for the predominant state are larger. Outgroup taxa are omitted for visualization.

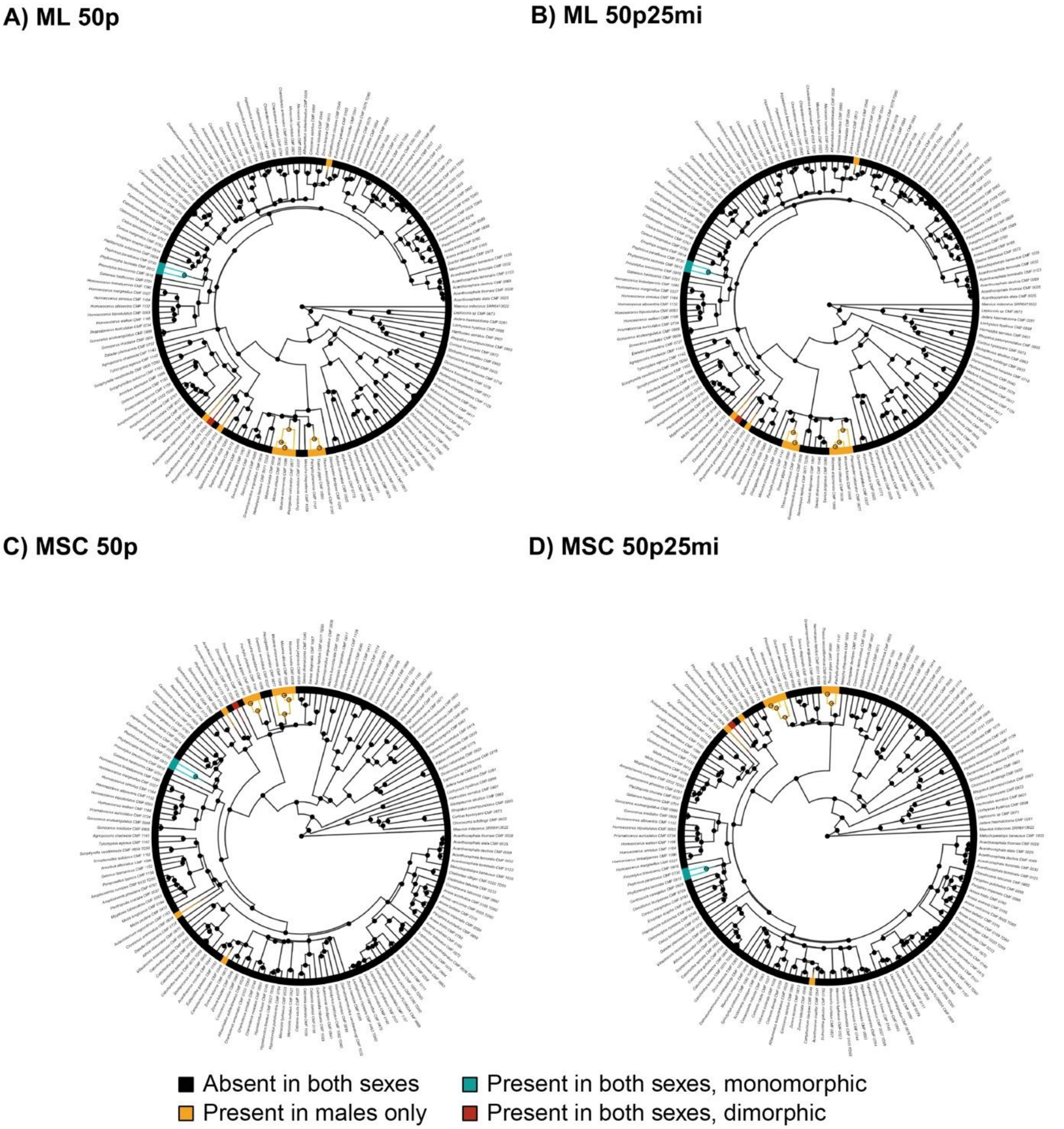


**Figure S4.** Stochastic character mapping of the coxal processes on the (A) ML 50p, (B) ML 50p25mi, (C) MSC 50p, and (D) MSC 50p25mi phylogenies. The terminal bars show the character state assigned to each species. Branch colors summarize model-averaged stochastic maps, illustrating inferred histories of transitions among character states. Pie charts at internal nodes represent posterior probabilities of each character state, with the radius reflecting node certainty: nodes with < 0.90 posterior support for the predominant state are larger. Outgroup taxa are omitted for visualization.

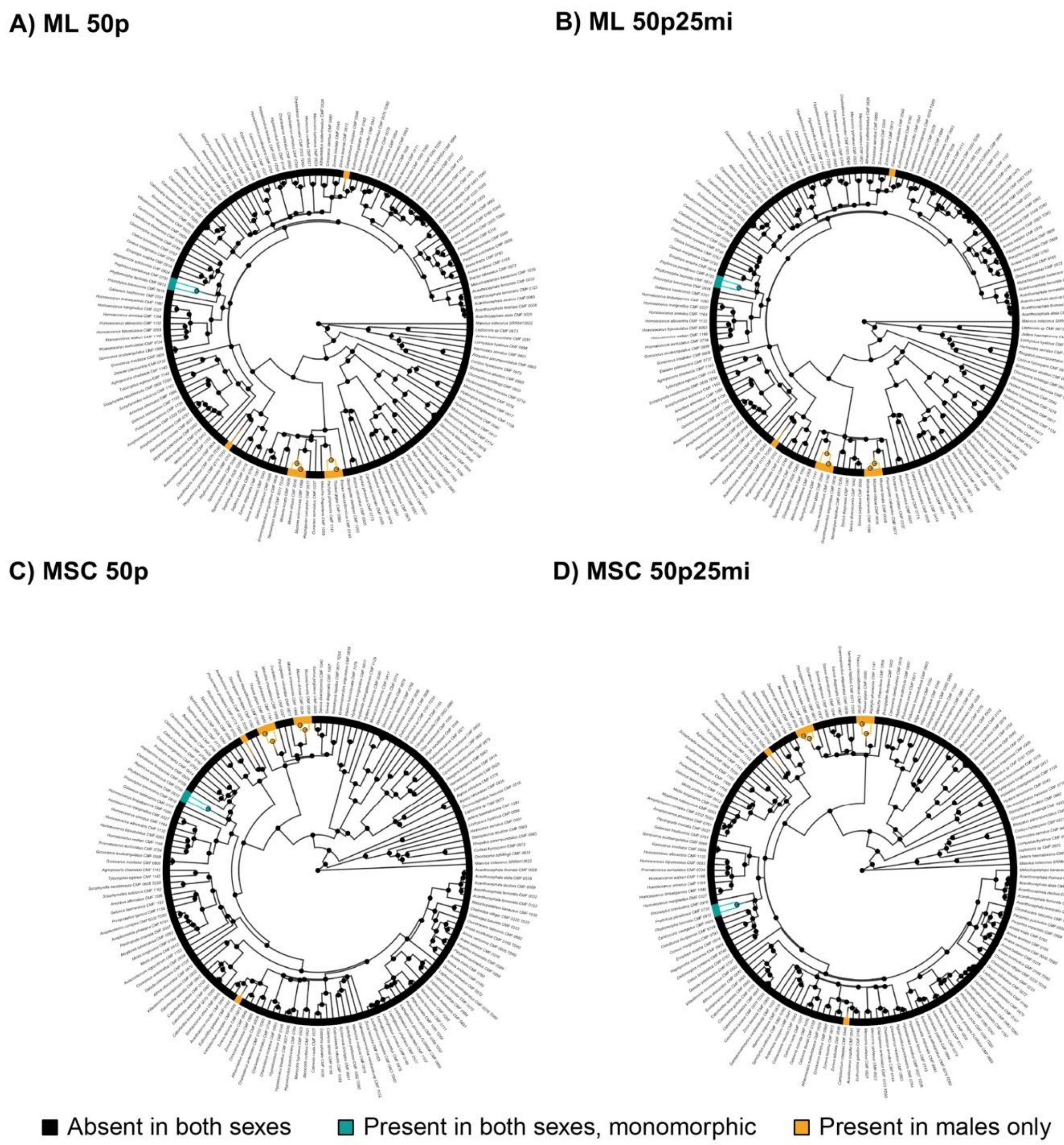


**Figure S5.** Stochastic character mapping of the trochanteral processes on the (A) ML 50p, (B) ML 50p25mi, (C) MSC 50p, and (D) MSC 50p25mi phylogenies. The terminal bars show the character state assigned to each species. Branch colors summarize model-averaged stochastic maps, illustrating inferred histories of transitions among character states. Pie charts at internal nodes represent posterior probabilities of each character state, with the radius reflecting node certainty: nodes with < 0.90 posterior support for the predominant state are larger. Outgroup taxa are omitted for visualization.

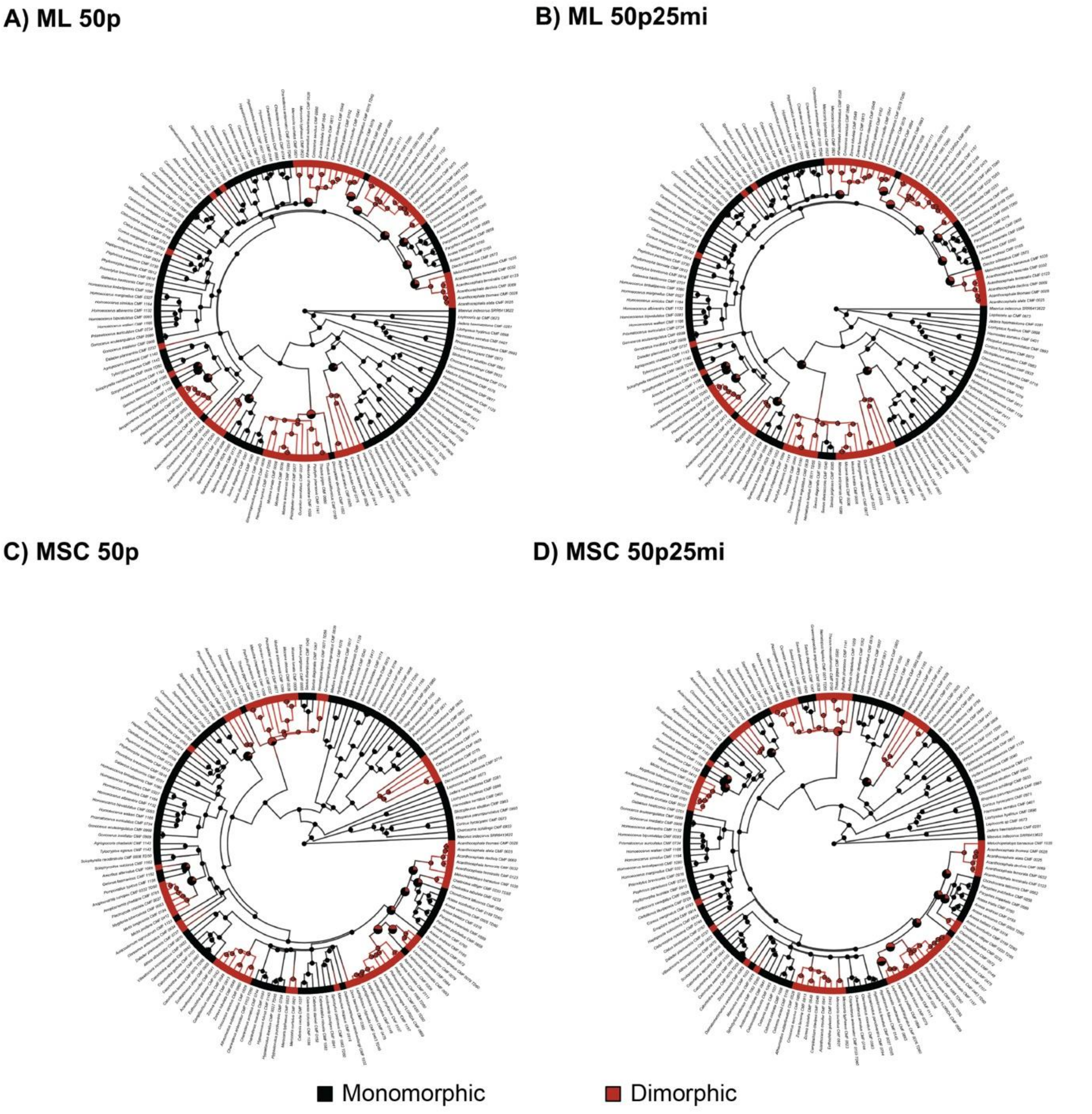


**Figure S6.** Stochastic character mapping of the femoral enlargement on the (A) ML 50p, (B) ML 50p25mi, (C) MSC 50p, and (D) MSC 50p25mi phylogenies. The terminal bars show the character state assigned to each species. Branch colors summarize model-averaged stochastic maps, illustrating inferred histories of transitions among character states. Pie charts at internal nodes represent posterior probabilities of each character state, with the radius reflecting node certainty: nodes with < 0.90 posterior support for the predominant state are larger. Outgroup taxa are omitted for visualization.

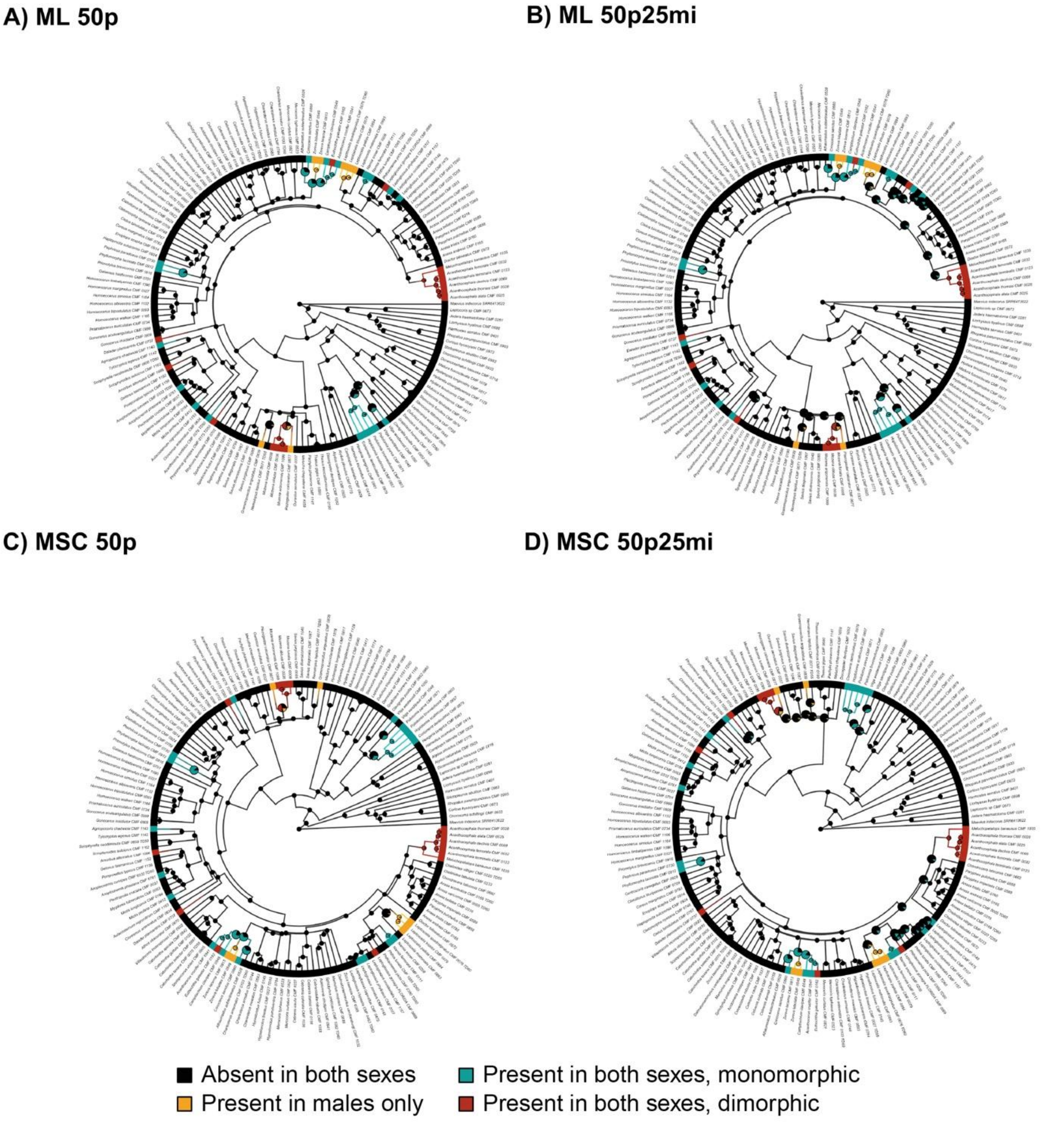


**Figure S7.** Stochastic character mapping of the dorsal femoral processes on the (A) ML 50p, (B) ML 50p25mi, (C) MSC 50p, and (D) MSC 50p25mi phylogenies. The terminal bars show the character state assigned to each species. Branch colors summarize model-averaged stochastic maps, illustrating inferred histories of transitions among character states. Pie charts at internal nodes represent posterior probabilities of each character state, with the radius reflecting node certainty: nodes with < 0.90 posterior support for the predominant state are larger. Outgroup taxa are omitted for visualization.

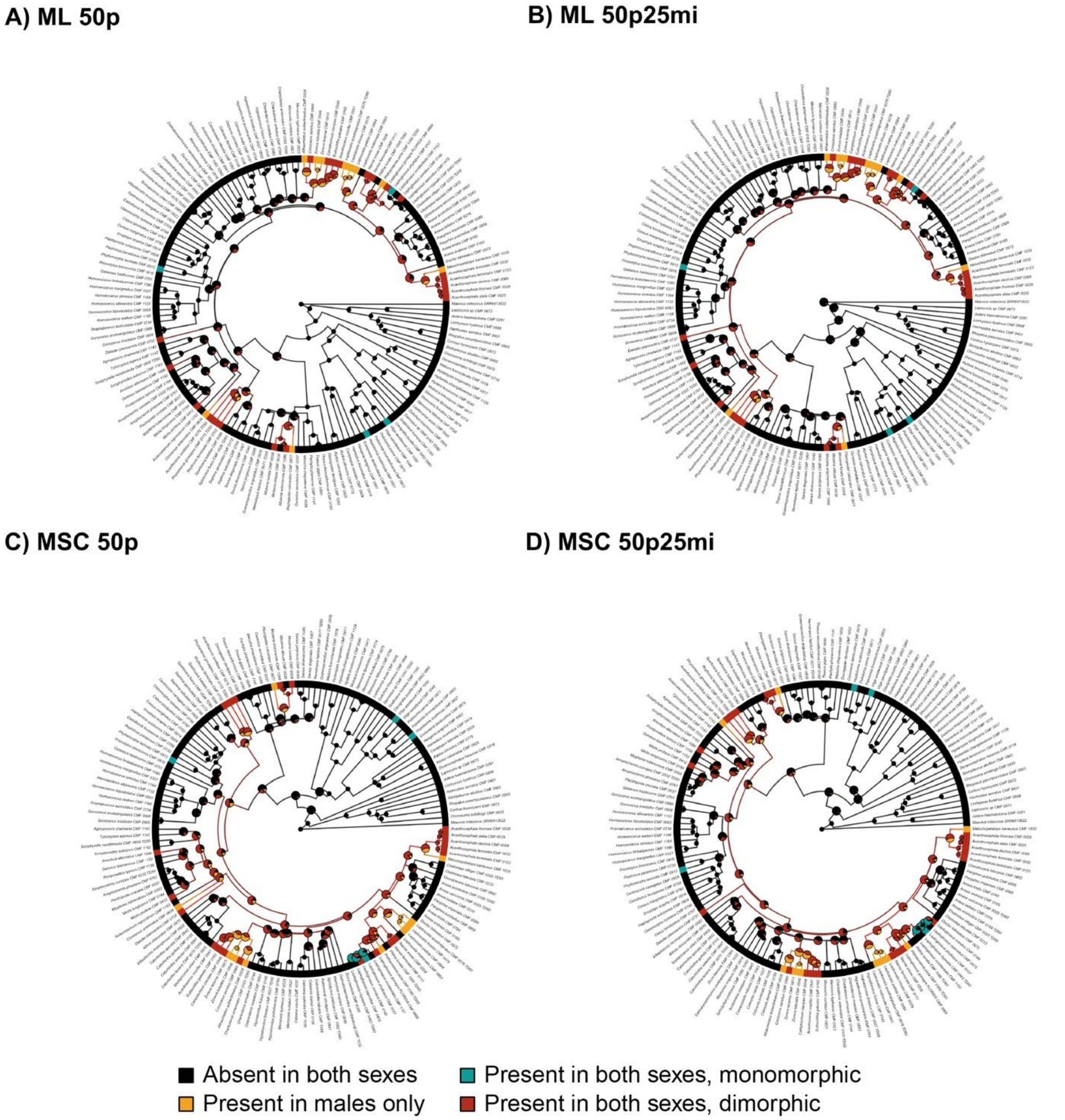


**Figure S8.** Stochastic character mapping of the lateral femoral processes on the (A) ML 50p, (B) ML 50p25mi, (C) MSC 50p, and (D) MSC 50p25mi phylogenies. The terminal bars show the character state assigned to each species. Branch colors summarize model-averaged stochastic maps, illustrating inferred histories of transitions among character states. Pie charts at internal nodes represent posterior probabilities of each character state, with the radius reflecting node certainty: nodes with < 0.90 posterior support for the predominant state are larger. Outgroup taxa are omitted for visualization.

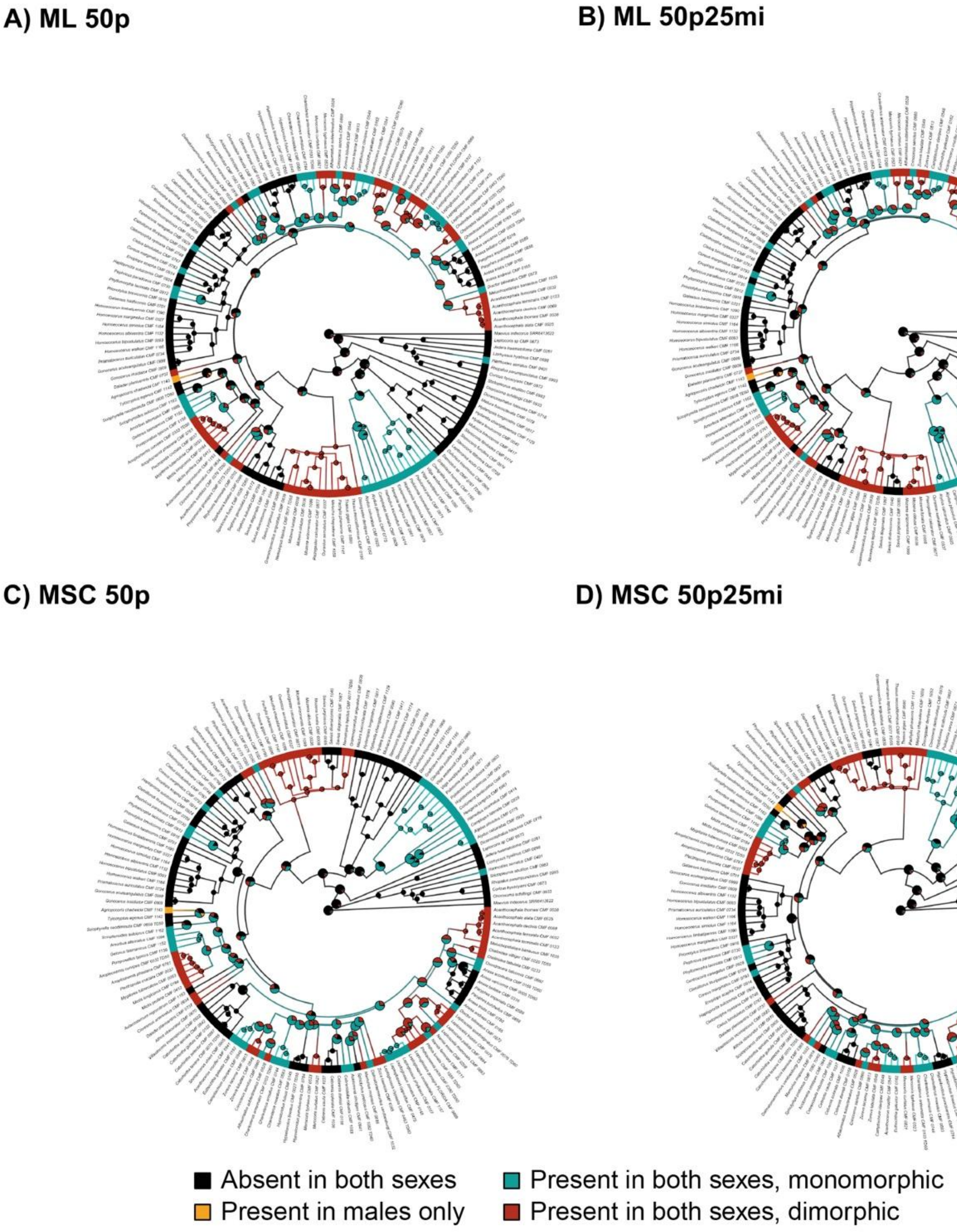


**Figure S9.** Stochastic character mapping of the ventral femoral processes on the (A) ML 50p, (B) ML 50p25mi, (C) MSC 50p, and (D) MSC 50p25mi phylogenies. The terminal bars show the character state assigned to each species. Branch colors summarize model-averaged stochastic maps, illustrating inferred histories of transitions among character states. Pie charts at internal nodes represent posterior probabilities of each character state, with the radius reflecting node certainty: nodes with < 0.90 posterior support for the predominant state are larger. Outgroup taxa are omitted for visualization.

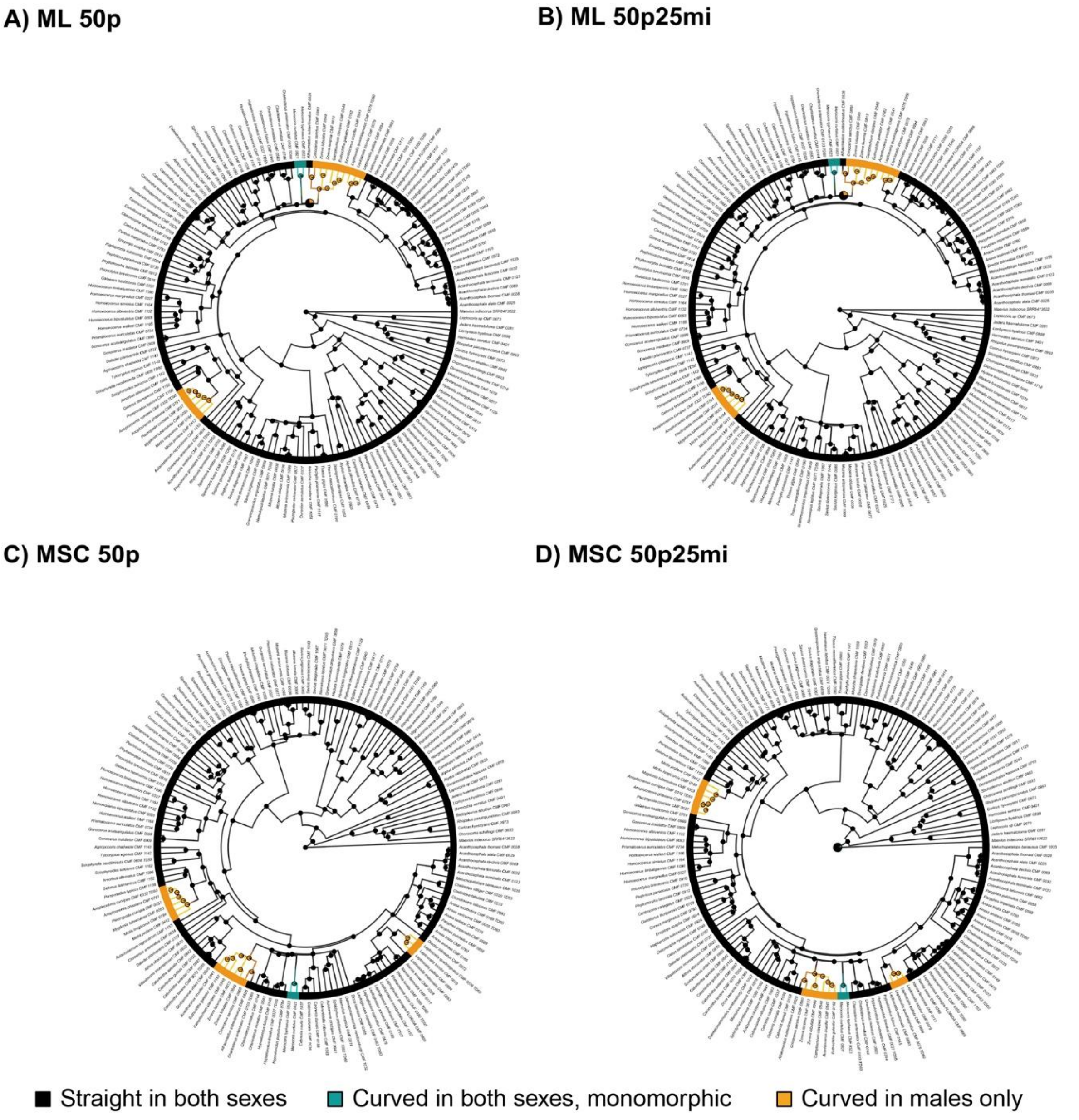


**Figure S10.** Stochastic character mapping of the femoral curvature on the (A) ML 50p, (B) ML 50p25mi, (C) MSC 50p, and (D) MSC 50p25mi phylogenies. The terminal bars show the character state assigned to each species. Branch colors summarize model-averaged stochastic maps, illustrating inferred histories of transitions among character states. Pie charts at internal nodes represent posterior probabilities of each character state, with the radius reflecting node certainty: nodes with < 0.90 posterior support for the predominant state are larger. Outgroup taxa are omitted for visualization.

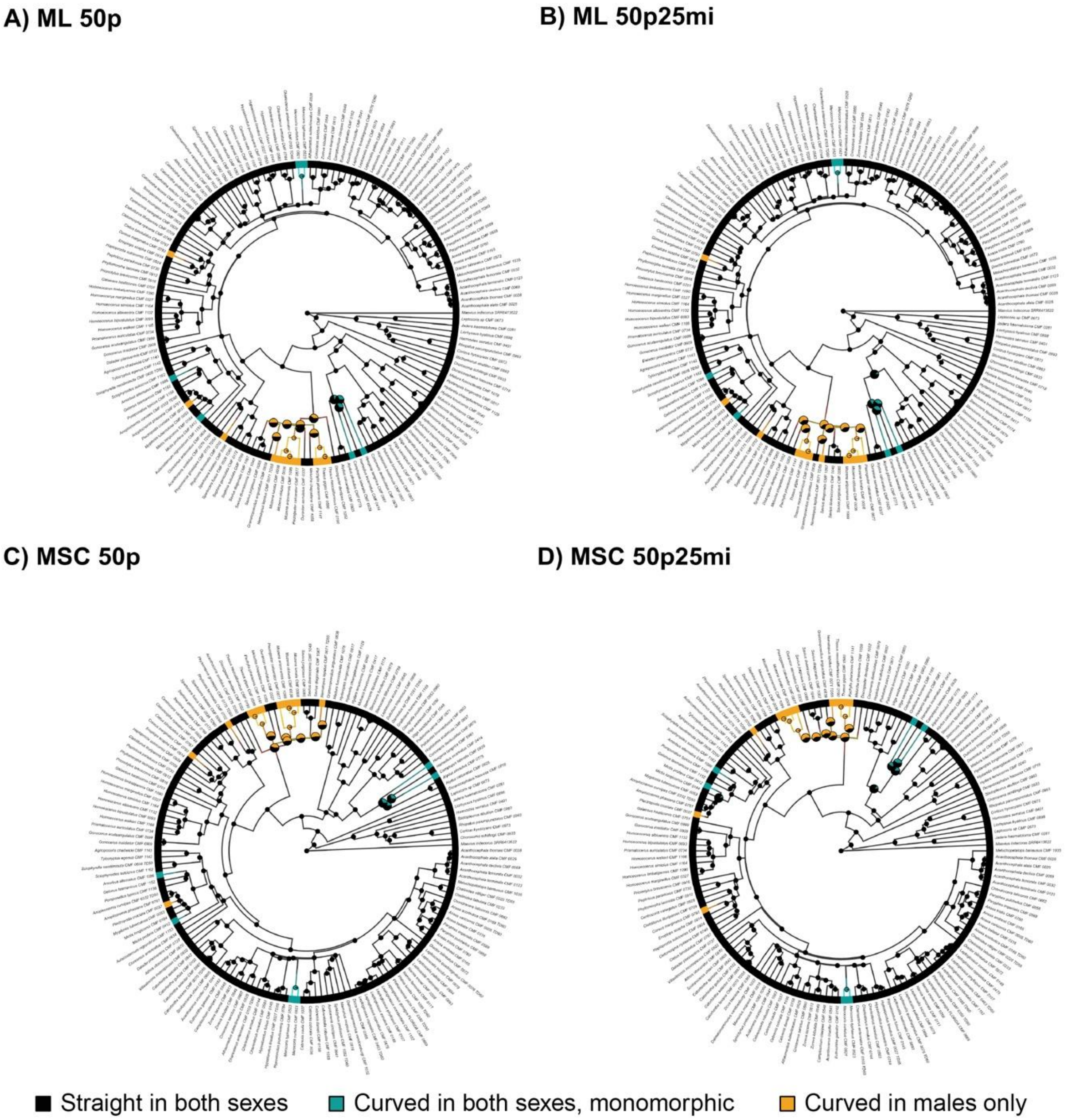


**Figure S11.** Stochastic character mapping of the tibial curvature on the (A) ML 50p, (B) ML 50p25mi, (C) MSC 50p, and (D) MSC 50p25mi phylogenies. The terminal bars show the character state assigned to each species. Branch colors summarize model-averaged stochastic maps, illustrating inferred histories of transitions among character states. Pie charts at internal nodes represent posterior probabilities of each character state, with the radius reflecting node certainty: nodes with < 0.90 posterior support for the predominant state are larger. Outgroup taxa are omitted for visualization.

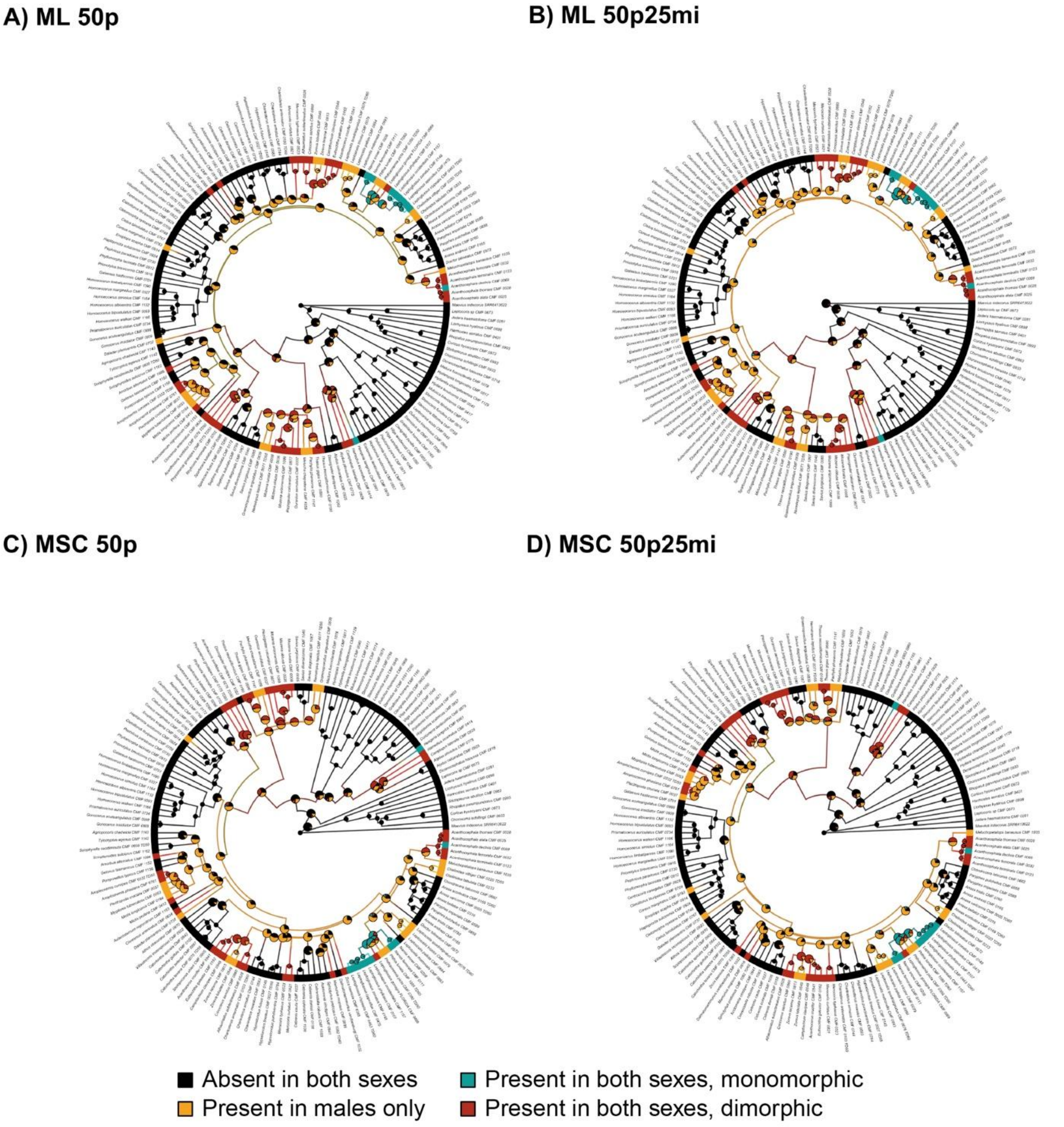


**Figure S12.** Stochastic character mapping of the internal tibial processes on the (A) ML 50p, (B) ML 50p25mi, (C) MSC 50p, and (D) MSC 50p25mi phylogenies. The terminal bars show the character state assigned to each species. Branch colors summarize model-averaged stochastic maps, illustrating inferred histories of transitions among character states. Pie charts at internal nodes represent posterior probabilities of each character state, with the radius reflecting node certainty: nodes with < 0.90 posterior support for the predominant state are larger. Outgroup taxa are omitted for visualization.

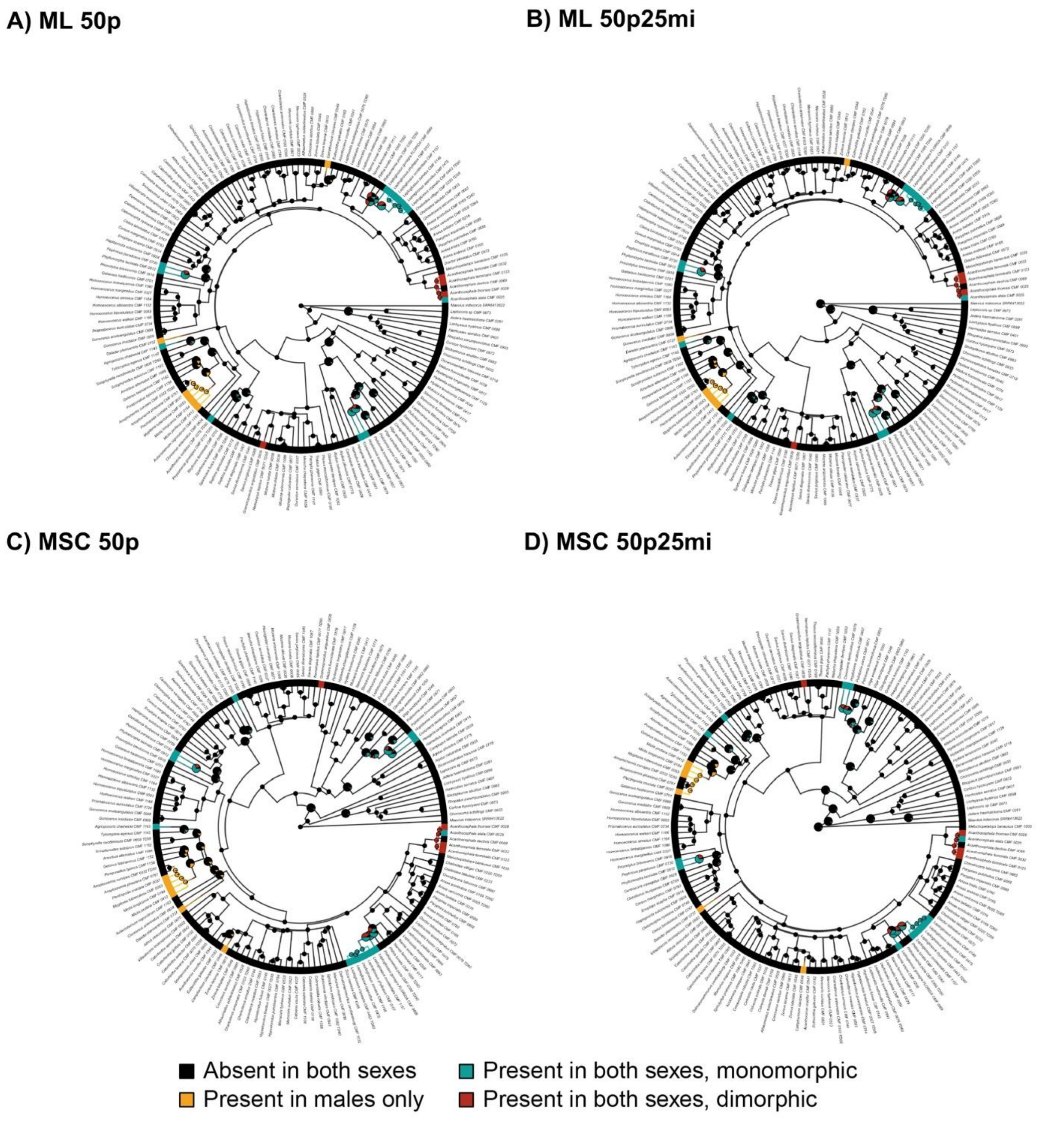


**Figure S13.** Stochastic character mapping of the external tibial processes on the (A) ML 50p, (B) ML 50p25mi, (C) MSC 50p, and (D) MSC 50p25mi phylogenies. The terminal bars show the character state assigned to each species. Branch colors summarize model-averaged stochastic maps, illustrating inferred histories of transitions among character states. Pie charts at internal nodes represent posterior probabilities of each character state, with the radius reflecting node certainty: nodes with < 0.90 posterior support for the predominant state are larger. Outgroup taxa are omitted for visualization.

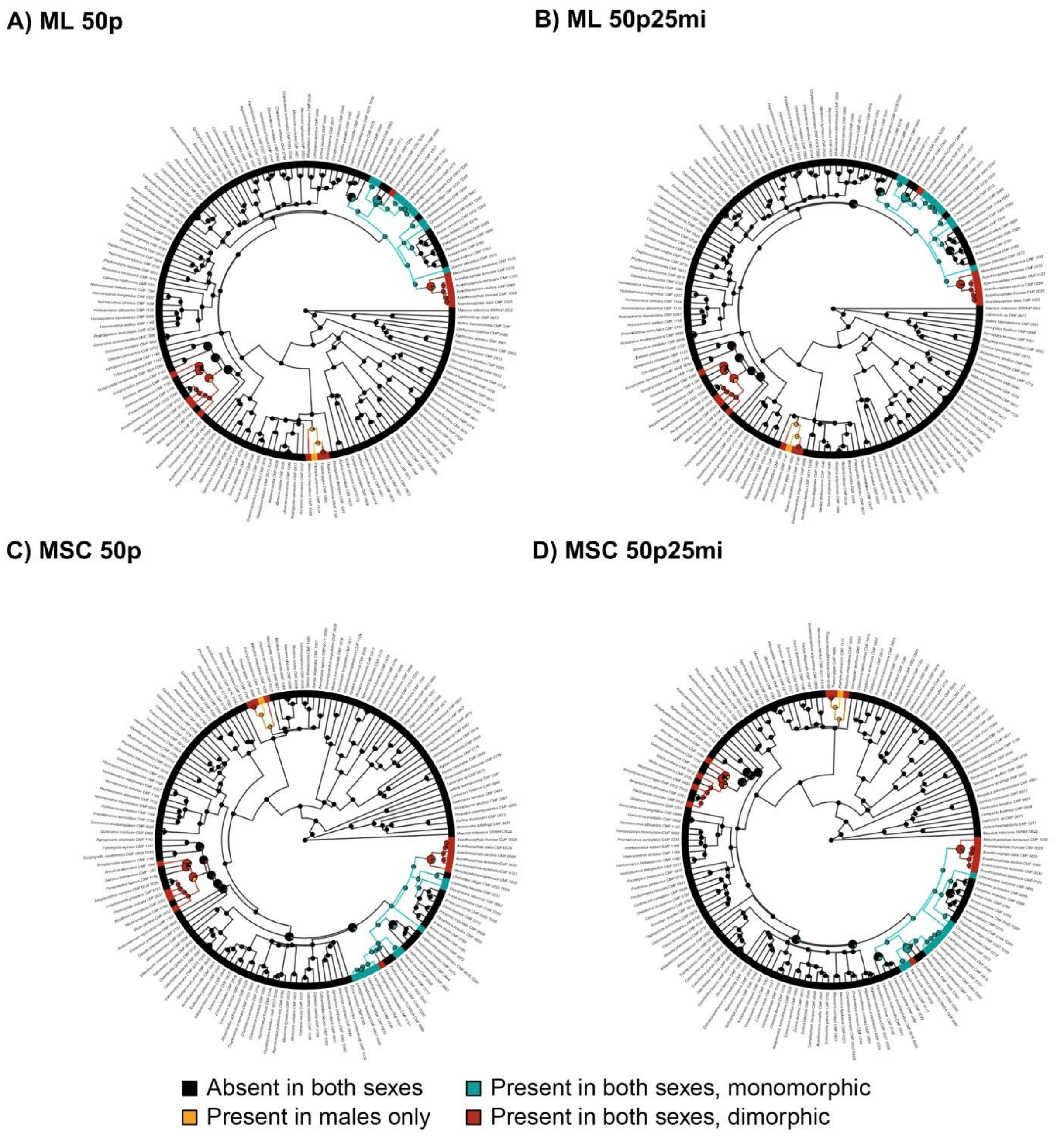


**Figure S14.** Stochastic character mapping of the internal tibial flag on the (A) ML 50p, (B) ML 50p25mi, (C) MSC 50p, and (D) MSC 50p25mi phylogenies. The terminal bars show the character state assigned to each species. Branch colors summarize model-averaged stochastic maps, illustrating inferred histories of transitions among character states. Pie charts at internal nodes represent posterior probabilities of each character state, with the radius reflecting node certainty: nodes with < 0.90 posterior support for the predominant state are larger. Outgroup taxa are omitted for visualization.

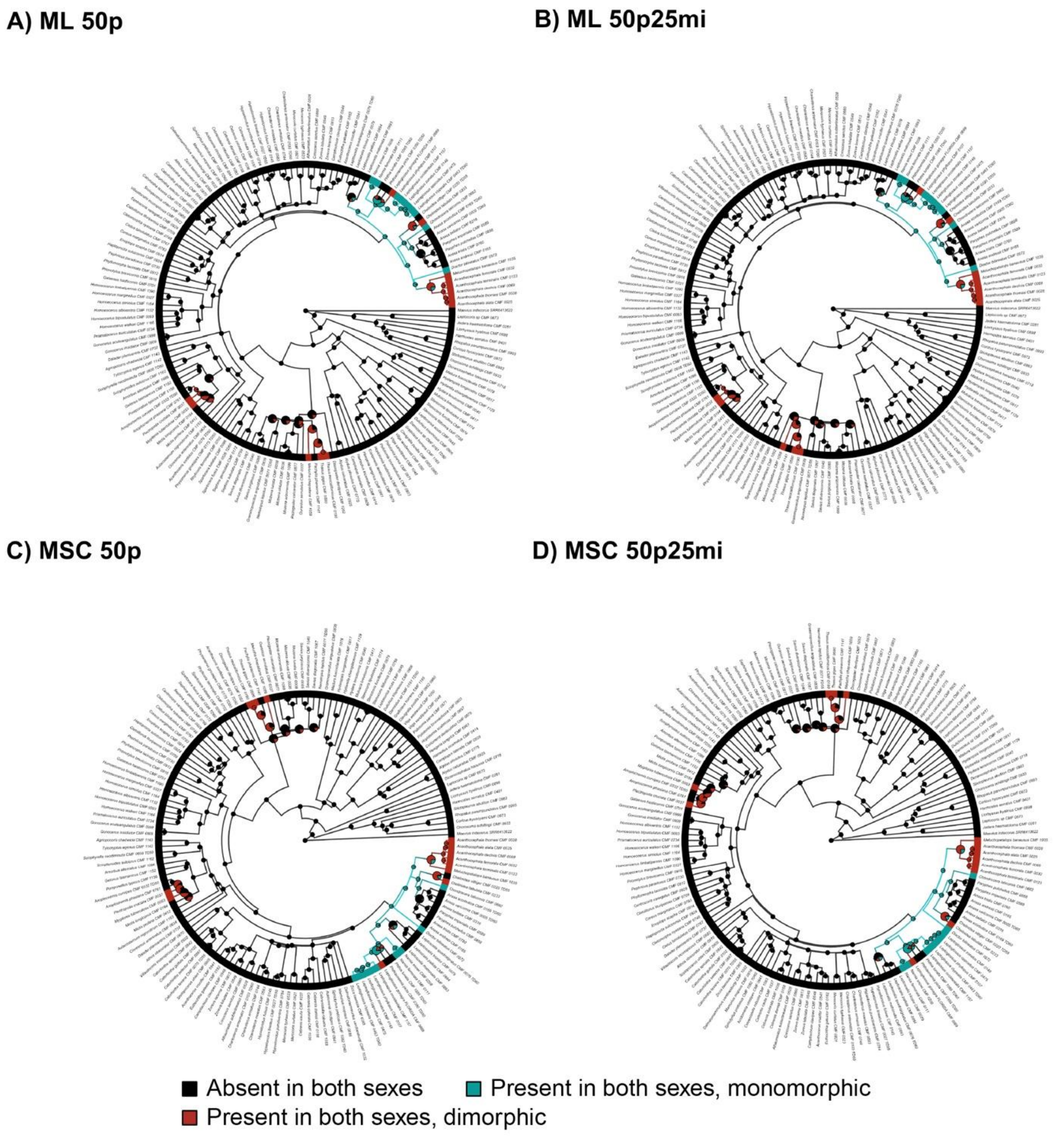


**Figure S15.** Stochastic character mapping of the external tibial flag on the (A) ML 50p, (B) ML 50p25mi, (C) MSC 50p, and (D) MSC 50p25mi phylogenies. The terminal bars show the character state assigned to each species. Branch colors summarize model-averaged stochastic maps, illustrating inferred histories of transitions among character states. Pie charts at internal nodes represent posterior probabilities of each character state, with the radius reflecting node certainty: nodes with < 0.90 posterior support for the predominant state are larger. Outgroup taxa are omitted for visualization.

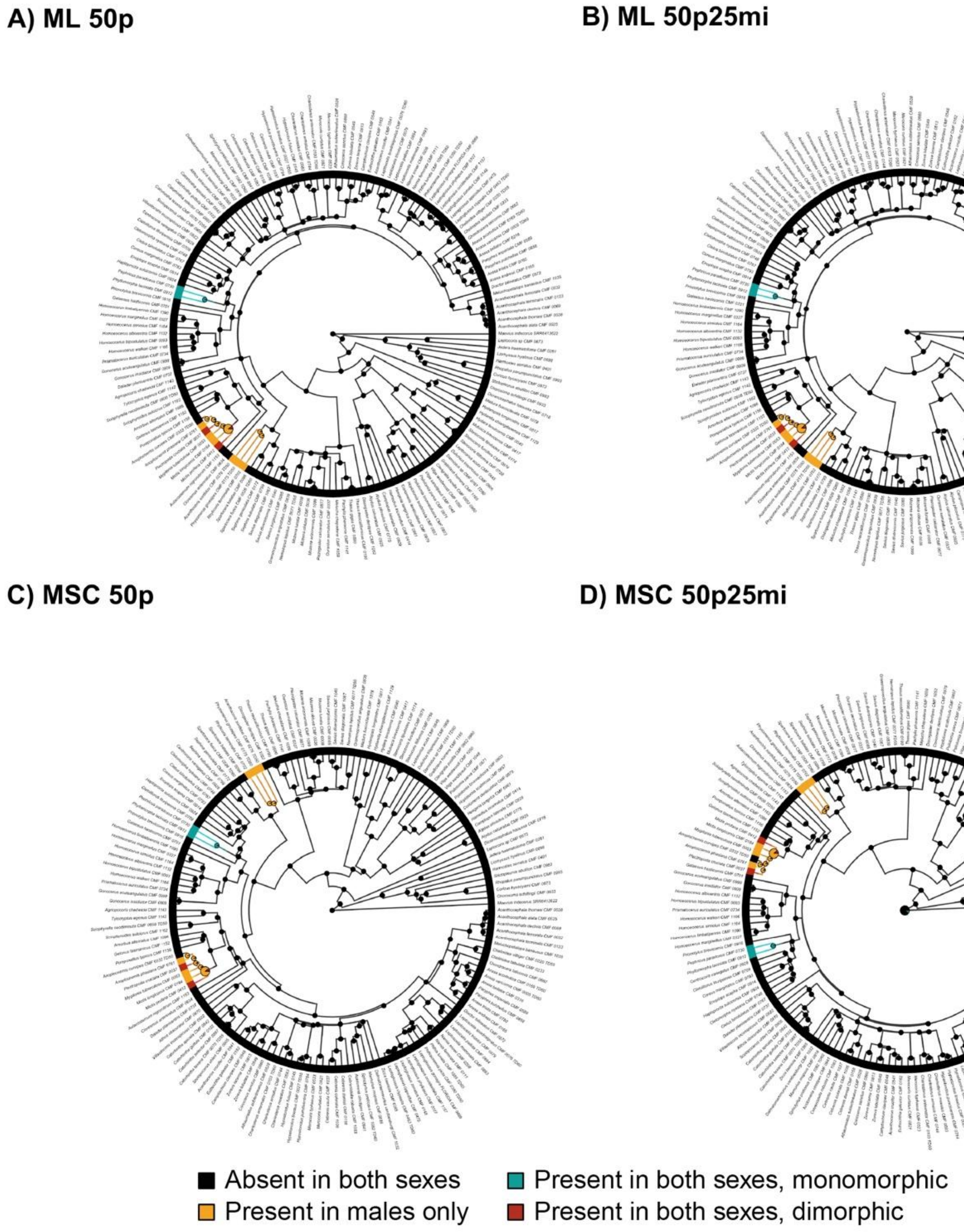


**Figure S16.** Stochastic character mapping of the ventral abdominal processes on the (A) ML 50p, (B) ML 50p25mi, (C) MSC 50p, and (D) MSC 50p25mi phylogenies. The terminal bars show the character state assigned to each species. Branch colors summarize model-averaged stochastic maps, illustrating inferred histories of transitions among character states. Pie charts at internal nodes represent posterior probabilities of each character state, with the radius reflecting node certainty: nodes with < 0.90 posterior support for the predominant state are larger. Outgroup taxa are omitted for visualization.

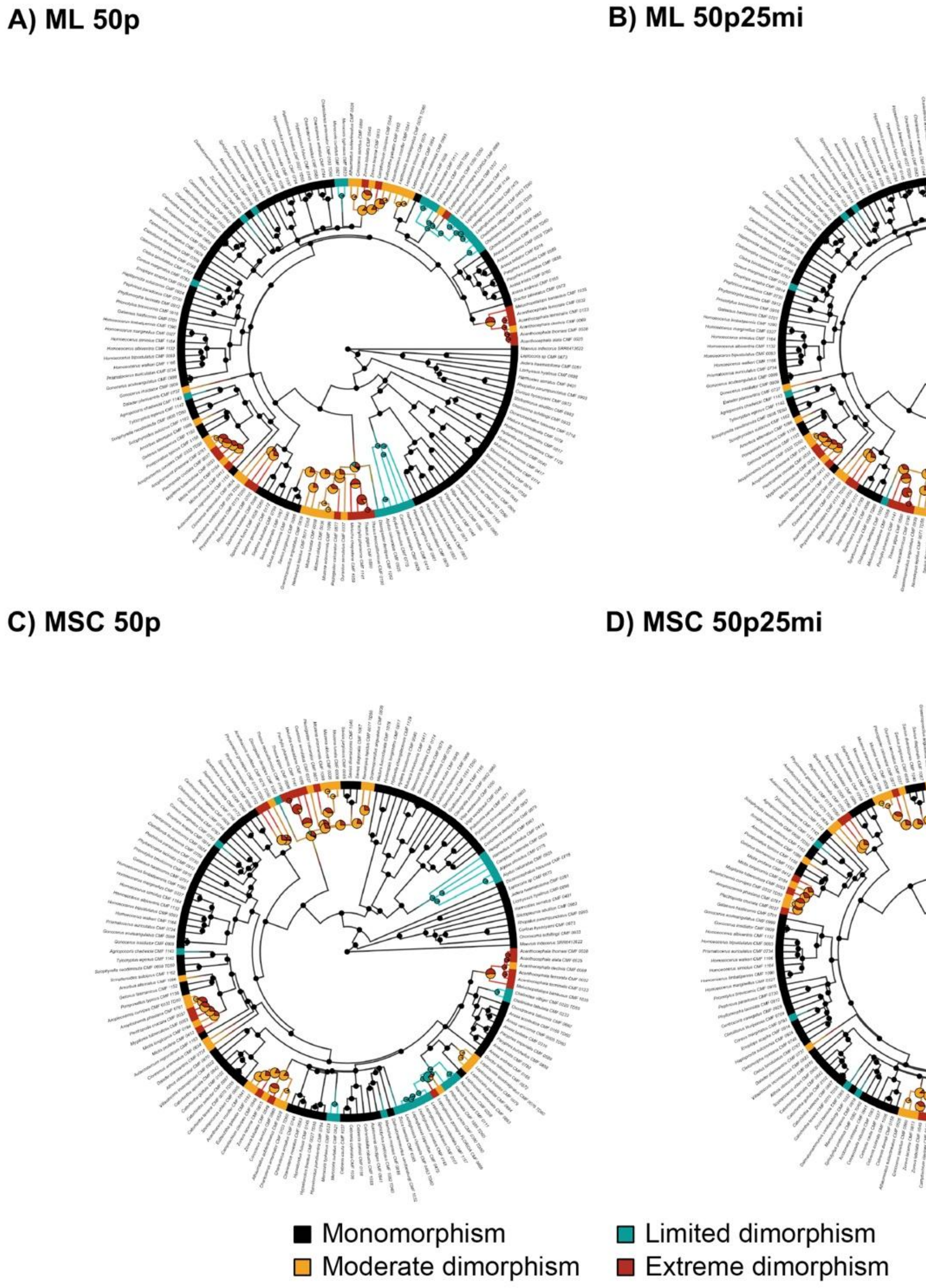


**Figure S17.** Stochastic character mapping of the overall degree of sexual dimorphism on the (A) ML 50p, (B) ML 50p25mi, (C) MSC 50p, and (D) MSC 50p25mi phylogenies. The terminal bars show the character state assigned to each species. Branch colors summarize model-averaged stochastic maps, illustrating inferred histories of transitions among character states. Pie charts at internal nodes represent posterior probabilities of each character state, with the radius reflecting node certainty: nodes with < 0.90 posterior support for the predominant state are larger. Outgroup taxa are omitted for visualization.
